# Characterization of recombinase-based genetic parts and circuits using nanopore sequencing

**DOI:** 10.1101/2025.05.27.656450

**Authors:** F. Veronica Greco, Sarah K. Cameron, Shivang Hina-Nilesh Joshi, Sarah Guiziou, Jennifer A. N. Brophy, Claire S. Grierson, Thomas E. Gorochowski

**Affiliations:** School of Biological Sciences, University of Bristol, 24 Tyndall Avenue, Bristol, BS8 1TQ, UK; Earlham Institute, Norwich Research Park, Norwich, Norfolk, NR4 7UZ, UK; Department of Bioengineering, Stanford University, Shriram Center 203, Stanford, CA 94305, USA

**Keywords:** recombinases, characterization, genetic circuits, biometrology, synthetic biology

## Abstract

Recombinases are versatile enzymes able to perform the precise insertion, deletion, and rearrangement of DNA and can act as a foundation for programmable genetic logic and memory. Fundamental to their use are accurate measurements of function. However, these are often laborious, time-consuming, and costly to collect. To address this, we developed a semi-automated workflow that combines low-cost liquid handling robotics, multiplexed long-read nanopore sequencing, and a supporting computational analysis tool to enable the high-throughput and detailed characterization of recombinase parts and circuits when used in a variety of contexts and organisms. Our approach overcomes the limitations of typically used fluorescence-based assays and is able to monitor temporal dynamics, observe structural changes at a nucleotide resolution, and unravel the internal workings of complex multi-state circuits. The ability to scale-up and automate genetic circuit characterization is an essential step towards more rigorous biological metrology that can support the construction of predictive models for efficiently engineering biology.

## INTRODUCTION

The ability to reprogram cellular behaviors through the engineering of biology has the potential to address global challenges spanning sustainability to healthcare [1]. Synthetic genetic circuits offer a way of tapping into the computational capabilities of cells, rewiring and extending their functionalities in new ways [2]. To date, most genetic circuits have been built using protein-based transcription factor networks. These offer a flexible platform for regulating the flows of transcription within a cell and allow for small regulatory motifs encoding basic computational operations (e.g., logic gates) to be easily combined to create larger circuits implementing more complex computations [3]. While these circuits have been used to implement diverse information processing in a wide range of organisms [3–10], they require precise and continuous expression of circuit components which can be affected by stochastic molecular fluctuations and circuit context (genetic, host and environmental) [11], and imparts a continuous burden on the cell [12]. This can lead to circuits whose functions are not always reliable and which are difficult to scale in complexity [3, 13–15].

In contrast, recombinase-based circuits encode their state directly within DNA, using enzymes (e.g., integrases) to cause modifications through the addition, removal, and rearrangement of DNA sequences [10, 16, 17]. This allows for the implementation of robust computational operations that are discrete by nature (i.e., a DNA sequence is either present or absent and has a specific orientation). Recombinase-based circuits have been used to create robust genetic logic [6–10, 18–21] and memory arrays that encode ‘bits’ of information in the orientation of short independent segments of DNA [16]. Such circuits hold much promise, but their development is hampered by a limited number of well-characterized recombinase enzymes [22] and difficulties debugging large non-functional circuits. This stems from the fact that typical monitoring tools like fluorescent proteins are only able to simultaneously capture a handful of internal circuit states [23], making it impossible to understand how a large multi-state circuit is actually functioning. Furthermore, expression of fluorescent reporter proteins often requires changes to be made to a circuit. These changes can themselves alter function and introduces delays in measurements, complicating the characterization of circuit dynamics. New experimental methods are needed that enable the full state of these systems to be monitored more directly over time. Long-read sequencing could hold the answer, but its high cost and typically low-throughput nature has so far limited its use in synthetic biology [24].

Here, we address this issue by developing a semi-automated workflow to enable the high-throughput and detailed characterization of large recombinase-based genetic circuits. We show how low-cost liquid handling robotics can be combined with nanopore sequencing and computational analysis tools to monitor the complete state of recombinase circuits over time. We demonstrate our workflow’s capabilities by simultaneously characterizing of the kinetics and off-target activity of six recombinase enzymes, monitor the dynamics of a multi-plasmid recombinase-based genetic cascade, and show how spatial activation of recombinase switches can be quantified across plant roots. The ability for our workflow to process up to 96 samples in 2 days using a single Opentrons OT-2 robot ensures that it is accessibility to most labs, supporting the roll-out of more rigorous metrology for engineering biology.

## RESULTS

### A workflow for characterizing recombinase-based parts and circuits

Our workflow consists of four steps that cover DNA extraction, target amplification, sequencing, and analysis (**Figure 1**; **Supplementary Notes 1 and 2**). DNA extraction depends specifically on the application and type of organism, but can be performed using standard methods like heat, chemical or mechanical lysis. After DNA extraction is complete, we then provide automated liquid handling protocols for Opentrons OT-2 robots based on a standard 96-well plate format, which can also be performed manually if required (**Supplementary Note 3**). The workflow starts by amplifying the recombinase circuit via a “tailing” PCR. This PCR uses primers designed to target the start and end of the recombinase circuit and append common DNA sequences to both ends of the amplicons to aid later barcoding. Once these PCR reactions are complete, they are separately purified using magnetic beads and the DNA concentration for each sample quantified. These DNA quantities are fed back into the workflow to allow for automated normalization of concentrations before a further barcoding PCR step is performed. This PCR makes use of primers that target the common tails added to each amplicon and further appends a unique barcode sequences to allow for distinguishing the source sample after sequencing. These sample specific PCR reactions are then separately purified using magnetic beads and their DNA quantified before pooling in equimolar amounts. This single pooled sample is then sequenced after a standard ligation-based nanopore sequencing library preparation.

**Figure 1:**
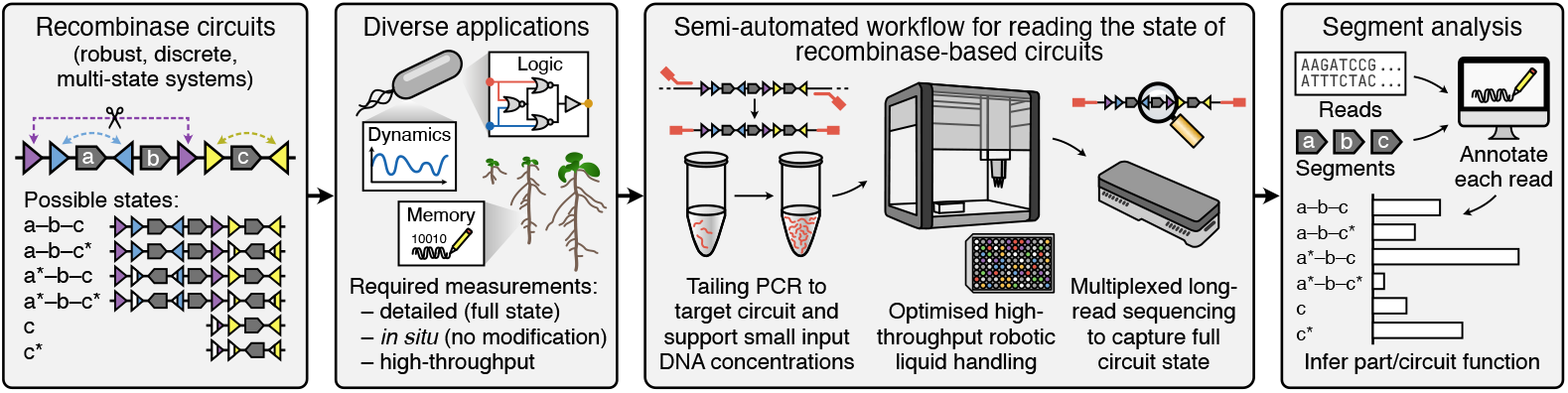
Overview of the characterization workflow. (left–right) Recombinase circuits enable the creation of robust multi-state systems for biological computation and memory. Our workflow allows for small DNA samples to be taken and uses PCR to “tail” a target region containing the circuit for analysis via long-read nanopore sequencing. The tailing PCR and subsequent sequencing library preparation can be carried out in high-throughput using optimized protocols on a low-cost liquid handling robot (Opentrons OT-2). These libraries are then nanopore sequenced to provide full-length information about circuit state. Finally, analysis of the sequencing data is performed using segment analysis, where user provided segments (DNA sequences) that can be used to capture the full state of the circuit are searched for within each read and for all samples processed. The ‘segment’ computational tool automates this step outputting the presence, order, and orientation of each provided segment within each read. Distributions of the states present in a sample can then be used to characterize circuit function and dynamics for samples taken over time.

During development of the experimental protocols, several optimizations were made to reduce sequencing costs and handling time, while maintaining data quality and quantity. Some of the key finders were that: (i) sample volumes as small as 25 µL from bacterial liquid cultures provided enough template material for amplification, (ii) that the use of a heat-lysis step after flash freezing in liquid nitrogen was sufficient to extract DNA for both bacteria and dissected plant tissues, and (iii) that magnetic bead purification could support size selection of target amplicons such that non-ligated sequencing adapters were efficiently removed (**Supplementary Figure 1**), improving overall sequencing throughput.

Once sequencing is complete, analysis of the sequencing data is carried out using our computational tool called ‘segment’ (**Supplementary Figure 2**; **Methods**). Because the state-space of a recombinase circuit grows exponentially as new recombination events become possible (i.e., the addition of a new set of recombinase sites can potentially triple the size of the state space through the absence or presence of a new region of DNA in either a forward or reverse direction), we do not align reads to all possible reference sequences. Instead, the segment tool takes as input short “segment sequences” in a multi-FASTA format. Segment sequences need to be provided for the start and end of the circuit, as well as any regions within the circuit that undergo change (e.g., addition/removal or a change in orientation) as the circuit functions. By focusing on the segments that change rather than every possible state of the full circuit, much less information needs to be provided by a user. Furthermore, the information given allows for every read to be fully classified by searching for the start and end segments to orientate the sequence, and then searching for all other possible segments to generate a full classification that is encoded as a concatenation of the segment names in the order they are found from the start to end segment, with the orientation of each segment being denoted by the absence or presence of an asterisk appended to the segment name (e.g., if segment S1 is found in a reverse orientation then it is denoted by S1* in the read classification). Using this approach, all reads can be classified and a summary of observed states and their frequency generated. From these summaries, it is then possible to assess overall circuit function (**Figure 1**).

**Figure 2:**
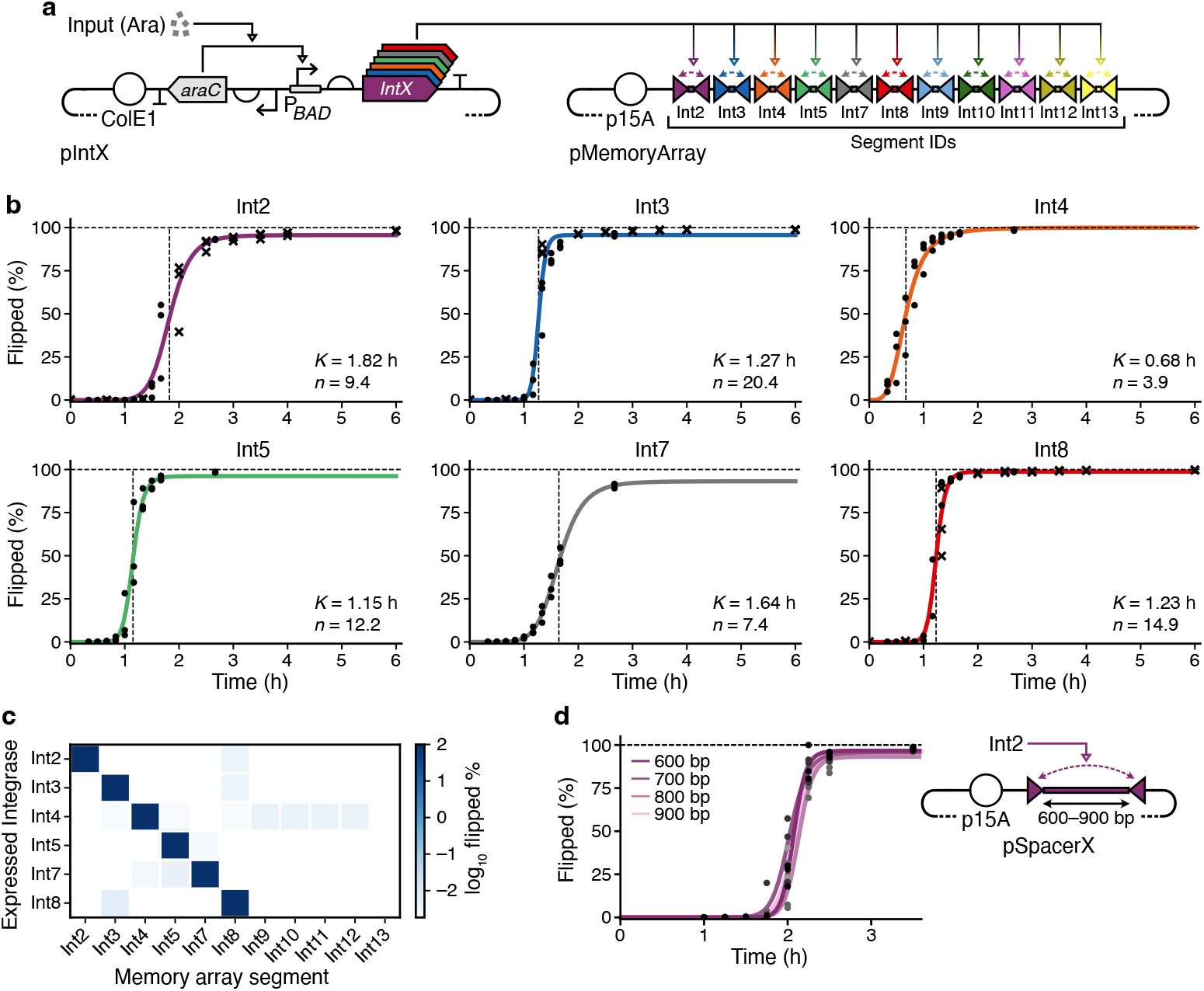
Detailed characterization of six recombinase enzymes. (**a**) Schematic of the integrase expression plasmids (pIntX, left) and the memory array (pMemoryArray, right) that contains 11 different integrase recognition site pairs (*attB* and *attP*) flanking short DNA sequences such that cognate integrase expression causes a non-reservable inversion (flip) of that specific segment of DNA, permanently recording an event. (**b**) Characterization of the temporal activity of 6 integrase enzymes. Solid lines show a fitted Hill function with *K* and *n* parameters shown in the bottom right corner of each plot. *K* value is also denoted by a vertical dashed line. Black circles and crosses are individual measurements from the biological replicates for samples taken at differing time points: 0, 40, 80, 120, 150, 180, 210, 240, and 360 min, and 20, 30, 40, 50, 60, 70, 80, 90, 100, and 160 min, respectively. (**c**) Orthogonality matrix for the 6 integrases tested and 11 integrase recognition sites. Off-target activity denoted by the flipping of non-cognate recognition sites. (**d**) Testing the effect of different lengths of DNA (600–900 bp) between recognition sites of the Int2 integrase on the temporal flipping of the target site. Schematic of the pSpacerX plasmids built for this experiment shown right of the plot.

### Detailed characterization of a recombinase library

To demonstrate the core features of our workflow, we performed a detailed characterization of six large-serine recombinases (Int2, Int3, Int4, Int5, Int7 and Int8) in *Escherichia coli* that were previously mined from sequence databases [16, 22]. First, we assessed the dynamic response of the recombinases by monitoring the inversion (flipping) of their cognate recognition sites over time after recombinase expression was induced. We made use of a previously assembled 11-bit memory array that consisted of 11 short DNA sequences where each is flanked by a specific pair of recombinase recognition sites in orientations that would cause an inversion in the DNA sequence between when the cognate recombinase is present (**Figure 2a**). The orientation of these 11 short sequences can capture up to 2^11^ = 2048 different states.

Cells were co-transformed with plasmids allowing for induction of a specific recombinase and containing the 11-bit memory array. Experiments were then preformed where recombianse expression was induced (at *t* = 0 hours) and then samples taken over a 6 hour period for processing by our workflow (**Figure 2b**, black filled circles). To analyze the state of the memory array, we provided sequences for each of the flanked regions as segments (labeled ‘IntX’, where ‘X’ is the recombinase of interest) and calculated the percentage that had flipped by comparing the IntX (unflipped) to IntX* (flipped) states for each recombinase in each read using the segment tool. After this initial experiment, we found that for some of the recombinases the complete response curve was not well captured. We therefore performed an additional with samples taken over a shorter 2 hour 40 min period for a subset of the recombinases (**Figure 2b**, black crosses). Combining data from both these experiments, we were able to recover complete response curves for all six recombinases. Even though the response of many of the recombinases was fast, taking less than 30 min to transition to 100% flipped (e.g., for Int3 and Int8), we saw fairly similar responses across measurements from biological replicates, demonstrating a good reproducibility in the method. Hill function fits to the data for strong expression of the recombinases showed a large diversity in response dynamics, with the time to reach 50% of cells having flipped DNA (*K* values) ranging from approximately 40 to 110 min and cooperativities (*n* values) spanning from 3.9 to 20.4 (**Figure 2b**; **Supplementary Table 3**).

For multiple recombinases to be used concurrently within a circuit it is essential that they display minimal off-target activities. Typically, when using fluorescence reporters to monitor recombination events, only a small number of alternative sites can be simultaneously tested (typically only a few at a time) due to the spectral overlap of the reporter proteins. Our methodology is not limited by this constraint. Because we use an array of recombinase recognition sites that is read in its entirety, we are able to simultaneously measure off-target activity of all 10 recombinase sites simultaneously. This substantially simplifies characterization of off-target effects and could easily be scaled to much larger numbers of potential sites as long-read sequencing has been shown able to produce reads more than 100 kb in length.

Further analysis of the sequencing data from the memory array showed that all recombinases precisely targeted their cognate sites, with only minor off-target activity for Int4 across numerous recognition sites (Int9–Int12), Int2–Int4 weakly recognizing Int8 sites, and some cross-reactivity between Int5 and Int7, and Int2 and Int8, respectively. In all cases, off-target activity was typically very small with *<*0.1% of regions flipped after 3 hours (**Figure 2c**).

Finally, we carried out a complementary experiment for Int2 to assess whether the length of DNA flanked by recombinase sites affected the efficiency and dynamics of flipping. For lengths from 600–900 bp (typical for a protein coding gene), we saw no substantial difference (**Figure 2d**; **Supplementary Table 3**). However, when comparing the rates of flipping (i.e., steepness of the response function) of 600–900 bp segments to the short 50 bp segment of the memory array, there was notable decrease with *n* values shifting from ∼21 to 9.4 for the short segment. This likely stems from physical constraints in the accessibility and bending of DNA that are required for efficient recombination. It also suggests that design of circuits using recombinases may need to consider the lengths of flanked DNA regions to ensure a desired function is achieved.

### Observing multi-state dynamics of recombinase-based circuits

A major advantage of using long-read sequencing to monitor recombinase-based circuits is that more complex multi-state changes can be directly quantified in full. To demonstrate this capability, we made use of a multi-component genetic cascade in which a set of three recombinases trigger the sequential expression of each other through the structural rearrangement (flipping) of subsequent recombinase protein coding sequences (**Figure 3a**) [16]. A further complication of this system is that not all components are present on a single plasmid but instead split across two, offering the chance to assess our methodology when targeting multiple loci within a single genetic system.

**Figure 3:**
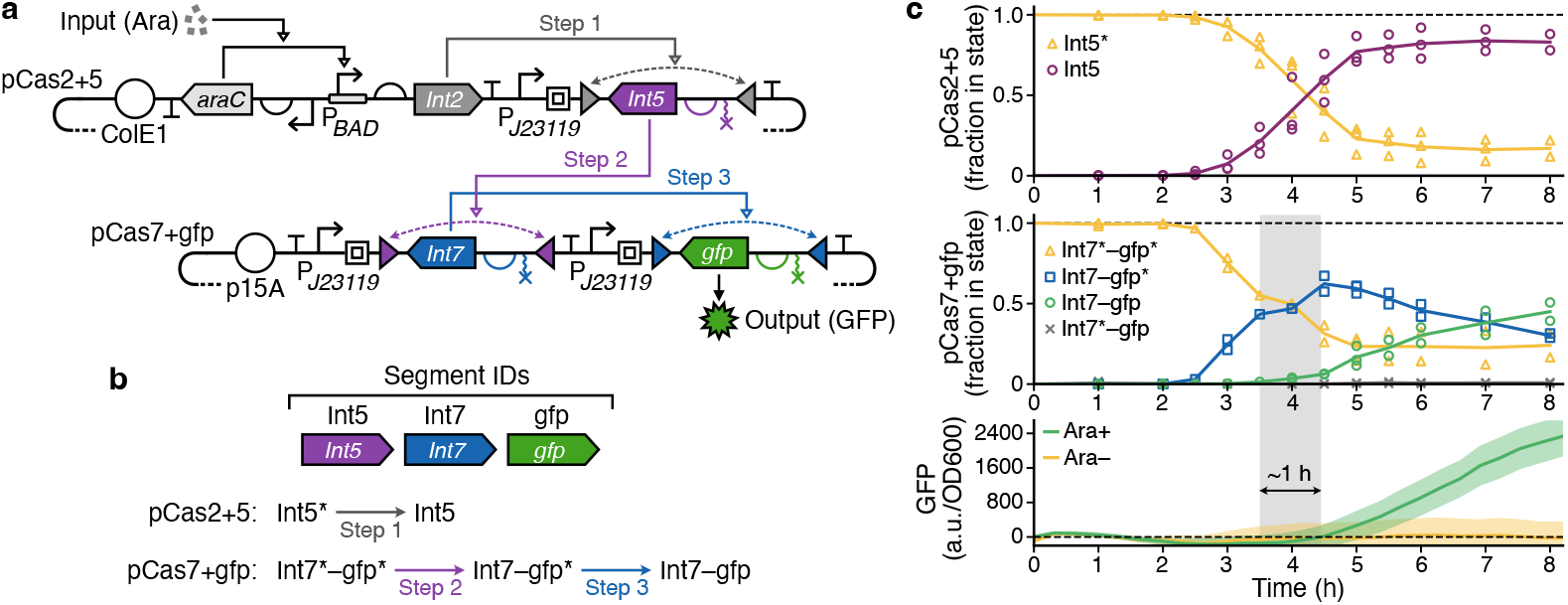
Monitoring of a multi-step recombinase-based cascade. (**a**) Schematic of the recombinase-based genetic cascade. The circuits spans two plasmids (pCas2+5 and pCas7+gfp) and requires the flipping of coding sequences of two recombinase enzymes (Int5 and Int 7) and an output green fluorescence protein (GFP). The cascade is triggered by the presence of arabinose (Ara). (**b**) Segments used for the analysis of circuit state. Transition steps between states for each of the plasmids are shown. (**c**) Top two plots show the fraction of reads for the pCas2+5 and pCas7+gfp plasmids in the different possible states over time. Solid lines show the average fraction and unfilled triangles, circles, squares and crosses show the individual measurements taken for all three of the biological replicates. Bottom plot shows GFP output normalized to OD600 over time when the cascade is induced with arabinose (Ara+, green solid line) or not (Ara–, yellow solid). The standard deviation of the three biological replicates is shown by the colored shaded regions. Grey shaded period denotes the delay in the flipping of the *gfp* coding sequence in the pCas7+gfp plasmid measured by sequencing, and the subsequent change in fluorescence after the time taken to express the protein and for it to mature (∼1 h).

We performed experiments in *E. coli* where the genetic cascade was triggered (at *t* = 0 hours) and then the state of the system monitored through sampling of the culture over time. Samples were processed using our automated workflow with two separate PCRs to target the two relevant regions making up the full circuit (present on the pCas2+5 and pCas7+gfp plasmids, respectively). To quantify circuit state, we used the two recombinase coding sequences that were initially flipped (Int5 and Int7) and the output green fluorescent protein (GFP) as segments to search for within each read (**Figure 3b**). This allowed us to quantify the proportion of cells at each step in the cascade.

Analysis showed a clear propagation of inversions in the coding sequences of the recombinases and GFP over time that corresponded to the sequential expression of each recombinase in the cascade. Similar to the characterization experiments of the recombinases in isolation, we saw increased rates in sequence inversion by Int5 compared to Int7. The ability to take very small samples from a growing culture for monitoring allowed us to perform these experiments in a plate reader and monitor GFP fluorescence throughout the experiment. Integrating this data with the dynamic changes in the complete genetic state via sequencing, we were able to see a clear delay of ∼1 hour between changes in the orientation of the *gfp* gene and the subsequent fluorescence produced after gene expression is activated. This highlights a key benefit that more direct sequencing-based measurements of circuit state have over more commonly used fluorescent reporter proteins.

### Quantifying the spatial activation of recombinase-based switches in plants

A major benefit of recombinase-based circuits is their ability to function robustly across diverse organisms [8, 25, 26]. A particularly promising avenue of their use is tracking cellular decision making during the development of more complex multicellular organisms and the creation of robust switches that can permanently turn on functionalities in specific cell types [27]. Currently, monitoring the state of these types of circuit is challenging. Fluorescence microscopy is typically used [27], however, this limits the number of states that can be simultaneously read and hampers accurate quantification of circuit state across large tissues, as cell boundaries are often difficult and time consuming to delineate.

To demonstrate the ability for our workflow to overcome these challenges, we analyzed the spatial activation of a recombinase-based switch that had been integrated into the genome of *Arabidopsis thaliana* (**Figure 4a**). The circuit was designed to strongly activate GFP expression in lateral roots. This was achieved by using the P_GATA23_ promoter (active in lateral root primordia [28]) to trigger expression of the PhiC31 recombinase, which then permanently flips the orientation a constitutive promoter to drive expression of GFP in primordia-derived cells (i.e., lateral roots).

**Figure 4:**
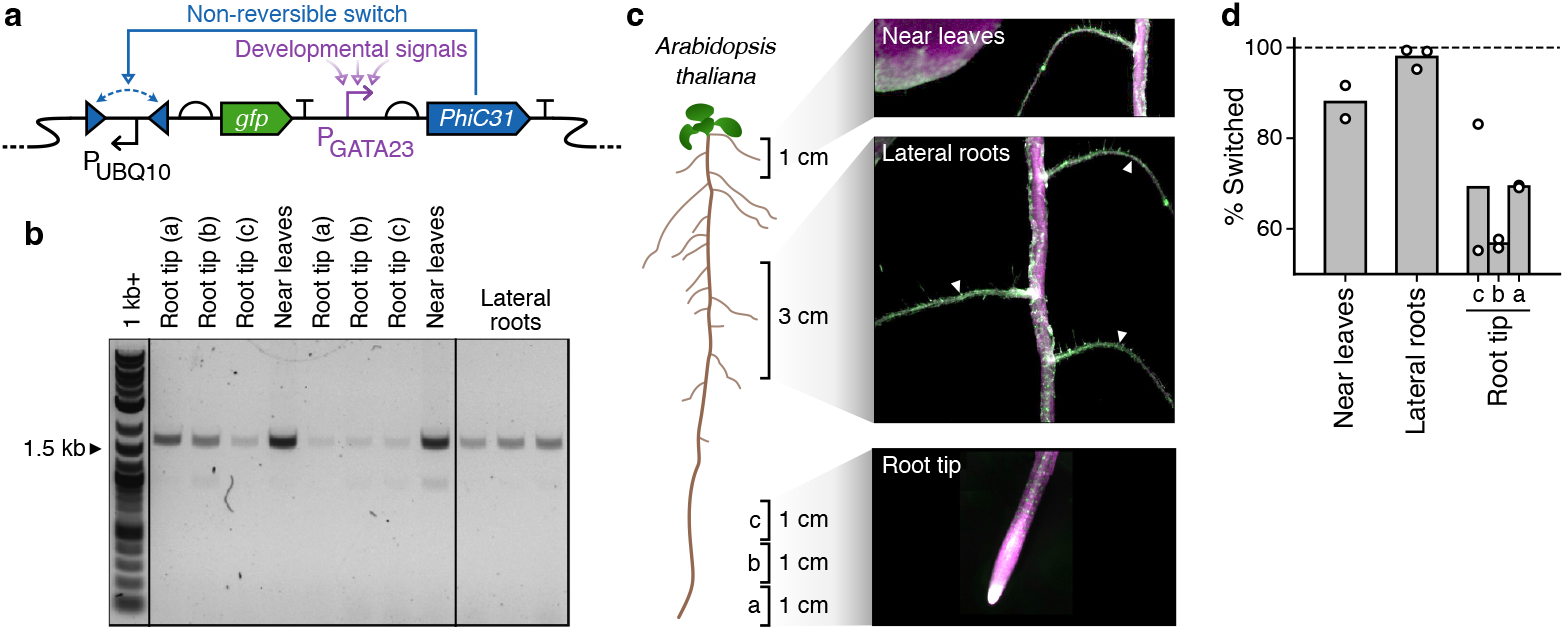
Monitoring spatial activation of a recombinase-based switch in plant roots. (**a**) Schematic of the recombinase-based genetic switch. Expression of the PhiC31 recombinase by the P_GATA23_ promoter (active in lateral roots) causes the non-reversible flipping of the P_UBQ10_ promoter, which is then able to drive green fluorescent protein (GFP) expression. (**b**) Tailed DNA from different sections of root after magnetic bead clean-up. Roots were cut into sections illustrated in panel c and then placed directly in tailing PCR reactions. Magnetic bead clean-ups were carried out on the PCR products. Amplicon expected to have an ∼1.5 kb length. (**c**) Sections of *Arabidopsis thaliana* roots that were dissected for analysis. White triangles denote the lateral roots that were separated from the primary root. (**d**) Activation of the genetic switch in different sections of the root. Values for biological replicates of the experiment shown by white filled circles.

To assess activation of the circuit throughout the root system, we dissected three key areas: (i) a 1 cm section of the primary root near the leaves, (ii) lateral roots in a 3 cm section in the middle of the root system, and (iii) the root tip into three separate 1 cm sections (**Figure 4c**; **Methods**). A significant issue we faced when working with this dissected material was the very small amounts of DNA that could be recovered. The use of PCR in our workflow enabled us to target and amplify the specific regions of interest. However, optimization of the conditions and polymerases used in this reaction was required before amplicons were consistently recovered for the tailing PCR step (**Figure 4b**; **Methods**).

As expected, analysis of the circuit showed near full activation (98%) in lateral roots (**Figure 4d**). Lowest activation was seen in the root tip (56–69%) and slightly higher activation in the primary root near the leaves (88%). This qualitatively matched the clear expression of GFP throughout lateral roots when imaged by microscopy (**Figure 4c**). We also found fairly consistent values across biological replicates of the experiment. The quantitative difference we observed in activity in the primary root near the leaves and the root tip is likely due to low basal expression from the P_GATA23_ promoter. Because cells near the leaves have lived for longer than those in the root tip, there is a higher chance of permanent activation in the cells near the leaves due to stochastic gene expression events (e.g., transcriptional bursts) over time.

## DISCUSSION

Recombinase-based genetic circuits offer much promise for the robust reprogramming of diverse organisms [25]. However, for this to become a reality, many researchers have highlighted the need for efficient and comprehensive approaches for reading and interpreting the state of these circuits [22, 29, 30]. Here, we have addressed this need, showing how a semi-automated experimental and computational workflow can enable high-throughput monitoring of entire recombinase-based circuits with minimal requirements in terms of starting DNA material and no requirements to make changes to the circuits themselves. Furthermore, the ability for nanopore sequencing to be carried out in the field at point of use [31], opens up new avenues for monitoring these circuits no matter where they are deployed.

To date, over 4,000 putative large-serine recombinases have been identified, but the vast majority of these remain uncharacterized [16, 25, 32]. A better understanding of recombinase diversity would significantly expand the toolkit of parts available for synthetic biologists and offer a more systematic understanding of their function across the tree of life. Our workflow is ideally suited to address this need. It allows for the sampling of time-course experiments to recover reaction kinetics and the use of large arrays of diverse recombinase recognition sites can characterize numerous off-target activities, simultaneously. Combining our sequencing approaches with high-throughput chip-based DNA synthesis, would enable an even more comprehensive understanding of how these molecular machines work and ideal data sets to train AI models for future *de novo* design of these molecular machines.

While our methodology offers insights into recombinase part and circuit function, there are several remaining challenges. First, the use of automation for preparing the sequencing libraries helps to greatly reduce costs, but these are still not insignificant and greater multiplexing of samples would be beneficial. The use of acoustic liquid handling systems to reduce sample volumes further to nanoliter-scales could potentially support this requirement. Second, our use of PCR to both target and amplify a recombinase circuit could potentially introduce bias in some cases. This would be especially problematic for circuits in which very large insertions or deletions of genetic material occurs, as these cause vastly different sequence compositions and lengths that may bias amplification. The application of alternative approaches to DNA targeting based on enrichment using biotinylated DNA probes [33] or Cas9 cleavage [34], plus the incorporation of barcoded sequencing adapters through direct ligation or the use of transposases could help overcome these issues and offers further elaborations on our workflow for specific use cases.

In summary, recombinase-based genetic circuitry offers bioengineers the rare ability to create reliable genetic logic and memory due to its direct encoding of state within the structural organization of a circuit’s own DNA. Scaling our ability to build and characterize such systems will be critical to their use in applications spanning agriculture to health [1]. This work provides the integrated experimental and computational tools needed to take steps towards this goal, helping broaden the metrology available for understanding large and complex genetic circuitry [13, 35], and ultimately supporting improved robustness in the engineered biology we create.

## METHODS

### Strains, media and chemicals

Cloning and characterization experiments in bacteria were performed using *E. coli* strain DH10-beta (Δ(ara–leu) 7697 araD139 fhuA ΔlacX74 galK16 galE15 e14– *ϕ* 80dlacZΔM15 recA1 relA1 endA1 nupG rpsL (StrR) rph spoT1 Δ(mrr-hsdRMS-mcrBC) (C3019I, New England Biolabs). *E. coli* were grown in NEB 10-beta stable outgrowth medium (B9035S, New England Biolabs) for transformation, and LB broth (L3522, Sigma-Aldrich) for general propagation and characterization. Antibiotic selection was performed using 50 µg/ml Chloramphenicol (C0378-25G, Sigma-Aldrich) and 50 µg/mL kanamycin (K1637, Sigma-Aldrich). Induction of circuit components for the recombinase characterization and cascade experiments was performed using 10 nM of L-Arabinose (1B1473-25G, VWR), and for samples where no induction was present, cells were grown in LB media supplemented with 0.5% D-glucose (G7528, Sigma-Aldrich) to ensure strong catabolite repression of the P_*BAD*_ promoter.

Transgenic plants were generated by *Agrobacterium tumefaciens*-mediated floral dip of *Arabidopsis thaliana* Columbia ecotype [36]. Transgenic seeds contain a constitutively expressed mCherry marker *proUBQ10*::*mCherry* that was used to isolate transgenic seeds via fluorescence microscopy. A single insertion homozygous transgenic line was isolated and used for switch analysis *in planta*.

### Plant growth conditions

Transgenic *Arabidopsis thaliana* plants were grown in Percival-Scientific growth chambers (CU41L4) at 22°C in 16/6-hour light/dark cycles with 60% humidity. Light was provided by fluorescent bulbs at a total intensity of ∼100 microEinsteins (µE). Plants for imaging and switch analysis were plated as seeds onto sterile 0.7% gelzan (Sigma G19010) media containing 1X Murashige and Skoog nutrients (Caisson MSP01-50LT) and 1% sucrose (Sigma S0389) at pH 5.7. Seeds were surface sterilized prior to plating with 95% ethanol and 20% bleach. Plates were sealed with micropore tape (3M 1530-0) for gas exchange and placed vertically into the growth chamber.

### Assembly of pSpacerX plasmids

The pSpacerX plasmids were assembled in a 2-step cloning process. First, a 4-part Golden-Gate reaction was used to create a template plasmid with a 1 kb spacer flanked by Int2-attP and Int2-attB sites. This reaction was composed of a destination backbone (p15A-CamR), two short fragments with the Int2-attP and Int2-attB sequences designed to flank the spacer, and a 1 kb inert spacer sequence derived from the LacI protein coding sequence. This plasmid was then used as a template for full-plasmid PCR and blunt-end ligation using a series of oligos that would truncate the LacI sequence, resulting in the spacers of the desired lengths. For the Golden-Gate reaction, PCR fragments were generated from pMemoryArray for the plasmid backbone, as well as the Int2-attP and Int2-attB recognition sequences, and the 1 kb LacI spacer was amplified from the *E. coli* genome. pMemoryArray was a gift from Christopher Voigt (Addgene plasmid #60585). The NEBridge Golden Gate Assembly Kit (BsaI-HF v2) (E1601, New England Biolabs) was used for the Golden-Gate assembly reaction using standard conditions, except reactions volumes were scaled to a quarter of the normal volume. For the subsequent blunt-end ligation cloning, a KLD Enzyme Mix (M0554, New England Biolabs) was used following the standard manufacturer protocol. For all assembly reaction transformations, 2 µL of the assembly reactions were transformed to 12.5 µL of chemically competent DH10-beta cells (C3019, New England Biolabs). All plasmid sequences were verified by Sanger sequencing (Eurofins Genomics).

### Recombinase characterization experiments

For experiments performing a detailed characterization of a set of recombinases (**Figure 2**), *E. coli* DH10-beta competent cells were transformed with both a plasmid expressing a recombinase (pIntegrase 2–8; #60574–60579, Addgene) and an 11-bit Memory Array (pMemoryArray; #60585, Addgene). For the experiments with varying segment lengths (pSpacerX plasmids), *E. coli* DH10-beta competent cells were transformed with both the pIntegrase2 plasmid and a specific pSpacerX plasmid with the appropriate segment length. Single colonies of transformed cells were then inoculated in 5 mL of LB media supplemented with kanamycin and chloramphenicol and 0.5% glucose to suppress expression of the P_BAD_ promoter. Cultures were grown overnight for 15 hours in a shaker incubator (SI500, Stuart) at 37°C and 250 rpm. After the overnight growth, measurements of optical density at 600 nm (OD600) were taken using a WPA, Biowave II spectrophotometer to assess cell growth. For induction, pre-cultures were diluted by the appropriate factor to an OD600 of 0.3 in 3 mL of media supplemented with kanamycin and chloramphenicol, and 0.5% glucose to maintain minimal expression for uninduced samples. These experiments were conducted in 24-deep well plates (V6831, Promega UK) at 37°C and 1250 rpm in a shaker incubator (SI505, Stuart). Recombinase expression was induced using 10 mM L-arabinose, after 1 hour of initial incubation. For the initial characterization experiments, 25 µL samples were collected and flash-frozen at 0, 40, 80, 120, 150, 180, 210, 240, and 360 minutes after induction. For the second characterization experiment, 25 µL samples were collected and flash-frozen at 20, 30, 40, 50, 60, 70, 80, 90, 100, and 160 min after induction. For the experiments with varying segment lengths, the same protocol was followed, however, 25 µL samples were collected and flash-frozen at 70, 80, 90, 100, 110, 120, 130, 140, 150, and 210 min after induction with 10 mM L-arabinose. All samples were immediately stored at –80°C before processing using the automated workflow (**Supplementary Note 1**) using tailing primers specific for the plasmids present (**Supplementary Table 4**).

### Experiments with a genetic cascade

For experiments studying the genetic cascade (**Figure 3**), *E. coli* DH10-beta competent cells were transformed with two plasmids encoding a three-integrase cascade (pCas2+5 and pCas7+gfp; #60594–60595, Addgene). Single colonies of transformed cells were then inoculated in 5 mL of LB media supplemented with kanamycin and chloramphenicol and 0.5% glucose to suppress expression of the P_BAD_ promoter. Cultures were grown overnight for 15 hours in a shaker incubator (SI500, Stuart) at 37°C and 250 rpm. After the overnight growth, measurements of optical density at 600 nm (OD600) were taken using a WPA, Biowave II spectrophotometer to assess cell growth. For induction, pre-cultures were diluted by the appropriate factor to an OD600 of 0.3 in 3 mL of media supplemented with kanamycin and chloramphenicol, and 0.5% glucose to maintain minimal expression for uninduced samples. These experiments were conducted in 24-deep well plates (V6831, Promega UK) at 37°C and 1250 rpm in a shaker incubator (SI505, Stuart). Recombinase expression was induced using 10 mM L-arabinose, after 1 hour of initial incubation and 25 µL samples were collected and flash-frozen at 0, 60, 120, 150, 180, 210, 240, 270, 300, 330, 360, 420 and 480 minutes after induction. All samples were immediately stored at –80°C before processing using the automated workflow (**Supplementary Note 1**) using tailing primers specific for the plasmids present (**Supplementary Table 4**).

### Experiments monitoring the spatial activation of recombinase switches in plants

For experiments monitoring the spatial activation of genetic switches in plant roots (**Figure 4**), we made use of *A. thaliana* plants that contained a stably integrated recombinase switch for monitoring lateral root development (**Figure 4a**). This strain had been generated via floral dip, with the first generation of seeds (T1) having been collected from selective media supplemented with antibiotic. These seeds were used to generate a second generation of transformant seeds (T2) that acted as the basis of our experiment. T2 *A. thaliana* seeds were first sterilized using ethanol to avoid bacterial and fungal contamination. Around 50 seeds were poured into a 1.5 mL microfuge tube and 1 mL of 70% ethanol was pipetted into the tube. The microfuge tube containing the seeds was incubated for 20 min on a tube rotator at room temperature. The ethanol was then removed by pipetting and particular care was taken to ensure that no residues were left. Following this, 1 mL of 95% ethanol was used to carry out a final wash of the seeds. After adding the ethanol, the seeds were incubated for 5 min on a tube rotator at room temperature. The tube containing the sterilized seeds was then transferred to a bio-safety cabinet. Ethanol was then removed by pipetting and 1 mL ultrapure water was added to the seeds to facilitate plating. Plating was done immediately after adding the water to avoid the water imbibing the seeds or causing them to stick together. Seeds were sown onto sterile plates containing 0.7% Gelzan (G19010, Sigma), 1X Murashige and Skoog nutrients (MSP01-50LT, Caisson) and 1% sucrose (S0389, Sigma) at pH 5.7. The pH was adjusted using KOH. Seeds were dispensed 1.5 cm apart to facilitate visualization and cutting of the seedling roots. Gas exchange was facilitated by sealing the plates using a breathable micropore tape (1530-0, 3M) and incubated in the dark for 72 hours at 4°C, making sure to maintain a vertical inclination of the plates all the time. Plates were subsequently transferred to a culture growth chamber and grown for 10 days at 22°C with cycles of 16 hours of light and 8 hours of dark, with 60% humidity levels. Light was provided by LED arrays at a total intensity of ∼100 µE.

For microscopy, 10 days after sowing, the plants were imaged using a Leica Thunder Model Organism widefield fluorescence microscope. Excitation of fluorophores was performed using 450/40 nm excitation and 525/50 nm emission filters and 546/11 nm excitation and 605/70 nm emission filters for GFP and mCherry, respectively.

After testing several methods based on chemical and physical DNA extraction, we found that direct use of dissected material in a 50 µL tailing PCR reaction using the KAPA Taq HotStart PCR Kit (KK1512, Kapa Biosystems) performed well. This approach was used for all experiments with plant roots.

### Automated experimental workflow

Automated protocols were developed for the Opentrons OT-2 (Opentrons Inc.) liquid handling robot to carry out cell lysis and multiplexed sequencing library preparation (**Supplementary Note 1**). These protocols allow for the processing of 96 samples in parallel and generate two separately barcoded sequencing libraries. We found that running more than 48 samples per flowcell was possible, but that variation in run-to-run throughput (typically caused by the quality of the input DNA) could result in some samples having insufficient reads to accurately assess circuit state. The total cost of running 96 samples through the automated experimental workflow was approximately £5.89 per sample (excluding the costs of the sequencing kits and flow cells) with a total runtime on the OT-2 of 13 hours and 39 minutes (excluding robot setup and calibration time). Detailed information about each automated protocol can be found in **Supplementary Note 1** and the optimization of these protocols is described in **Supplementary Note 2**. All OT-2 protocols were written using Python version 3.9.6 and the Opentrons API version 2.11. For lower-throughput requirements, we also provide descriptions in **Supplementary Note 3** of manual protocols to carry out the experimental workflow.

In all of our experiments, heat lysis of the cells was conducted using the Temperature Module with PCR Aluminium Block (GEN2, Opentrons) and all PCR reactions were completed using the on-deck Thermocycler (GEN1, Opentrons). Both DNA purification steps were completed using the Magnetic Module (GEN2, Opentrons) with KAPA pure magnetic beads (KK8002, KAPA Biosystems) at 0.4:1 beads to sample ratio. Opentrons GEN2 electronic pipettes were used during the pipeline (Single channels: P20, P300, P1000. Multi-channels: P20, P300). **Supplementary Table 1** summarizes the steps involved in the automated pipeline, with protocol script numbers, modules and pipettes that were used.

All DNA quantification for the automated workflow was performed using the Quantifluor ONE dsDNA system (E4870, Promega) in two 96-well plates. Standards were pre-prepared using the OT-2 robot to the standard curve suggested by the manufacturer. Specifically, 1 µL of each sample was added to 200 µL of Quantifluor Dye. This was mixed and incubated for 5 minutes at room temperature and protected from the light before being read using a SpectraMax plate reader (ID5, Molecular Devices) with 485 nm excitation and 535 nm emission filters. Analysis was performed using the Quantifluor Dye Systems Data Analysis Excel Workbook provided by Promega. Sequence-specific tailing primers were synthesized by IDT (**Supplementary Table 4**). Tailed samples were barcoded using a PCR Barcoding Expansion 1–96 (EXP-PBC096, Oxford Nanopore Technologies) to allow the pooling and running together of 48 sample sequencing libraries. Final nanopore sequencing libraries were generated using the standard ligation sequencing kit (SQK-LSK109, Oxford Nanopore Technologies).

### Nanopore sequencing

DNA sequencing libraries were sequenced using R9.4.1 flow cells (FLO-MIN106D, Oxford Nanopore Technologies) on a MinION Mk1B device (Oxford Nanopore Technologies) controlled using MinKNOW version 21.11.7. The resulting FAST5 sequencing data was basecalled and demultiplexed using guppy version 6.5.7 using the ‘dna r9.4.1 450bps hac.cfg’ high-accuracy model and ‘EXP-PBC096’ barcode kit to generate sample specific FASTQ files that could then be used as input for downstream analyses.

### Computational analysis

Analysis of sequencing data was performed using segment version 1.0.0. Additional computational analyses and all plots were produced using custom Python scripts. These were run using Python version 3.12.9 with matplotlib version 3.10.0, pandas version 2.2.3, numpy version 1.26.4 and scipy version 1.14.1.

### Data Availability

Python scripts for all OT-2 protocols and custom labware definitions for the automated elements of the workflow are included as **Supplementary Data 1**. A description of all the pre-requisite reagents/equipment plus a summary of all steps involved are provided in **Supplementary Note 1**. Code for the segment tool can be found at: https://github.com/BiocomputeLab/segment.

## ACKNOWLEDGMENTS

F.V.G. was supported by a Royal Society PhD Studentship. T.E.G. and C.S.G. were supported by BrisEng-Bio, a UKRI-funded Engineering Biology Research Centre grant BB/W013959/1. T.E.G. was additionally supported by the UKRI Engineering Biology Mission Award CYBER under BBSRC grant BB/Y007638/1, a Turing Fellowship from The Alan Turing Institute under EPSRC grant EP/N510129/1, and a Royal Society University Research Fellowship grant URF*\*R*\*221008. J.A.N.B. is supported by U.S. National Science Foundation CAREER Award Grant No. 2340175, a Career Award at the Scientific Interface from BWF, and a Chan Zuckerberg Biohub – San Francisco Investigatorship. The funders had no role in study design, data collection and analysis, and decision to publish or preparation of the manuscript.

## AUTHOR CONTRIBUTIONS

T.E.G. and F.V.G. conceived the study. T.E.G., C.S.G. and J.A.N.B. supervised the work. F.V.G. designed, developed, and optimized the full experimental pipeline, performed all experiments with the recombinase enzymes and carried out the nanopore sequencing. S.K.C. developed an automated version of the pipeline and performed the automated preparation of all nanopore sequencing libraries. S.H.-N.J. helped assemble the pSpacer plasmids. J.A.N.B. provided the recombinase circuits for plants. T.E.G. developed the bioinformatics analyses and processed the sequencing data. All authors contributed to the analysis of the results and the writing of the manuscript.

## REFERENCES

[1] Voigt, C. A. (2020) Synthetic biology 2020–2030: six commercially-available products that are changing our world. Nature Communications, 11(1), 6379.

[2] Brophy, J. A. N. and Voigt, C. A. (2014) Principles of genetic circuit design. Nature Methods, 11(5), 508–520.

[3] Nielsen, A. A. K., Der, B. S., Shin, J., Vaidyanathan, P., Paralanov, V., Strychalski, E. A., Ross, D., Densmore, D., and Voigt, C. A. (2016) Genetic circuit design automation. Science, 352(6281), aac7341.

[4] Brophy, J. A. N., Magallon, K. J., Duan, L., Zhong, V., Ramachandran, P., Kniazev, K., and Dinneny, J. R. (2022) Synthetic genetic circuits as a means of reprogramming plant roots. Science, 377(6607), 747–751.

[5] Gander, M. W., Vrana, J. D., Voje, W. E., Carothers, J. M., and Klavins, E. (2017) Digital logic circuits in yeast with CRISPR-dCas9 NOR gates. Nature Communications, 8(1), 15459.

[6] Guiziou, S., Mayonove, P., and Bonnet, J. (2019) Hierarchical composition of reliable recombinase logic devices. Nature Communications, 10(1), 456.

[7] Roquet, N., Soleimany, A. P., Ferris, A. C., Aaronson, S., and Lu, T. K. (2016) Synthetic recombinase-based state machines in living cells. Science, 353(6297), aad8559.

[8] Lloyd, J. P. B., Ly, F., Gong, P., Pflueger, J., Swain, T., Pflueger, C., Fourie, E., Khan, M. A., Kidd, B. N., and Lister, R. (2022) Synthetic memory circuits for stable cell reprogramming in plants. Nature Biotechnology, 40(12), 1862–1872.

[9] Weinberg, B. H., Pham, N. T. H., Caraballo, L. D., Lozanoski, T., Engel, A., Bhatia, S., and Wong, W. W. (2017) Large-scale design of robust genetic circuits with multiple inputs and outputs for mammalian cells. Nature Biotechnology, 35(5), 453–462.

[10] Bonnet, J., Yin, P., Ortiz, M. E., Subsoontorn, P., and Endy, D. (2013) Amplifying Genetic Logic Gates. Science, 340(6132), 599–603.

[11] Cardinale, S. and Arkin, A. P. (2012) Contextualizing context for synthetic biology – identifying causes of failure of synthetic biological systems. Biotechnology Journal, 7(7), 856–866.

[12] Ceroni, F., Algar, R., Stan, G.-B., and Ellis, T. (2015) Quantifying cellular capacity identifies gene expression designs with reduced burden. Nature Methods, 12(5), 415–418.

[13] Gorochowski, T. E., Espah Borujeni, A., Park, Y., Nielsen, A. A., Zhang, J., Der, B. S., Gordon, D. B., and Voigt, C. A. (2017) Genetic circuit characterization and debugging using RNA-seq. Molecular Systems Biology, 13(11), 952.

[14] Espah Borujeni, A., Zhang, J., Doosthosseini, H., Nielsen, A. A. K., and Voigt, C. A. (2020) Genetic circuit characterization by inferring RNA polymerase movement and ribosome usage. Nature Communications, 11(1), 5001.

[15] LaFleur, T. L., Hossain, A., and Salis, H. M. (2022) Automated model-predictive design of synthetic promoters to control transcriptional profiles in bacteria. Nature Communications, 13(1), 5159.

[16] Yang, L., Nielsen, A. A. K., Fernandez-Rodriguez, J., McClune, C. J., Laub, M. T., Lu, T. K., and Voigt, C. A. (2014) Permanent genetic memory with □1-byte capacity. Nature Methods, 11(12), 1261–1266.

[17] Olorunniji, F. J., Rosser, S. J., and Stark, W. M. (2016) Site-specific recombinases: molecular machines for the Genetic Revolution. Biochemical Journal, 473(6), 673–684.

[18] Fernandez-Rodriguez, J., Yang, L., Gorochowski, T. E., Gordon, D. B., and Voigt, C. A. (2015) Memory and Combinatorial Logic Based on DNA Inversions: Dynamics and Evolutionary Stability. ACS Synthetic Biology, 4(12), 1361–1372.

[19] Short, A. E., Kim, D., Milner, P. T., and Wilson, C. J. (2023) Next generation synthetic memory via intercepting recombinase function. Nature Communications, 14(1), 5255.

[20] Huang, B. D., Kim, D., Yu, Y., and Wilson, C. J. (2024) Engineering intelligent chassis cells via recombinase-based MEMORY circuits. Nature Communications, 15(1), 2418.

[21] Courbet, A., Endy, D., Renard, E., Molina, F., and Bonnet, J. (2015) Detection of pathological biomarkers in human clinical samples via amplifying genetic switches and logic gates. Science Translational Medicine, 7(289), 289ra83–289ra83.

[22] Gomide, M. S., Sales, T. T., Barros, L. R. C., Limia, C. G., de Oliveira, M. A., Florentino, L. H., Barros, L. M. G., Robledo, M. L., José, G. P. C., Almeida, M. S. M., Lima, R. N., Rehen, S. K., Lacorte, C., Melo, E. O., Murad, A. M., Bonamino, M. H., Coelho, C. M., and Rech, E. (2020) Genetic switches designed for eukaryotic cells and controlled by serine integrases. Commun Biol, 3(255).

[23] Zúñiga, A., Guiziou, S., Mayonove, P., Meriem, Z. B., Camacho, M., Moreau, V., Ciandrini, L., Hersen, P., and Bonnet, J. (2020) Rational programming of history-dependent logic in cellular populations. Nature Communications, 11(1), 4758.

[24] Tarnowski, M. J. and Gorochowski, T. E. (2022) Massively parallel characterization of engineered transcript isoforms using direct RNA sequencing. Nature Communications, 13(1), 434.

[25] Durrant, M. G., Fanton, A., Tycko, J., Hinks, M., Chandrasekaran, S. S., Perry, N. T., Schaepe, J., Du, P. P., Lotfy, P., Bassik, M. C., Bintu, L., Bhatt, A. S., and Hsu, P. D. (2023) Systematic discovery of recombinases for efficient integration of large DNA sequences into the human genome. Nature Biotechnology, 41(4), 488–499.

[26] Guiziou, S., Maranas, C. J., Chu, J. C., and Nemhauser, J. L. (2023) An integrase toolbox to record gene-expression during plant development. Nature Communications, 14(1), 1844.

[27] Maranas, C. J., George, W., Scallon, S. K., VanGilder, S., Nemhauser, J. L., and Guiziou, S. (2023) A history-dependent integrase recorder of plant gene expression with single-cell resolution. Nature Communications, 15(1), 9362.

[28] De Rybel, B., Vassileva, V., Parizot, B., Demeulenaere, M., Grunewald, W., Audenaert, D., Van Campenhout, J., Overvoorde, P., Jansen, L., Vanneste, S., Möller, B., Wilson, M., Holman, T., Van Isterdael, G., Brunoud, G., Vuylsteke, M., Vernoux, T., De Veylder, L., Inzé, D., Weijers, D., Bennett, M. J., and Beeckman, T. (2010) A Novel Aux/IAA28 Signaling Cascade Activates GATA23-Dependent Specification of Lateral Root Founder Cell Identity. Current Biology, 20(19), 1697–1706.

[29] Sheth, R. U., Yim, S. S., Wu, F. L., and Wang, H. H. (2017) Multiplex recording of cellular events over time on CRISPR biological tape. Science, 358(6369), 1457–1461.

[30] Brown, W. R., Lee, N. C., Xu, Z., and Smith, M. C. (2011) Serine recombinases as tools for genome engineering. Methods, 53(4), 372–379 Methods For Extracting Function From Mammalian Genomes.

[31] Gowers, G.-O. F., Vince, O., Charles, J.-H., Klarenberg, I., Ellis, T., and Edwards, A. (2019) Entirely Off-Grid and Solar-Powered DNA Sequencing of Microbial Communities during an Ice Cap Traverse Expedition. Genes, 10(11).

[32] Merrick, C. A., Zhao, J., and Rosser, S. J. (2018) Serine Integrases: Advancing Synthetic Biology. ACS Synthetic Biology, 7(2), 299–310.

[33] Metsky, H. C., Siddle, K. J., Gladden-Young, A., Qu, J., Yang, D. K., Brehio, P., Goldfarb, A., Piantadosi, A., Wohl, S., Carter, A., Lin, A. E., Barnes, K. G., Tully, D. C., Corleis, B., Hennigan, S., Barbosa-Lima, G., Vieira, Y. R., Paul, L. M., Tan, A. L., Garcia, K. F., Parham, L. A., Odia, I., Eromon, P., Folarin, O. A., Goba, A., Simon-Lorière, E., Hensley, L., Balmaseda, A., Harris, E., Kwon, D. S., Allen, T. M., Runstadler, J. A., Smole, S., Bozza, F. A., Souza, T. M. L., Isern, S., Michael, S. F., Lorenzana, I., Gehrke, L., Bosch, I., Ebel, G., Grant, D. S., Happi, C. T., Park, D. J., Gnirke, A., Sabeti, P. C., Matranga, C. B., and Consortium, V. H. F. (2019) Capturing sequence diversity in metagenomes with comprehensive and scalable probe design. Nature Biotechnology, 37(2), 160–168.

[34] Nikolaeva-Reynolds, L., Cammies, C., Crichton, R., and Gorochowski, T. E. (2025) Cas9-based en-richment for targeted long-read metabarcoding. Royal Society Open Science, 12(4), 242110.

[35] Gorochowski, T. E., Chelysheva, I., Eriksen, M., Nair, P., Pedersen, S., and Ignatova, Z. (2019) Absolute quantification of translational regulation and burden using combined sequencing approaches. Molecular Systems Biology, 15(5), e8719.

[36] Clough, S. J. and Bent, A. F. (1998) Floral dip: a simplified method for Agrobacterium-mediated transformation of Arabidopsis thaliana. The Plant Journal, 16(6), 735–743.

