## Supplementary Information for "Characterization of recombinase-based genetic parts and circuits using nanopore sequencing"

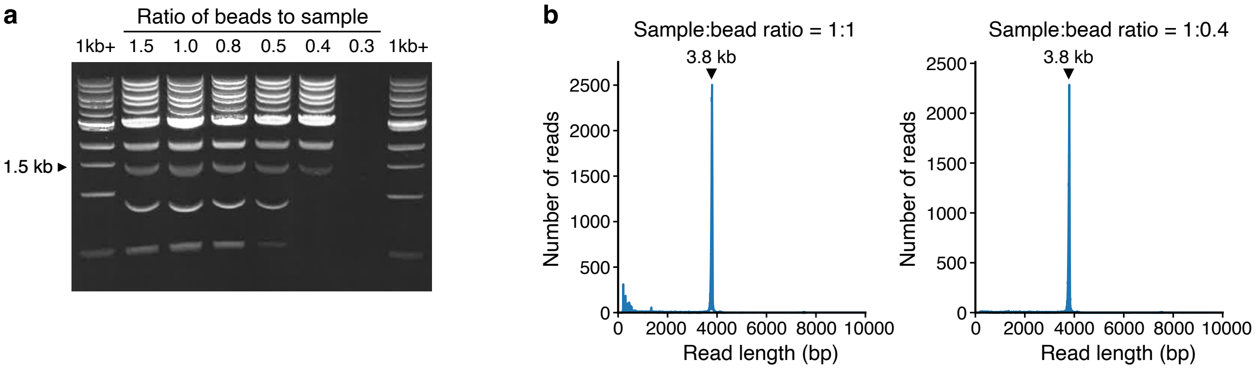

**Supplementary Figure 1: DNA size selection using magnetic beads.** (**a**) Gel showing the DNA products of a magnetic bead clean-up of a 1 kb ladder using different sample to bead ratios. (**b**) Nanopore sequencing reads from a sample containing a 3.8kb circuit amplicon (highlighted on plots) with clean-up using sample to bead ratios of 1:1 (left) and 1:0.4 (right). Expected peak at 3.8 kb highlighted with black filled triangle. Note the removal of small unwanted DNA fragments when a sample:bead ratio of 1:0.4 is used (left plot).

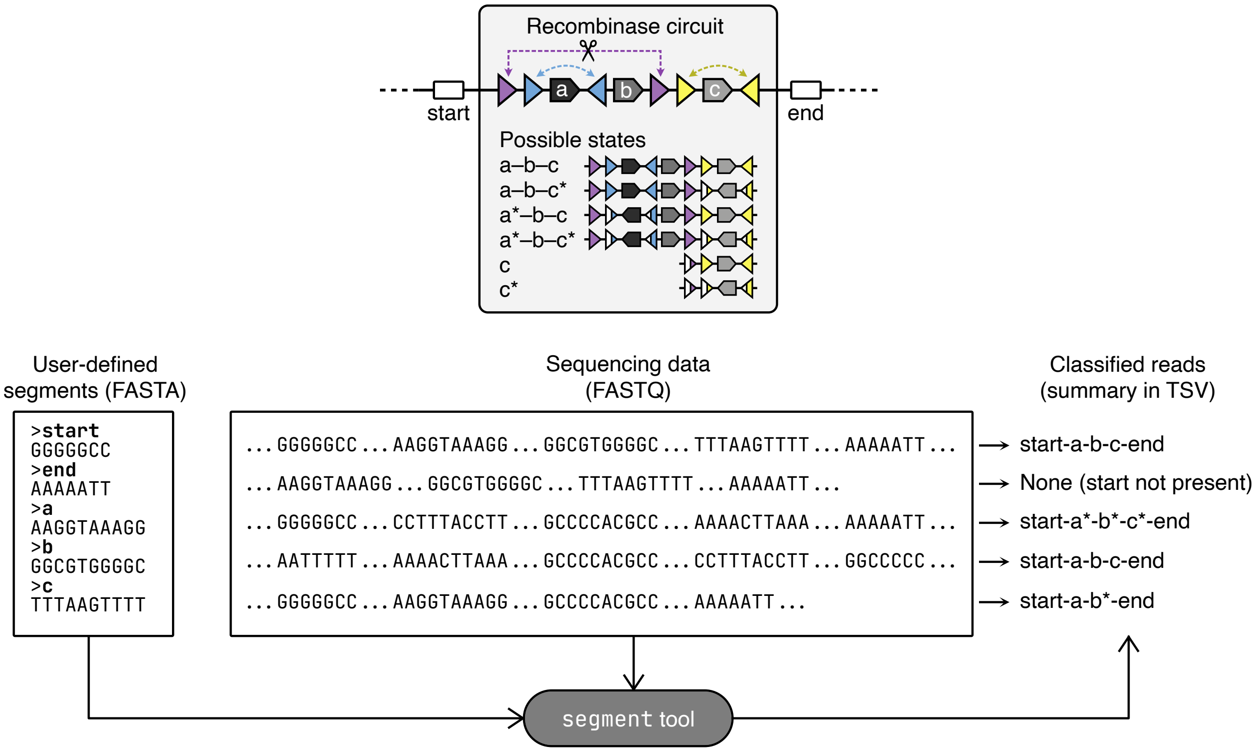

**Supplementary Figure 2: Segment analysis tool.** For a given recombinase circuit (example shown at top), the user must define a set of segments (typically short DNA sequences >30 bp in length) that are themselves unique and which can be used to classify the full state of the circuit (i.e., they capture points at which changes in the circuit that change). In addition, a 'start' and 'end' segment sequence must be provided to ensure a read captures the entire region of interest and to enable correct orientation of the read (as reads could occur in either direction for a specific amplicon). User-define segment sequences are provided in a multi-FASTA format and sequencing data from the experimental workflow is in a FASTQ format. Both these files are provided to the `segment` tool, which then classifies each read in terms of the order and orientation of the segments found. This allows for a unique state ID to be generated where segment names (from the FASTA file) are concatenated with a hyphen and orientation of the segment is encoded in the segment name, being post-fixed with an asterisk if that segment has a reverse orientation. Once all reads are classified a summary of all state IDs found with their count is provided as a text-based TSV file. We always present the state IDs from 'start' to 'end' to ensure each is unique.

### Supplementary Table 1: Overview of the automated experimental workflow

| Step | Protocol | Description | Modules[a] | Pipettes[b] | Runtime[c] |
|---|---|---|---|---|---|
| 1. Tailing PCR | 1 | Preparing mastermix in reservoir. | — | P300, P1000 | 7 m |
| | 2 | Heat lysis of samples and tailing PCR reaction. | TM, TC | M20, M300 | 3 h 32 m |
| 2. DNA purification | 3 | Purification of DNA from tailing PCR reactions (20 µL). | MM | M20, M300 | 1 h 51 m |
| 3. DNA Quantification | 4 | Preparing DNA standard curve dilution series. | — | P20 | 7 m |
| | 5 | Adding eluted DNA and standards to Quantifluor dye. **Output:** File containing DNA concentrations for use in Protocol 6 (next step) | — | M20, M300 | 18 m |
| 4. Barcoding PCR | 6 | Calculating the dilutions for each sample and picking any out of the plate too low to dilute 1 in 7. **Input:** File containing DNA concentrations from previous step. | — | P20 | 5 m |
| | 7 | Diluting all samples left in the original elution plate 1 in 7. | — | M20, M300 | 15 m |
| | 8 | Diluting each sample individually to 1.18 ng/µL for PCR. | — | M20, M300 | 1 h 32 m |
| | 9 | Barcoding PCR reaction. | TC | M20, M300 | 1h 45m |
| 5. DNA purification | 10 | Purification of DNA from barcoding PCR reaction. (40 µL) | MM | M20, M300 | 2 h 5 m |
| 6. DNA Quantification | 5 | Adding eluted DNA and standards to Quantifluor dye. **Output:** File containing DNA concentrations for use in Protocol 11 (next step) | — | M20, M300 | 18 m |
| 7. Sequencing libraries | 11 | Calculating the amount of DNA needed from each sample in each flow cell, diluting to that concentration and combining into two tubes for each flow cell. **Input:** File containing DNA concentrations from previous step. | — | M20, M300 | 1 h 45 m |

a. TM = Temperature Module, TC = Thermocycler, MM = Magnetic Module
b. P = single channel pipette, M = multi-channel pipette
c. Protocol runtime on the OT-2 only in hours (h) and minutes (m). Does not include preparation time or time taken to calibrate prior to running.

#### Supplementary Table 2: Cost of consumables for the automated workflow

| Description | Manufacturer | Product code | Quantity | Item cost | Total cost |
|---|---|---|---|---|---|
| 20 uL Tip Racks | Opentrons | 999-00014 | 28 | £3.27 | £91.84 |
| 300 uL Tip Racks | Opentrons | 999-00015 | 18 | £3.27 | £58.86 |
| 1000 uL Tip Racks | Opentrons | 999-00016 | 1 | £3.27 | £3.27 |
| 12-Channel Reservoirs | NEST | 999-00076 | 5 | £3.35 | £16.75 |
| Polyolefin PCR Seal | StarLab | E2796-9793 | 2 | £0.42 | £0.84 |
| 0.1 mL PCR Plate | NEST | 999-00050 | 6 | £1.88 | £11.28 |
| 0.2 mL PCR Plate | StarLab | E1403-5200 | 7 | £1.82 | £12.74 |
| 0.2 mL PCR 8-Strip Tubes | StarLab | A1402-3700 | 12 | £0.41 | £4.92 |
| 96-Well Flat, Black, Clear-Bottom Microplates | Greiner | 655087 | 4 | £2.84 | £11.36 |
| 1.5 mL LoBind Tubes | Eppendorf | 0030.108.051 | 6 | £0.05 | £0.10 |
| 1.5 mL Tubes | StarLab | S1615-5500 | 5 | £0.02 | £0.10 |

**Supplementary Table 3: Parameters of Hill function fits for recombinase characterization[a]**

| Recombinase | Plasmid | Segment Length | $y_{\min}$ (%) | $y_{\max}$ (%) | $K$ (min) | $n$ |
|---|---|---|---|---|---|---|
| Int2 | pMemoryArray | 50 bp | 0.0 | 95.5 | 109 | 9.4 |
| Int3 | pMemoryArray | 50 bp | 0.0 | 95.6 | 76 | 20.2 |
| Int4 | pMemoryArray | 50 bp | 0.0 | 100.0 | 41 | 3.9 |
| Int5 | pMemoryArray | 50 bp | 0.0 | 96.0 | 69 | 12.0 |
| Int7 | pMemoryArray | 50 bp | 0.0 | 93.1 | 99 | 7.4 |
| Int8 | pMemoryArray | 50 bp | 0.0 | 98.6 | 73 | 16.2 |
| Int2 | pSpacer600 | 600 bp | 0.0 | 94.0 | 125 | 19.1 |
| Int2 | pSpacer700 | 700 bp | 0.0 | 91.0 | 128 | 23.0 |
| Int2 | pSpacer800 | 800 bp | 0.0 | 92.0 | 121 | 21.9 |
| Int2 | pSpacer900 | 900 bp | 0.0 | 94.9 | 125 | 21.8 |

a. Hill function had the form: $y(x) = y_{\min} + (y_{\max} - y_{\min}) \frac{x^n}{K^n + x^n}$, where $y(x)$ is the flipped % at time $x$ minutes, $y_{\min}$ is the minimum flipped %, $y_{\max}$ is the maximum flipped %, $K$ is the time to reach 50% flipped, and $n$ is the cooperativity.

#### Supplementary Table 4: Primer sequences used for tailing PCRs

| Target | Primer | Primer Sequence[a] |
|---|---|---|
| pMemoryArray | prMemory-F | **TTTCTGTTGGTGCTGATATTGC**TGAAACTCACCCAGGGATTG |
| | prMemory-R | **CTTGCCTGTCGCTCTATCTTC**CGATTCAGGTTCATCATGCCGT |
| pSpacerX | prSpacer-F | **TTTCTGTTGGTGCTGATATTGC**TGAAACTCACCCAGGGATTG |
| | prSpacer-R | **CTTGCCTGTCGCTCTATCTTC**CGATTCAGGTTCATCATGCCGT |
| pCas2+5 | pCasscade1-F | **TTTCTGTTGGTGCTGATATTGC**CTCAGGCGCAATCACGAATG |
| | pCasscade1-R | **ACTTGCCTGTCGCTCTATCTTC**GAGGCATAAATTCCGTCAGC |
| pCas7+gfp | pCasscade2-F | **TTTCTGTTGGTGCTGATATTGC**TGAAACTCACCCAGGGATTG |
| | pCasscade2-R | **CTTGCCTGTCGCTCTATCTTC**CGATTCAGGTTCATCATGCCGT |
| pGATA23+PhiC31 | prRoot-F | **TTTCTGTTGGTGCTGATATTGC**GATTGAATCCTGTTGCCGGTCTTG |
| | prRoot-R | **CTTGCCTGTCGCTCTATCTTC**GTAACGGGAGAAGCACTGCAC |

a. Bold sections denote the common ends added to the amplicons for the subsequent barcoding step.

### Supplementary Note 1: Automated experimental workflow for Opentrons OT-2

We developed a set of protocols for the Opentrons OT-2 robot to automate the majority of the experimental workflow. The automated workflow consists of 11 different protocols covering the preparation of reagents, heat-based cell lysis, PCR, DNA clean-up and DNA quantification steps (see **Supplementary Table 1** for details). This workflow has been run using a single OT-2 over two extended days where the pipettes and deck were reconfigured after each protocol. When running the entire workflow for the first time using a single OT-2, we advise allowing for at least 3 days, which includes setting up the nanopore sequencing. It is possible to reduce the overall running time of the workflow by using two OT-2s: one with both multi-channels installed for use with all three modules interchangeably (Thermocycler, Magnetic Module, Temperature Module), and the other with single channel pipettes installed for dilutions and reagent preparation steps. This enables only one pipette change being required and allows for some protocols to be run in parallel and set up in advance. Using two OT-2s, the pipeline can be run comfortably in two working days, which includes setting up the nanopore sequencing.

**Custom Labware Files:** The protocols in this pipeline include labware that is not part of the standard labware library in the Opentrons Python API. To use these, download from the GitHub repository (or **Supplementary Data 1**) and move into your custom labware definitions folder.

**Transferring Files to and from the OT-2:** During the workflow there are input files required by certain steps (Protocols 6, 8 and 11) and output files generated at several steps (Protocols 6 and 11). You should ensure you have SSH access set up to the OT-2 and can use SCP to transfer files via the command line between your computer and the OT-2. Instructions on how to do this can be found on the Opentrons support site. We also expect there to be two directories present on the OT-2 to allow for input and output files, respectively:

- `/data/user_storage/input_files`
- `/data/user_storage/output_files`

**Calibration:** Before each protocol was run a calibration step was performed to check alignments of all labware used and to apply any offsets where needed.

### OT-2 deck configurations

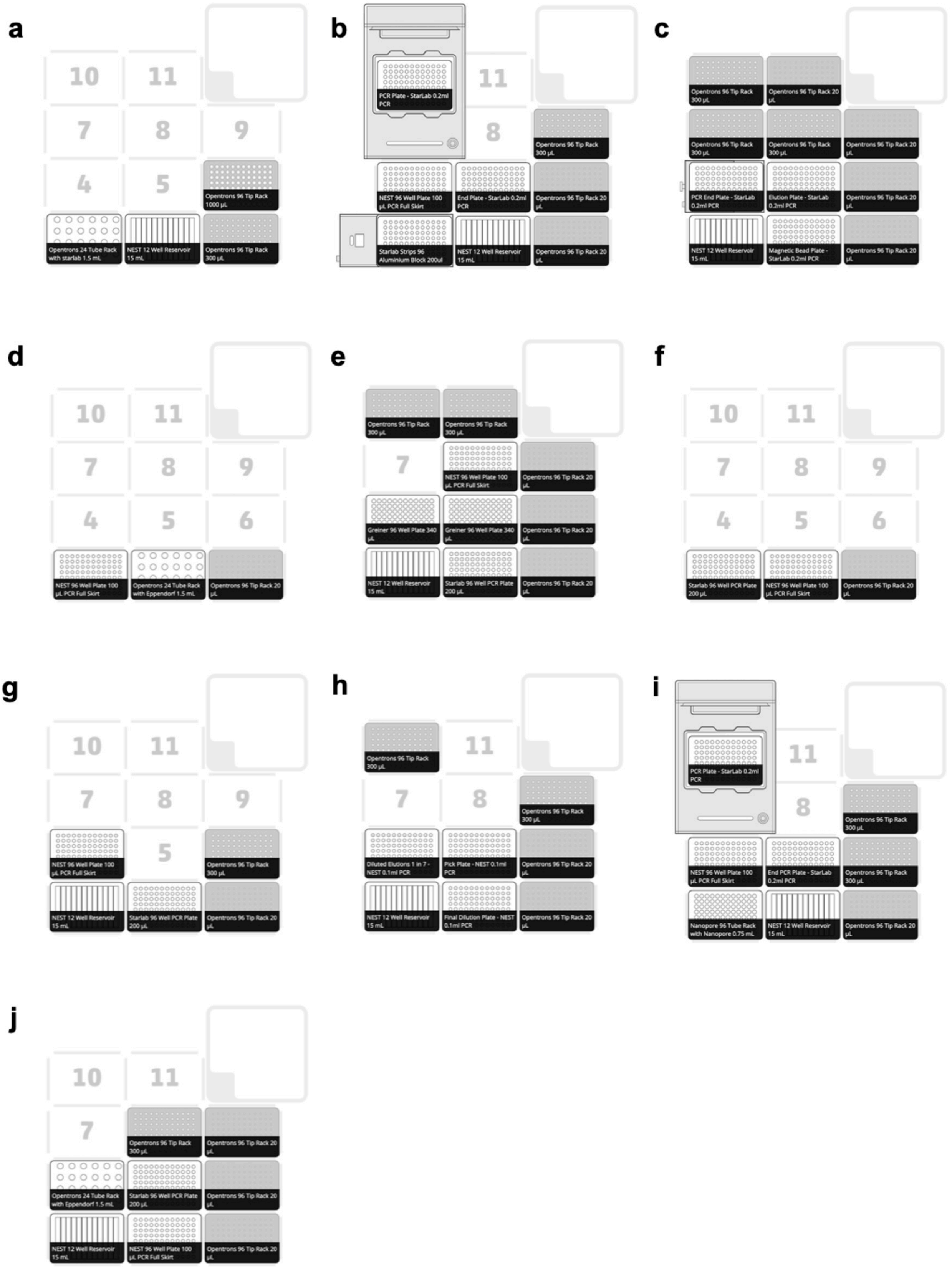

### Protocol 1: Prepare mastermix for tailing PCR

---

**Script Name:** `1_Prep_Tailing_PCR_Mastermix.py`

**Description:** Prepares the mastermix for 96 tailing PCR reactions into a reservoir to allow easy transfer by multi-channel to the PCR plate in Protocol 2.

**Runtime:** 7 minutes

**Pipettes Required:** P300 Single GEN2 and P1000 Single GEN2

**Modules Required:** None

**Labware Required:** 1 × 12-channel reservoir (999-00076, NEST), 1 × 4-in-1 Rack with 24-Tube insert (999-00050, Opentrons), 5 × 1.5 mL microcentrifuge Tubes (S1615-5500, StarLab), 1 × P300 Tip Rack (999-00015, Opentrons), 1 × P1000 Tip Rack (999-00016, Opentrons).

**Reagents Required:** Nuclease Free Water (PD092, BioTek), LongAmp Taq 2X Master Mix (M0287L, NEB), Forward and Reverse Primers for the PCR reaction.

**Preparation Required:** Load 170 µL of each of the primers into a 1.5 mL tube, 800 µL of Taq into one tube and 700 µL into two tubes. Place all tubes into the tube rack into correct positions. A1 = Forward Primer, B1 = Reverse Primer, A2 = Taq (800 µL), B2 & C2 = Taq (700 µL). In the reservoir add 3 mL of nuclease free water (NFW) in column 12.

**Deck configuration:** A

**Protocol Steps:**

1. The P1000 transfers 1266.5 µL of NFW to column 1 of the reservoir in 2 transfers.
2. The p300 mixes the forward primers 3x and transfers 149 µL into column 1 and mixes 5x. This is process is repeated with the reverse primers.
3. The P1000 transfers 1862.5 µL of LongAmp Taq polymerase in 3 transfers from 3 tubes into column 1, and mixes 10x using slower liquid handling controls to prevent bubbles and incorrect aspiration.

### Protocol 2: Tailing PCR reaction from mastermix

---

**Script Name:** `2_PCR_Tailing_Reaction.py`

**Description:** Performs heat lysis of cells and tailing PCR reaction.

**Runtime:** 3 hours 32 minutes

**Pipettes Required:** P20 Multi GEN2 and P300 Multi GEN2

**Modules Required:** Thermocycler and Temperature Module

**Labware Required:** 12 × 0.2 mL PCR Tube Strips (A1402-3700, StarLab), 2 × 0.2 mL PCR plates (E1403-5200, StarLab), 1 × 0.1 mL PCR plate (999-00050, NEST), 1 × PCR Tube Aluminium Block (Opentrons), 1 × 12-Channel Reservoir (999-00016, NEST), 2 × 20 µL Tip Racks (999-00014, Opentrons), 1 × 300ul Tip Racks (999-00015, Opentrons), 2 × PCR Tube Racks (TRC9601, BioRad), 1 × Polyolefin StarSeal (E2796-9793, StarLab).

**Reagents Required:** Samples (25 µL) in PCR Tube Strips on PCR rack and mastermix from Protocol 1 in column 1 of reservoir.

**Other Equipment Required:** Plate centrifuge (5810 R, Eppendorf)

**Preparation Required:** Fill PCR aluminum block wells with 20 µL of MilliQ water. Set the temperature of the Temperature block to 95°C manually in Opentrons App 10 mins before starting the protocol to allow the block to come to temperature. Load Samples in the PCR rack to the aluminum block and calibrate before starting. The PCR rack is not included the labware definition because it does not alter the fit of tubes into the block, the rack is only used here to keep tubes together for easy transport from the deck to the centrifuge.

**Deck configuration:** B

#### Protocol Steps:

1. Temperature module is held at 95°C for 10 minutes to heat-lyse the cells.
2. The temperature module is cooled to 20°C.
3. The protocol is paused and the PCR rack with samples in is removed from the module manually by the user and centrifuged at 4,000 rpm for 2 minutes and placed back onto the temperature module.
4. The P300 multi-channel transfers 23 µL of mastermix into the PCR plate on the Thermocycler.

5. The P20 multi-channel aspirates 12 µL of supernatant from 2 mm above the bottom of each PCR tube to avoid touching the pellet and 10 µL is dispensed into the corresponding well in a plate for storage. The last 2 µL is dispensed into the corresponding well of the PCR plate on the Thermocycler and mixed 5x.
6. The protocol is paused, and a polyolefin seal is manually applied to the PCR plate and secured in place with a hand roller. The storage plate is removed from the deck, sealed, and stored at 4°C in case the pipeline needs to be restarted.
7. The protocol is resumed by the user, the thermocycler lid closes, and the lid temperature is set to 105°C. The thermocycler block temperature is set to 94°C and then held for 5 minutes.
8. The thermocycler then cycles through 30 cycles of 94°C for 20 seconds, 60°C for 15 seconds then 65°C for 3 minutes and 16 seconds.
9. The thermocycler block is then set 65°C and held for 10 minutes.
10. The thermocycler lid is deactivated, the block is then set to 20°C and the lid is opened.
11. The protocol pauses and the user removes the seal.
12. The P20 multi-channel transfers 20 µL from each well into the corresponding well of a fresh PCR plate.

### Protocol 3: DNA purification of tailing PCR products

---

**Script Name:** `3_Pur_DNA_Mag_20ul.py`

**Description:** DNA purification of tailing PCR products (20 µL) using KAPA beads and the Opentrons Magnetic Module.

**Runtime:** 1 hour 51 minutes

**Pipettes required:** P20 Multi GEN2 and P300 Multi GEN2

**Modules required:** Magnetic Module GEN2

**Labware Required:** 3 × 0.2 mL PCR Plates (E1403-5200, StarLab), 1 × 12-Channel reservoir (999-00016, NEST), 7 × 20µLTip Racks (999-00014, Opentrons), 3 × 300µLTip Racks (999-00015, Opentrons).

**Reagents Required:** KAPA Pure Magnetic Beads (KK8002, KAPA Biosystems), 40 mLs of 80% Ethanol (freshly prepared), Nuclease Free Water (PD092, BioTek). Plate with PCR products from Protocol 2.

**Preparation Required:** Load 105 µL of magnetic beads (room temperature, vortexed) into each well of column 1 in one of the 0.2 mL PCR plates. Load 10 mL of 80% ethanol into columns 1 to 4 of reservoir. Add 2 mL of nuclease-free water into column 5 of reservoir. Start with the end plate from the PCR reaction on deck slot 5 instead of on the magnetic module to allow full suspension of the beads in the product without pelleting.

**Deck configuration:** C

**Protocol Steps:**

1. The P20 multi-channel transfers 8 µL of magnetic beads from column 1 into all wells of the PCR end plate on deck slot 5 and mixes first 10x at 2.5x speed and at different aspiration and dispense heights then again at the bottom of the well 5x at 1x speed.
2. The protocol is then delayed for 10 minutes for an incubation.
3. The protocol is paused, the PCR end plate is moved from deck slot 5 to the magnetic module on deck slot 4, a clean PCR plate is placed on slot 5 to collect the eluted DNA.
4. The user resumes the protocol and magnet on the module is engaged to 8 mm and the protocol is delayed for 4 minutes to allow pelleting.

5. The P20 multi-channel aspirates the supernatant from the side of each well (odd columns on the left and even on the right to avoid picking the beads) of each column in 2 transfers and dispenses the waste into the reservoir (column 8) with an air gap of 2 µL between each transfer to prevent dripping from tips over the plate.
6. The P300 multi-channel adds 200 µL of ethanol from columns 1 and 2 to each well across the whole plate.
7. The ethanol is then removed from the first half of the plate (columns 1 to 6) only to prevent the beads from drying out during this process. The removal of ethanol process is the same throughout the protocol. It first uses the P300 multi-channel to remove 180 µL to the reservoir (column 9), then the P20 multi-channel is used to aspirate 15 µL out of the well into the reservoir, the pipette then shakes over the waste channels to remove any drips of ethanol before aspirating a final 5 µL out of the well.
8. Ethanol is then added to the first half of the plate.
9. Ethanol is removed from the second half of the plate (columns 7 to 12).
10. The protocol pauses to allow the user to empty the tip waste and refill the P20 tip racks (slots 3, 6, 9).
11. Ethanol is added to the second half of the plate.
12. Ethanol is then removed from the first half of the plate and the protocol is delayed for 2 minutes to allow drying.
13. The P20 multi-channel then adds 16 µL of ultrapure water to the first half of the plate.
14. The protocol pauses to allow the user to empty the tip waste.
15. Ethanol is removed from the second half of the plate and the protocol is delayed for 2 minutes.
16. The P20 multi-channel then adds 16 µL of ultrapure water to the second half of the plate.
17. The magnetic module is disengaged.
18. The P20 multi-channel mixes each well in the plate 20x at 10x speed from the bottom of the well to resuspend the beads in the ultrapure water.
19. The protocol is delayed for 10 minutes.
20. The magnetic module is engaged at 8 mm and the protocol delayed for 2 minutes to allow pelleting.
21. The P20 multi-channel then transfers 13 µL of eluate from the side of each well to the corresponding well in the elution plate on slot 5.

### Protocol 4: Preparing DNA standards for quantification

---

**Script Name:** `4_Prep_Quantifluor_Stds.py`

**Description:** Diluting dsDNA standard from the Quantifluor dsDNA ONE kit, in a 96-well plate, to create a standard curve (0.2–400 ng/µL) for DNA quantification that can easily be transferred by multi-channel in Protocol 5. This follows the manufacturer's instructions except the dilution is directly in a multi-well plate. This protocol makes enough of the standards for both DNA quantification steps and can be kept overnight at 4°C in between.

**Runtime:** 7 minutes

**Pipettes required:** P20 Single GEN2

**Modules required:** None

**Labware Required:** 1 × 0.1 mL PCR Plate (999-00050, NEST), 2 × 1.5 mL LoBind Tubes (0030 108.051, Eppendorf), 1 × 4-in-1 Rack with 24-Tube insert (999-00050, Opentrons), 1 × 20 µL Tip Rack (999-00014, Opentrons)

**Reagents Required:** Lamda DNA standard and 1X TE Buffer from Quantifluor ds DNA ONE kit (E4870, Promega)

**Preparation Required:** Load 50 µL of dsDNA (400 ng/µL) to one of the tubes and 150 µL of 1X TE Buffer in the other. Load to rack in set positions: A1 = TE Buffer, C1 = dsDNA Standard. The user can set COL_1 = to the column in number they would like the standards to be made in.

**Deck configuration:** D

**Protocol Steps:**

1. The P20 pre-wets the tip in the TE buffer by mixing 3x and transfers 10 µL to well B of the column the standards will be made in, then 15 µL to wells C to H using the same tip.
2. The P20 mixes the dsDNA standard 5x before transferring 30 µL to well A of the plate.
3. The P20 transfer 10 µL from well A to well B and mixes 10x at 2.5x speed for a 1 in 2 dilution.
4. Using fresh tips each time the P20 then transfers 5 µL from well B to well C and mixes 10x at 2.5x speed creating a 1 in 4 dilution. This is repeated from each well to the next until well G. Well H remains only 15 µL of TE buffer as a blank.

### Protocol 5: Preparing DNA samples for quantification

---

**Script Name:** `5_Prep_Quantifluor_Samples.py`

**Description:** Adds unknown concentrations of purified DNA and a set of standards to Quantifluor dsDNA ONE dye in two 96-well plates to prepare for quantification using a fluorescent plate reader.

**Runtime:** 18 minutes

**Pipettes required:** P20 Multi GEN2, P300 Multi GEN2

**Modules required:** None

**Labware Required:** 1 × 12-Channel Reservoir (999-00076, NEST), 2 × Black, Clear-Bottomed 96 Well Plates (655087, Greiner), 2 × 20 µLTip Racks (999-00014, Opentrons), 2 × 300 µL Tip Racks (999-00015, Opentrons), 2 × Foil Seals, 1 × 0.2 mL PCR plate (E1403-5200, StarLab) with eluted DNA from either purification protocol, 1 × 0.1 mL PCR plate (999-00050, NEST) with Quantifluor standards prepared in (prepared in Protocol 4).

**Reagents Required:** QuantiFluor ONE dsDNA Dye from kit (E4870, Promega)

**External equipment required:** Vortex mixer with microplate adapter (SA8, Stuart), plate reader like a SpectraMax with 485 nm and 535 nm filters (iD5, Molecular Devices).

**Preparation Required:** Fill channel 1 and 2 in reservoir with 10.5 mL of Quantifluor Dye and channel 3 with 7.5 mL. To ensure standards solutions are homogenous, just before protocol mix plate on vortex mixer at 600 rpm for at least 5 seconds. 1. Set the variable STDS_COL equal to well A of the column you have prepared the Quantifluor standards in. For example, STDS_COL = 'A1' for column 1. Calibrate before starting and for best results keep OT-2 deck lights off and avoid turning bright lights on directly above the OT-2.

**Deck configuration:** E

#### Protocol Steps:

1. Protocol pauses and asks user to check standards column in protocol corresponds to what is observed in real life before resuming.
2. The P300 multi-channel pre-wets the tips with Quantifluor dye by mixing 3x. It then transfers 200 µL of dye from the reservoir to all the wells in plate 1 (reservoir channel 1 fills columns 1 to 6, channel 2 fills columns 7 to 12) and then the first 4 columns in plate 2 from channel 3.

3. The P20 multi-channel mixes the standards 5x.
4. The P20 adds 1 µL of each standard to column 1 and then 12 in plate 1.
5. The P20 adds 1 µL of each standard to column 1 and then 4 in plate 2.
6. The P20 transfers 1 µL of each sample from the plate of eluted DNA from columns 2 to 11 to the corresponding column in plate 1.
7. The P20 transfers 1 µL of each sample from columns 1 and 12 of the plate of eluted DNA into column 2 and 3 of plate 2 respectively.
8. The P300 then mixes all wells 3x at 1.5x speed.
9. Cover plates 1 and 2 with foil seal / black lid to protect from light whilst transferring to plate reader.
10. All the next steps take place off the OT-2.
11. Incubate inside the plate reader for 5 minutes before reading each plate at 485 nm (excitation) and 535 nm (emission) from the bottom.
12. Collect the data files for each plate from the plate reader.
13. Download the Quantifluor Dye Systems Data Analysis Workbook (Single Sample Worksheet) from Promega (https://www.promega.co.uk/resources/tools/quantifluor-dye-systems-data-analysis-workbook/).
14. Set up a workbook for each plate read and follow the instructions given in the workbook to calculate the DNA concentration of each sample.
15. If running for first DNA quantification (after Protocol 3) create a CSV file to transfer the results into with two columns only (no header): column 1 = well ID (in order A1, B1, C1) of sample, column 2 = result in ng/µL, save as 'DATE_Elution_Concentrations.csv'. DATE format = YYYY-MM-DD.
16. If running for second quantification (after Protocol 10) create a CSV file to transfer the results into with three columns (no header) – column 1 = well ID (in order A1, B1, C1) of sample, column 2 = sample name/ID, column 3 = result in ng/µL. Save as 'DATE_Library_Elution_Concentrations.csv'. DATE format = YYYY-MM-DD.
17. Use SCP to transfer the CSV file to '/data/user_storage/input_files/' on the OT-2.

### Protocol 6: Calculations for sample dilutions for barcoding PCR

---

**Script Name:** `6_Prep_Barcode_Dilution_Pt_1.py`

**Description:** Take input file of DNA concentrations from Protocol 5 and prepare calculations for dilution of all samples to 1.18 ng/µL for the barcoding PCR. If any samples are below 20 ng/µL they will be picked out of the plate eluted DNA in this protocol and put into another plate to be diluted separately.

**Runtime:** 5 minutes + (depending on how many samples have concentrations below 20 ng/µL)

**Pipettes required:** P20 Single GEN2

**Modules required:** None

**Labware Required:** 1 × 0.2 mL PCR Plate (E1403-5200, StarLab) with eluted DNA in (from Protocol 3), 1 × 0.1 mL PCR Plate (999-00050, NEST), 1 × 20 µL Tip Rack (999-00014, Opentrons).

**Reagents Required:** N/A

**Preparation Required:** Ensure CSV file of the DNA concentrations is uploaded to OT-2 (see Protocol 5, steps 15–17). Calibrate before running. If the date the eluted DNA was quantified differs from the day this protocol being run, change 'DATE =' to the date DNA was quantified at the top of the script, e.g., 'DATE = YYYY-MM-DD'.

**Deck configuration:** F

**Protocol Steps:**

1. Reads in input file and creates a data frame of all samples below 19.83 ng/µL as these samples are too low to be diluted 1 in 7 in the next step.
2. The P20 transfers all 13 µL of these samples (if any) into another plate (pick plate).
3. Creates another data frame of all samples above 19.83 ng/µL and calculates what the concentration of DNA will be after the 1 in 7 dilution that will occur in Protocol 7 and how much of each dilution will needed to create a final concentration of 1.18 ng/µL in 48 µL. Writes all this information to CSV file to
   '/data/user_storage/input_files/' called 'DATE_Barcode_Dilutions.csv'. This will be used by Protocol 8.

4. Of the samples too low to be diluted 1 in 7 (now in pick plate), any samples below 5.66 ng/µL are too low to be diluted to 1.18 ng/µL in 48 µL. Regardless of final concentration 8 µL of each of these sample will be diluted in 18 µL of ultrapure water in Protocol 8. The final concentration of this dilution for any relevant wells is calculated and then this information is written to 'DATE_Below_Threshold_Dilutions.csv' in the input files directory on the OT-2.
5. Of the samples too low to be diluted 1 in 7 (now in pick plate), any samples above 5.66 µL can be diluted directly to 1.18 ng/µL in 48 µL. The volume of sample needed for each dilution is calculated and written to 'DATE_Elution_Dilutions.csv' and stored in the input files directory on the OT-2.
6. An output file with all calculations is written to '/data/user_storage/output_files/' called 'DATE_Barcode_PCR_Preparation_Calculations.csv'. This can be retrieved by the user by SCP to check and follow along in Protocol 8.

### Protocol 7: Samples dilutions for barcoding PCR

---

**Script Name:** `7_Prep_Barcode_Dilution_Pt_2.py`

**Description:** Dilute all eluted DNA 1 in 7 to allow dilution to 1.18 ng/µL in Protocol 8 without pipetting outside of pipette range.

**Runtime:** 15 minutes

**Pipettes required:** P20 Multi GEN2, P300 Multi GEN2

**Modules required:** None

**Labware Required:** 1 × 12-Well Reservoir (999-00076, NEST), 1 × 0.1 mL PCR Plate (999-00050, NEST), 1 × 20 µL Tip Rack (999-00014, Opentrons), 1 × 300 µL Tip Rack (999-00015, Opentrons), 1 × 0.2 mL PCR Plate (E1403-5200, StarLab) with purified DNA samples.

**Reagents Required:** Nuclease Free Water (PD092, BioTek)

**Preparation Required:** Load 6 mL of ultrapure water into column 12 of the reservoir. Calibrate before running.

**Deck configuration:** G

**Protocol Steps:**

1. The P300 multi-channel adds 48 µL of ultrapure water to all wells of dilution plate.
2. The P20 multi-channel adds 8 µL of each sample of eluted DNA to the corresponding well on the dilution plate and mixes 10x at 2x speed at different aspiration and dispense heights.

### Protocol 8: Sample final dilutions for barcoding PCR

---

**Script Name:** `8_Prep_Barcode_Dilution_3.py`

**Description:** Continue from barcode dilution preparation protocol part 1 and 2. Dilutes all samples to 1.18 ng/µL from known concentrations.

**Runtime:** 1 hour 32 minutes

**Pipettes required:** P20 Single GEN2, P300 Single GEN2

**Modules required:** None

**Labware Required:** 1 × 12-Well Reservoir (999-00076, NEST), 2 × 20 µL Tip Racks (999-00014, Opentrons), 2 × 300 µL Tip Racks (999-00015, Opentrons), 3 × 0.1 mL PCR Plate (999-00050, NEST) where one is the pick plate from Protocol 6, another is the 1 in 7 dilution plate from Protocol 7 and the last one is where all samples will be diluted to 1.18 ng/µL in this protocol.

**Reagents Required:** Nuclease Free Water.

**Preparation Required:** Load 6 mL of ultrapure water into column 12 of the reservoir. Protocol 6 must have been run on this OT-2 to have created the necessary input files to calculate the dilutions for each sample. If the date Protocol 6 was run differs from the day this protocol is being run, change 'DATE =' to the date Protocol 6 was run at the top of the script, e.g., 'DATE = YYYY-MM-DD'.

**Deck configuration:** H

**Protocol Steps:**

1. Reads in 'DATE_Barcode_Dilutions.csv' (all the samples diluted 1 in 7).
2. The P300 adds the calculated amount of ultrapure water to all the wells in the final dilution plate from this input file.
3. The P20 transfers the calculated volume of sample to the corresponding wells in the final dilution plate and mixes 10x and 3x speed at different aspiration and dispense heights.
4. Reads in 'DATE_Elution_Dilutions.csv' (all the samples undiluted that can be diluted directly to 1.18 ng/µL)
5. The P300 adds the calculated amount of ultrapure water to all the wells in the final dilution plate from this input file.
6. The P20 transfers the calculated volume of sample to the corresponding wells in the final dilution plate and mixes 10x and 3x speed at different aspiration and dispense heights.

7. Reads in 'DATE_Below_Threshold_Dilutions.csv' (all the samples too low to dilute to 1.18 ng/µL).
8. The P20 adds 18 µL of ultrapure water to each well in the final dilution plate from this input file.
9. The P20 adds 8 µL of each sample to the corresponding wells in the final dilution plate and mixes 10x and 3x speed at different aspiration and dispense heights.

### Protocol 9: Barcoding PCR

---

**Script Name:** `9_PCR_Barcoding_Reaction.py`

**Description:** Preparation and execution of barcoding PCR reaction.

**Runtime:** 1 hour 45 minutes

**Pipettes required:** P20 Multi GEN2, P300 Multi GEN2

**Modules required:** Thermocycler Module

**Labware Required:** 2 × 0.2 mL PCR plates (E1403-5200, StarLab), 1 × 0.1 mL PCR plate (999-00050, NEST) with DNA Samples Diluted to 1.18 ng/µL, 1 × 12-Channel Reservoir (999-00016, NEST), 1 × 20 µL Tip Rack (999-00014, Opentrons), 3 × 300 µL Tip Racks (999-00015, Opentrons), 1 × Polyolefin StarSeal (E2796-9793, StarLab).

**Reagents Required:** LongAmp Taq 2X Mastermix (M0287L), Nanopore Barcoding Primers in rack (EXP-PBC096).

**Preparation Required:** Defrost and spin nanopore barcodes plate at 4,000 rpm for 1 min to ensure all liquid is at the bottom of the tubes. Load 3.5 mL of mastermix in column 4 of reservoir. If DNA samples have been left overnight vortex before use.

**Deck configuration:** H

**Protocol Steps:**

1. The P300 multi-channel transfers 25 µL of mastermix to all wells in PCR plate on thermocycler
2. The P20 multi-channel transfers 1 µL of the barcoding primers from the Nanopore rack to the corresponding well on the PCR plate, mixes 5x at 2x speed.
3. The P300 multi-channel transfers 24 µL of each sample to the corresponding well in the PCR plate, mixes 10x at 1x rate.
4. The protocol is paused, and a polyolefin seal is manually applied to the PCR plate and secured in place with a hand roller.
5. The protocol is resumed by the user, the thermocycler lid closes, and the lid temperature is set to 105°C. The thermocycler block temperature is set to 95°C and then held for 3 minutes and 50 seconds.
6. The thermocycler then cycles through 12 cycles of 95°C for 15 seconds, 62°C for 15 seconds then 65°C for 3 minutes and 16 seconds.

7. The thermocycler block is then set 65°C and held for 4 minutes.
8. The thermocycler lid is deactivated, the block is then set to 10°C and the lid is opened.
9. The protocol pauses and the user removes the seal and replaces the rack on slot 6 with a new 300 µL tip rack.
10. The P300 multi-channel transfers 40 µL from each well into the corresponding well of a fresh PCR plate.

### Protocol 10: Magnetic bead DNA purification

---

**Script Name:** `10_Pur_DNA_Mag_40ul.py`

**Description:** DNA purification of tailing PCR products (40 µL) using KAPA beads and the Opentrons Magnetic Module.

**Runtime:** 2 hours 5 minutes

**Pipettes required:** P20 Multi GEN2 and P300 Multi GEN2

**Modules required:** Magnetic Module GEN2

**Labware Required:** 3 × 0.2 mL PCR Plates (E1403-5200, StarLab), 1 × 12-Channel reservoir (999-00016, NEST), 7 × 20 µL Tip Racks (999-00014, Opentrons), 3 × 300 µLTip Racks (999-00015, Opentrons).

**Reagents Required:** KAPA Pure Magnetic Beads (KK8002, KAPA Biosystems), 40 mL of 80% Ethanol (freshly prepared), Nuclease Free Water (PD092, BioTek). Plate with PCR products from Protocol 9.

**Preparation Required:** Load 200 µL of magnetic beads (room temperature, vortexed) into each well of column 1 in one of the 0.2 mL PCR plates. Load 10 mL of 80% ethanol into columns 1 to 4 of reservoir. Add 2 mL of ultrapure water into column 5 of reservoir. Start with the end plate from the PCR reaction on deck slot 5 instead of on the magnetic module to allow full suspension of the beads in the product without pelleting.

**Deck configuration:** C

**Protocol Steps:**

1. The P20 multi-channel transfers 16 µL of magnetic beads from column 1 into all wells of the PCR end plate on deck slot 5 and mixes first 15x at 2.5x speed and at different aspiration and dispense heights then again at the bottom of the well 5x at 1x speed.
2. The protocol is then delayed for 10 minutes for an incubation.
3. The protocol is paused, the PCR end plate is moved from deck slot 5 to the magnetic module on deck slot 4, a clean PCR plate is placed on slot 5 to collect the eluted DNA and the tip waste is emptied.
4. The user resumes the protocol and magnet on the module is engaged to 8 mm and the protocol is delayed for 2 minutes to allow pelleting.

5. The P20 multi-channel aspirates the supernatant from the side of each well (odd columns on the left and even on the right to avoid picking the beads) of each column in 4 transfers and dispenses the waste into the reservoir (column 8) with an air gap of 2 µL between each transfer to prevent dripping from tips over the plate.
6. The P300 multi-channel adds 200 µL of ethanol from columns 1 and 2 to each well across the whole plate.
7. The ethanol is then removed from the first half of the plate (columns 1 to 6) only to prevent the beads from drying out during this process. The removal of ethanol process is the same throughout the protocol. It first uses the P300 multi-channel to remove 180 µL to the reservoir (column 9), then the P20 multi-channel is used to aspirate 18 µL out of the well into the reservoir, the pipette then shakes over the waste channels to remove any drips of ethanol before aspirating a final 20 µL out of the well.
8. Ethanol is then added to the first half of the plate.
9. Ethanol is removed from the second half of the plate (columns 7 to 12).
10. The protocol pauses to allow the user to empty the tip waste and refill the P20 tip racks (slots 3, 6, 9).
11. Ethanol is added to the second half of the plate.
12. Ethanol is then removed from the first half of the plate and the protocol is delayed for 2 minutes to allow drying.
13. The P20 multi-channel then adds 17 µL of ultrapure water to the first half of the plate.
14. The protocol pauses to allow the user to empty the tip waste.
15. Ethanol is removed from the second half of the plate and the protocol is delayed for 2 minutes.
16. The P20 multi-channel then adds 17 µL of ultrapure water to the second half of the plate.
17. The magnetic module is disengaged.
18. The P20 multi-channel mixes each well in the plate 20x at 8x speed from the bottom of the well to resuspend the beads in the ultrapure water.
19. The protocol is delayed for 10 minutes.
20. The magnetic module is engaged at 8 mm and the protocol delayed for 5 minutes to allow pelleting.
21. The P20 multi-channel then transfers 13 µL of elute from the side of each well to the corresponding well in the elution plate on slot 5.
22. Run Protocol 5 to quantify DNA eluted.

### Protocol 11: DNA libraries preps for nanopore sequencing

---

**Script Name:** `11_Prep_DNA_Libraries.py`

**Description:** Creates two libraries of 1 mg DNA in 47 µL for nanopore sequencing by diluting all samples to concentrations needed to add equal amounts of DNA. Columns 1 to 6 of plate of purified DNA will be prepared into one library and columns 7 to 12 in another. To prevent pipetting outside of the P20 pipette range (below 1 µL) samples will be diluted in 10 µL to 2x amount needed per µL) and 1 µL of each sample will be added to a tube, vortexed and half the total volume added to a final library tube. Samples with a concentration between the amount of DNA needed and twice the amount needed, will be diluted to 1X concentration in 5 µL and 1 µL of each will be added directly to the final library tube.

**Runtime:** 1 hour 45 minutes

**Pipettes required:** P20 Single GEN2, P300 Single GEN2

**Modules required:** None

**Labware Required:** 1 × 0.1 mL PCR Plate (999-00050, NEST), 4 × LoBind 1.5 mL Tubes (0030 108.051, Eppendorf), 1 × 4-in-1 Rack with 24-Tube insert (999-00050, Opentrons), 1 × 12-Channel Reservoir (999-00076, NEST), 3 × P20 Tip Racks (999-00014, Opentrons), 1 × P300 Tip Rack (999-00015, Opentrons).

**Reagents Required:** Nuclease free water (PD092, BioTek)

**Preparation Required:** Ensure CSV file of the DNA concentrations is uploaded to OT-2 (see Protocol 5 steps 15-17). Place 4 tubes into rack into the set positions A1 = samples for flow cell 1 at 2x concentration, A2 = samples for flow cell 1 at 1x concentration, C1 = samples for flow cell 2 at 2x concentration, C2 = samples for flow cell 2 at 1x concentration. Calibrate before running. If the date the eluted DNA was quantified differs from the day this protocol being run, change 'DATE =' to the date DNA was quantified at the top of the script, e.g., 'DATE = YYYY-MM-DD'.

**Deck configuration:** J

**Protocol Steps:**

1. Reads in input file as a data frame.
2. Splits data frame into two for each library.
3. The steps below are repeated for each library:

4. The number of samples above 2 ng/μL is calculated.
5. The concentration of each DNA sample required is calculated by dividing 1 by the number of samples above 2 ng/μL.
6. Samples between 2 ng/μL and the sample concentration required are calculated and will be added directly to the final library tube without dilution.
7. Samples that can not be diluted to 2x the concentration required in 10 μL (the sample concentration is between 1x and 2x) will be diluted to 1x concentration in 5 μL.
8. The individual volumes needed of all samples to be diluted to 2x concentration in 10 μL is calculated.
9. The P20 adds the calculated amount of ultrapure water to all the wells of samples to be diluted to 2x in the dilution plate.
10. The P20 adds the calculated amount of sample to all the corresponding wells in the dilution plate to be diluted to 2x and mixes 5x at 3x speed.
11. The P20 adds 1 μL of each sample diluted to 2x to the bottom of the first library tube.
12. The protocol pauses and asks user to remove the 2x library tube and vortex before loading back onto the OT-2 in the same position.
13. Half the total volume of the 2x library is transferred to the final library tube with the P20/P300 depending on the volume calculated (P300 if $>$20 μl or P20 if $\leq$ 19 μL).
14. The volumes of all samples to be diluted to 1x concentration in 5 μL is calculated.
15. The P20 adds the calculated amount of ultrapure water to all the wells of samples to be diluted to 1x in the dilution plate.
16. The P20 adds the calculated amount of sample to all the corresponding wells in the dilution plate to be diluted to 2x and mixes 5x at 3x speed.
17. The P20 adds the calculated amount of sample to all the corresponding wells in the dilution plate to be diluted to 1x and mixes 5x at 3x speed.
18. The P20 adds 1 μL of each sample diluted to 1x directly to the final library tube.
19. The volumes of the samples that can not be diluted are calculated by dividing the concentration needed by the concentration of the sample.
20. The P20 transfers the volume of each sample to be added neat directly the final library tube and mixes 5x.
21. The library tube is topped up to 47 μL either with ultrapure water or any samples below 2 ng/μL and with ultrapure water if still below 47 μL.
22. Writes an output file with all calculations for each sample to '/data/user_storage/output_files' called 'DATE_Flow_Cell_NUMBER_Library_Preparation_Calculations.csv'.

#### Supplementary Note 2: Opentrons OT-2 protocol optimization

#### Issues with evaporation during PCR steps

When establishing the PCR steps in our workflow, we initially faced issues with evaporation. During the first testing of the tailing PCR step with the OT-2 Thermocycler, which made use of a sterilized silicone reusable seal, we noticed around three quarters of the wells in the PCR plate were almost empty due to evaporation. After checking whether the plate type might have not fitted the seal correctly, we then tested two other different types of plates: (i) a 0.1 mL PCR plate recommended by Opentrons; and (ii) a 0.2 mL PCR plate from StarLab. Both types of plate were filled with 25 µL of ultrapure water and loaded into the thermocycler, pre-heated to 94°C, and held at this temperature for 30 min. We found that the 0.2 mL plate from StarLab had fewer wells affected by evaporation. However, both plates still showed some evaporation. Specifically, the PCR plate recommended by Opentrons had high evaporation levels ($<$10 µL of total volume remaining) for most of the wells and particularly those wells located close to the edges of the plate. The PCR plate from StarLab had less overall evaporation, but showed a higher evaporation level in the wells located near the plate edge.

To try an mitigate this issue, we tested whether an additional seal could help reduce evaporation. As the former tests were performed using only the sterilized, silicone, re-usable seal recommended for use with the OT-2 Thermocycler Module, we wanted to see whether addition of seal would help maintain a uniform volume inside the wells during the PCR reaction. Since most evaporation happened around the edges of the plate, we filled only wells located around the edges of two Starlab plates. We tested them, using the same conditions as before, the first sealed with a polyolefin seal and the second one with an aluminium foil seal. Overall, the polyolefin seal performed well, displaying very little evaporation (1–3 µL per well), while the foil seal remained stuck to the module and ripped off when the thermocycler opened. We therefore included an additional step of manually sealing the PCR plate with a polyofelin seal in the final workflow.

As slight evaporation in wells around the plate edge could not be completely avoided, we wanted to remove any uncertainty in sample volumes this might cause due to well-to-well differences. These could affect the bead-to-sample ratios during purification and hamper these purification steps. To overcome this issue, rather than using the entire contents of each well, we instead took a smaller set volume of sample, ensuring this was present in every well on the plate. Specifically, we set the robot to pick up 5 µL below the the initial PCR reaction volume such as from 25 µL and 45 µL to 20 µL and 40 µL for the tailing and barcoding PCR reactions, respectively. Working with uniform volumes of samples ensured consistency across the plate and removed the need to perform multiple calculations and change the pipette settings. As these steps leave some of the PCR products in the original plate, if needed, this material can be loaded onto an agarose gel or used for other downstream analysis (e.g., to troubleshoot problems).

#### Quantifluor Standards for DNA concentration quantification

For each DNA purification step performed post PCR, it was vital that accurate measurements of DNA concentration were made to allow for normalization of concentrations. This helped reduce biases between samples and ensured even sequencing of the barcoded samples. To quantify the DNA concentration, we chose to use the Quantifluor dsDNS ONE kit, which allows for the quantification of the concentration of small DNA samples using a plate reader with high accuracy and sensitivity in the range of 0.2–400 ng/µL. After measurements, a standard curve is generated using the Quantifluor dsDNS ONE system Excel analysis workbook using a linear regression. We evaluated the accuracy of the measurements using the coefficient of determination ($R^2$). Sample concentrations are interpolated from the standard curve automatically within the same workbook. To generate data for the standard curve, we automated the creation of the dilution series to reduce possible user error (Protocol 4). Initially, even when we used fresh standards prepared according to manufacturer instructions, we were only able to achieve an $R^2$ average of 0.96436. As fluorometric assays rely on reagents that are photosensitive, we optimized the protocol to minimize exposure of reagents to the light. We found that running the protocol in low light conditions and vortexing the plate immediately prior to use increased the average $R^2$ from 0.96436 to 0.9967.

### Supplementary Note 3: Running the experimental workflow manually

To carry out the automated part of the experimental workflow manually, we recommend following the PCR Barcoding Expansion 1–96 protocol (EXP-PBC096) from Oxford Nanopore Technologies to barcode the tailed PCR amplicons, and then performing the standard ligation sequencing protocol (SQK-LSK114) to create final library that are compatible with nanopore sequencing. For all clean up steps a ratio of 1:0.4 of sample to beads should be used.
