## Supplementary Data 1 for "Characterization of recombinase-based genetic parts and circuits using nanopore sequencing": CUSTOMLABWARE.md

# Custom Labware Dictionary 
All defined labware in the Opentrons module can be found [here](https://labware.opentrons.com) and custom labware definitions we have in the repository are listed below.

 Labware Type | Brand | Order Code | API Name
 ------------ | ----- | ---------- | --------
 24 Tube Aluminum Block | StarLab | S1615-5500-C | starlab_24_aluminumblock_1500ul
 24 Tubes in Rack | StarLab | S1615-550-C | starlab_24_tuberack_1500ml
 24 Tubes in Rack | Sarstedt | 72.690.004 | sarstedt_tuberack_1500ul
 24 Tubes in Rack | Eppendorf | 0030 108.051 | eppendorf_lo_bind_24_tuberack_1500ul
 24 Deep Well Plate | StarLab | E2824-0000 | starlab_24_wellplate_10000ul
 96 PCR Plate | StarLab | E1403-5200 | starlab_96_wellplate_200ul_pcr
 96 PCR Tube Strips in Rack | Tubes: Starlab, Rack: BioRad | Tubes: A1402-3700, Rack: TRC9601 | biorad_96_tuberack_200ul
 96 PCR Tube Strips in Aluminum Block | StarLab | A1402-3700 | starlab_strips_96_aluminumblock_200ul
 96 Deep Well Plate | StarLab | E2896-1810 | starlab_96_wellplate_2200ul
 96 Well Plate | Costar | 3603 | costar_96_wellplate_360ul
 96 Well Plate | Greiner | 655087 | greiner_96_wellplate_340ul
 96 Well Plate | Thermo Fisher | 249935 | thermo_96_wellplate_250ul
 96 Tubes in Rack | Thermofisher (Nanopore Barcodes) | 4273 | nanopore_96_tuberack_750ul
 384 Well Plate | Greiner | 781086 | greiner_384_wellplate_110ul
