## Supplementary Data 1 for "Characterization of recombinase-based genetic parts and circuits using nanopore sequencing": README.md

# labware
Contains all custom labware set up on the OT-2 with a list of the API names.

## Creating Custom Labware:
Go to the custom labware creator on the Opentrons [website](https://labware.opentrons.com/create) and follow their instructions. Be sure to use callipers to get accurate measurements and test the calibration with the test protocol you download at the same time as the file. Instructions on testing labware files are [here](https://opentrons-publications.s3.us-east-2.amazonaws.com/labwareDefinition_testGuide.pdf).

Once you have tested that the labware file you created is accurate. Move it to the custon labware folder for use and record new API name in CUSTOMLABWARE.md.
