## Supplementary Data 1 for "Characterization of recombinase-based genetic parts and circuits using nanopore sequencing": 1_Prep_Tailing_PCR_Mastermix.py

```
###############################################################################
# Author: Sarah Cameron, Biocompute Lab <>
# Last Updated: 14/03/22
###############################################################################
# Preparation of Tailing PCR Mastermix into Reservoir for one set of 96 Samples
# through the Automated Memory Readout Pipeline.
###############################################################################
# Protocol Length: 7 minutes
# Pipettes Required: P300 & P1000 Single Gen2
# Modules Required: N/A 
# Labware Required: NEST 12-Channel Reservoir (999-00076), Opentrons 4-in-1 
# 					Rack with 24-Tube insert (999-00050), 5x StarLab 1.5ml
# 					Microcentrifuge Tubes (S1615-5500), 1x p300 Opentrons Tip
# 				    Rack (999-00015), 1x p1000 Opentrons Tip Rack (999-00016). 
# Reagents Required: Nuclease Free Water, LongAmp Taq 2X Master Mix (M0287L), 
# 					 Forward and Reverse Primers for the PCR reaction. 
# Notes Before Starting: Load 170ul of each of the primers into an 1.5ml tube, 
#                        800ul of Taq into one tube and 700ul into two tubes.
# 						 Place all tubes into tube rack. A1 = Fwd Primer, 
#                        B1 = Rev Primer, A2 = Taq (800ul), 
# 						 B2 & C2 = Taq (700ul). In the reservoir add 3mls of 
#                        nuclease free water in column 12. The mastermix will 
#                        be made in column 1. See deck layout 1. 
###############################################################################

from opentrons import protocol_api
from datetime import date
import time
import sys
sys.path.append('/data/user_storage/modules/')
import video, log

metadata = {
	'protocolName': 'Prep_Tailing_PCR_Mastermix',
	'author': 'Sarah Cameron <>',
	'description': 'Creation of Mastermix in Bulk for Tailing PCR',
	'apiLevel': '2.11'
}

def run(protocol: protocol_api.ProtocolContext):
	#User defined constants
	USER = 'Sarah'
	PROTOCOL_NAME = 'PCR_Prep_Mastermix'
	DATE = date.today()
	WRITE_TO_LOG = True
	VIDEO = True
	VIDEO_LENGTH = '00:05:00'
	MODULES = 'NA'

	P300 = 'p300_single_gen2'
	P300_MOUNT = 'right'

	P1000 = 'p1000_single_gen2'
	P1000_MOUNT = 'left'

	#Load in Labware
	tiprack_1000_1 = protocol.load_labware('opentrons_96_tiprack_1000ul', '6')
	p1000_tipracks = [tiprack_1000_1]

	tiprack_300_1 = protocol.load_labware('opentrons_96_tiprack_300ul', '3')
	p300_tipracks = [tiprack_300_1]

	reagents = protocol.load_labware('starlab_24_tuberack_1500ml', '1')
	Primer_Fwd = reagents.wells_by_name()['A1'] #170 ul needed
	Primer_Rv = reagents.wells_by_name()['B1'] #170 ul needed
	Taq_1 = reagents.wells_by_name()['A2'] #800 ul needed 
	Taq_2 = reagents.wells_by_name()['B2'] #700 ul needed 
	Taq_3 = reagents.wells_by_name()['C2'] #700 ul needed 
	reservoir = protocol.load_labware('nest_12_reservoir_15ml', '2') 
	MM = reservoir.wells_by_name()['A1'] #Mastermix
	NFW = reservoir.wells_by_name()['A12'] #Nuclease Free Water

	#Load in Pipettes
	p1000 = protocol.load_instrument(P1000, P1000_MOUNT, tip_racks= p1000_tipracks)
	p300 = protocol.load_instrument(P300, P300_MOUNT, tip_racks= p300_tipracks)
	
	#Get ready to start protocol and ensure temp block set to 10C
	protocol.set_rail_lights(True) #Turn on lights
	start_time = time.time()

    #If the protocol is not simulating take a video of the protocol
	if not protocol.is_simulating() & VIDEO == True:
		video.take_video(VIDEO_LENGTH, PROTOCOL_NAME, USER, DATE, start_time)

	def adv_transfer(pipette, source, destination, move_to_speed, 
		transfer_volume, asp_flow_rate, asp_delay, tip_withdraw, 
		disp_flow_rate, disp_delay, mix_rep, mix_vol, mix_rate, blowout,
    	blow_flow, default_bf, touchtip, radius):
		'''A single transfer step that allows specification of advanced
		liquid handling settings not available in .transfer()'''
		
		#Move to top of liquid 
		pipette.pick_up_tip()
		pipette.move_to(source.top())

 		#Move to bottom of the well
		pipette.default_speed = move_to_speed 
		protocol.comment(f'Move To Speed = {move_to_speed}')

    	#Aspirate at 1/10th default flow rate for that pipette
		pipette.aspirate(transfer_volume, source.bottom(), rate = asp_flow_rate)

    	#Set the default speed back
		pipette.default_speed = 400

    	#Add delay after aspiration 
		protocol.delay(seconds=asp_delay)

    	#Withdraw tip to top of well at 1mm/s
		pipette.move_to(source.top(), speed = tip_withdraw)
		protocol.comment(f'Tip Withdrawal Speed - {tip_withdraw}')

    	#Dispense Liquid
    	#Move to top of well in the plate
		pipette.move_to(destination.top())

   	 	#Move to bottom of the labware for glycated liquids
		pipette.move_to(destination.bottom(),speed=move_to_speed)

    	#Dispense liquid at 1/10th setting for that pipette
    	#Surfactant liquids may require multiple dispense cycles
		pipette.dispense(transfer_volume, destination, rate=disp_flow_rate)

    	#Add post-dispense delay 
    	#Surfactant liquids and oils may require longer delay
		protocol.delay(seconds=disp_delay)
		
		pipette.mix(mix_rep, mix_vol, destination, rate = mix_rate)

    	#Blowout remaining volume from tip in liquid for glycated liquids 
    	#All others viscous liquid types remain at the side and top of the well 
		if blowout == True:
			pipette.flow_rate.blow_out = blow_flow
			protocol.comment(f'Blowing Out at {blow_flow} ul/s')
			pipette.blow_out()
			pipette.flow_rate.blow_out = default_bf

    	#Withdraw tip to top of the labware 
		pipette.move_to(destination.top(),speed=tip_withdraw)
		
		if touchtip == True: 
			pipette.touch_tip(radius=radius)
			protocol.comment(f'Touching Tip at {radius} Radius')
			
		pipette.drop_tip()

	#Define function for adding primers:
	def add_primers(source):
		'''Function containing transfer steps to take for each addition of 
		primers'''
		p300.pick_up_tip()
		p300.mix(3, 140, source, rate = 1.5)
		p300.aspirate(149, source)
		p300.dispense(149, MM)
		p300.mix(5, 300, MM, rate = 2)
		p300.blow_out()
		p300.drop_tip()

	#Protocol Commands
	protocol.comment('Add Nuclease Free Water')
	p1000.pick_up_tip()
	p1000.aspirate(800, NFW, rate=0.8)
	p1000.dispense(800, MM)
	p1000.aspirate(466.5, NFW)
	p1000.dispense(466.5, MM)
	p1000.drop_tip()
	
	protocol.comment('Add Forward Primers')
	add_primers(Primer_Fwd)
	
	protocol.comment('Add Reverse Primers')
	add_primers(Primer_Rv)
	
	protocol.comment('Add Taq Polymerase')
	adv_transfer(p1000, Taq_1, MM, 20, 
		662.5, 0.75, 10, 5, 1, 2, 10, 1000, 2, False, 100, 274.7, False, 0)

	adv_transfer(p1000, Taq_2, MM, 20, 
		600, 0.75, 10, 5, 1, 2, 10, 1000, 2, False, 100, 274.7, False, 0)

	adv_transfer(p1000, Taq_3, MM, 20, 
		600, 0.75, 10, 5, 1, 2, 10, 1000, 2, False, 100, 274.7, False, 0)

	#Turn lights off 
	protocol.set_rail_lights(False)

	#Write details of OT-2 use to log file 
	if not protocol.is_simulating() & WRITE_TO_LOG == True:
		log.log_record(start_time, P300_MOUNT, P300, P1000, DATE,
         USER, PROTOCOL_NAME, VIDEO, MODULES)
```
