## Supplementary Data 1 for "Characterization of recombinase-based genetic parts and circuits using nanopore sequencing": 2_PCR_Tailing_Reaction.py

```
###############################################################################
# Author: Sarah Cameron, Biocompute Lab <>
# Last Updated: 14/03/22
###############################################################################
# Preparation and execution of Tailing PCR reaction for High-Throughput 
# Memory Readout Pipeline.
###############################################################################
# Protocol Length: 3 hours 32 minutes
# Pipettes Required: P20 Multi Gen2 and P300 Multi Gen 2
# Modules Required: Thermocycler and Temperature Module 
# Labware Required: 12x 0.2ml StarLab PCR Strips (A1402-3700), 2x 0.2ml StarLab
#                   PCR plates (E1403-5200), 1x 0.1ml NEST PCR plate 
#                   (999-00050), Opentrons PCR Tube Aluminium Block, NEST 12-
#                   Channel Reservoir (999-00016), 2x Opentrons 20ul Tips 
#                   (999-00014), 1x Opentrons 300ul Tips (999-00015). 
#                   2x BioRad PCR Tube Racks (TRC9601), 1x StarLab Polyolefin 
#                   StarSeal (E2796-9793). 
# Reagents Required: Mastermix from PCR_Prep_Mastermix protocol in column 1 of 
#                    reservoir, Samples (25ul) in PCR Tube Strips on PCR Rack. 
# Notes Before Starting: Fill PCR aluminium block wells with 20ul of MilliQ 
#                        water. Set the temperature of the Temperature block 
#                        to 95C in Opentrons App 10 mins before starting the 
#                        protocol. Load Samples in rack to the aluminium block
#                        and start.  
###############################################################################

from opentrons import protocol_api
from datetime import date
import time
import sys
sys.path.append('/data/user_storage/modules/')
import video, log

metadata = {
	'protocolName': 'PCR_Tailing_Reaction',
	'author': 'Sarah',
	'description': 'Preparation and execution of tailing PCR reaction',
	'apiLevel': '2.11'
}

def run(protocol: protocol_api.ProtocolContext):
	#User defined constants
	USER = 'Sarah'
	PROTOCOL_NAME = 'PCR_Tailing_Reaction'
	DATE = date.today()
	WRITE_TO_LOG = True
	VIDEO = False
	VIDEO_LENGTH = '00:05:00'
	MODULES = ['thermocycler module', 'temperature module gen2']

	M20 = 'p20_multi_gen2'
	M20_MOUNT = 'left'

	M300 = 'p300_multi_gen2'
	M300_MOUNT = 'right'

	#Load in Modules
	tc_module = protocol.load_module(MODULES[0])
	temp_module = protocol.load_module(MODULES[1], '1')

	#Load in Labware
	tiprack_20_1 = protocol.load_labware('opentrons_96_tiprack_20ul', '3')
	tiprack_20_2 = protocol.load_labware('opentrons_96_tiprack_20ul', '6')
	m20_tipracks = [tiprack_20_1, tiprack_20_2]

	tiprack_300_1 = protocol.load_labware('opentrons_96_tiprack_300ul', '9')
	m300_tipracks = [tiprack_300_1]

	pcr_plate = tc_module.load_labware('starlab_96_wellplate_200ul_pcr', 
		label='PCR Plate')
	pcr_tube_block = temp_module.load_labware('starlab_strips_96_aluminumblock_200ul')
	storage_plate = protocol.load_labware('nest_96_wellplate_100ul_pcr_full_skirt', '4')
	end_plate = protocol.load_labware('starlab_96_wellplate_200ul_pcr', '5',
		label='End Plate')
	reservoir = protocol.load_labware('nest_12_reservoir_15ml', '2') 
	MM = reservoir.wells_by_name()['A1'] #Mastermix

	#Load in Pipettes
	m20 = protocol.load_instrument(M20, M20_MOUNT, tip_racks= m20_tipracks)
	m300 = protocol.load_instrument(M300, M300_MOUNT, tip_racks= m300_tipracks)
	
	#Get ready to start protocol and ensure thermocycler set to 95C
	protocol.set_rail_lights(True) #Turn on lights
	start_time = time.time()
	
    #If the protocol is not simulating take a video of the protocol
	if not protocol.is_simulating() & VIDEO == True:
		video.take_video(VIDEO_LENGTH, PROTOCOL_NAME, USER, DATE, start_time)

	#Protocol Commands
	temp_module.set_temperature(95)
	protocol.comment('Hold Samples at 95C for 10 mins')
	protocol.delay(minutes = 10)
	temp_module.deactivate()
	protocol.comment('''Remove Samples and Centrifuge for 2 mins at 4,000 rpm. 
		Once Complete, Load Tube Rack Back (Including any Samples prepared by 
		MiniPrep to Temp Module''')
	temp_module.set_temperature(20)

	protocol.pause('''Load Tube Rack to Temp Module before Resuming''')

	protocol.comment('''Transfer Mastermix to PCR Plate in Thermocycler''')
	tc_module.open_lid()

	columns = ['A1', 'A2', 'A3', 'A4', 'A5', 'A6', 'A7',
		 'A8', 'A9', 'A10', 'A11', 'A12']

	m300.pick_up_tip()
	m300.mix(10, 300, MM, rate = 1.5)
	for column in columns:
		m300.aspirate(23, MM, rate=0.75)
		m300.dispense(23, pcr_plate.wells_by_name()[column].bottom(0.25), rate=1) 
	m300.drop_tip()

	protocol.comment('''Transfer 10ul of Supernatant to Fresh PCR Storage
	 Plate and Add 2ul of Sample to Mastermix''')

	temp_module.deactivate()
	
	for column in columns:
		m20.pick_up_tip()
		m20.well_bottom_clearance.aspirate = 2 #Needs to not touch pellet (mm)
		m20.aspirate(12, pcr_tube_block.wells_by_name()[column], rate = 0.75)
		m20.well_bottom_clearance.aspirate = 1
		m20.dispense(10, storage_plate.wells_by_name()[column].bottom(), rate = 1) 
		m20.dispense(2, pcr_plate.wells_by_name()[column]) 
		m20.mix(5, 20, pcr_plate.wells_by_name()[column], rate=2)
		m20.drop_tip()

	protocol.pause('''Apply Thermocyler Seal to Plate and Remove Sample 
		Storage Plate to Store at 4C''')

	protocol.comment('Start PCR Reaction')
	tc_module.close_lid()
	tc_module.set_lid_temperature(105)
	tc_module.set_block_temperature(94, hold_time_minutes = 5, block_max_volume = 25)
	protocol.comment(f'Starting Cycle of {30} Repetitions')

	profile = [
		{'temperature': 94, 'hold_time_seconds': 20},
		{'temperature': 60, 'hold_time_seconds': 15},
		{'temperature': 65, 'hold_time_minutes': 3, 'hold_time_seconds':16}
	]

	tc_module.execute_profile(steps = profile, repetitions = 30, block_max_volume = 25)

	protocol.comment('Cycling Complete, Holding at 65C for 10 mins')
	tc_module.set_block_temperature(65, hold_time_minutes = 10, block_max_volume = 25)
	
	protocol.comment('PCR Reaction Complete, Holding Samples at 20C')
	tc_module.deactivate_lid()
	tc_module.set_block_temperature(20)
	tc_module.open_lid()

	protocol.pause('Remove Thermocycler Seal')

	protocol.comment('Move Reactions to Clean Plate')
	for column in columns:
		m20.pick_up_tip()
		m20.aspirate(20, pcr_plate.wells_by_name()[column].bottom(0.1), rate = 0.5)
		m20.dispense(20, end_plate.wells_by_name()[column], rate = 1) 
		m20.drop_tip()

	#Turn lights off 
	protocol.set_rail_lights(False)
	tc_module.deactivate_block()

	#Write details of OT-2 use to log file 
	if not protocol.is_simulating() & WRITE_TO_LOG == True:
		log.log_record(start_time, M20_MOUNT, M20, 'NA', DATE,
         USER, PROTOCOL_NAME, VIDEO, MODULES)
```
