## Supplementary Data 1 for "Characterization of recombinase-based genetic parts and circuits using nanopore sequencing": 3_Pur_DNA_Mag_20ul.py

```
###############################################################################
# Author: Sarah Cameron, Biocompute Lab <>
# Last Updated: 14/03/22
###############################################################################
# DNA purificiation protocol of 20ul PCR reaction using magnetic beads and
# magnetic module. 
###############################################################################
# Protocol Length: 1 hour 51 minutes 
# Pipettes Required: M20 and M300 Multi-Channel Gen 2 Pipettes.
# Modules Required: Magnetic Module Gen 2
# Labware Required: 3x 0.2ml StarLab 96 PCR Plates (E1403-5200), 1x NEST 
#                   12-Channel reservoir (999-00016), 7x 20ul Tip Racks 
#                   (999-00014), 3x 300ul Tip Racks (999-00015).
# Reagents Required: KAPA Pure Magnetic Beads (KK8002), 80% Ethanol (freshly 
#                    prepared), Nuclease Free Water.
# Notes Before Starting: Load 105ul of magnetic beads (room temperature, 
#                        vortexed) into each well of column 1 in one of the
#                        0.2ml PCR plates. Freshly prepare 40mls of 80% ethanol
#                        and load into columns 1 to 4 of reservoir. Add 2mls of
#                        NFW into column 5 of reservoir. See deck layout 3. 
###############################################################################

from opentrons import protocol_api
from opentrons import types
from datetime import date
import time
import sys
sys.path.append('/data/user_storage/modules/')
import video, log

metadata = {
	'protocolName': 'Pur_DNA_Mag_20ul',
	'author': 'Sarah Cameron <>',
	'description': 'Purification of DNA using Magnetic Beads',
	'apiLevel': '2.11'
}

def run(protocol: protocol_api.ProtocolContext):
	#User defined constants
	USER = 'Sarah'
	PROTOCOL_NAME = 'Pur_DNA_Mag_20ul'
	DATE = date.today()
	WRITE_TO_LOG = True
	VIDEO = True
	VIDEO_LENGTH = '00:10:00'
	MODULES = 'magnetic module gen2'

	M20 = 'p20_multi_gen2'
	M20_MOUNT = 'left'

	M300 = 'p300_multi_gen2'
	M300_MOUNT = 'right'
	
	BEAD_VOL = 8 #Reaction vol x 0.4 
	ENGAGE_HEIGHT = 8 #mm to engage magnet to
	ELUTION_VOL = 13 #ul of final elution product
	DNA_MIX_VOL = 10 #ul slowly mixed 5x with PCR rxs and beads
	BEAD_WASTE_1 = 15 #ul initally picked when removing rx waste 
	BEAD_WASTE_2 = 5 #ul picked when removing rx waste

	#Load in Modules
	mag_module = protocol.load_module(MODULES, '4')

	#Load in Labware
	tiprack_20_1 = protocol.load_labware('opentrons_96_tiprack_20ul', '3')
	tiprack_20_2 = protocol.load_labware('opentrons_96_tiprack_20ul', '6')
	tiprack_20_3 = protocol.load_labware('opentrons_96_tiprack_20ul', '9')
	tiprack_20_4 = protocol.load_labware('opentrons_96_tiprack_20ul', '11')
	m20_tipracks = [tiprack_20_1, tiprack_20_2, tiprack_20_3, tiprack_20_4]

	tiprack_300_1 = protocol.load_labware('opentrons_96_tiprack_300ul', '7')
	tiprack_300_2 = protocol.load_labware('opentrons_96_tiprack_300ul', '8')
	tiprack_300_3 = protocol.load_labware('opentrons_96_tiprack_300ul', '10')
	m300_tipracks = [tiprack_300_1, tiprack_300_2, tiprack_300_3]

	pcr_plate = mag_module.load_labware('starlab_96_wellplate_200ul_pcr', 
		label=-'PCR Plate')
	bead_plate = protocol.load_labware('starlab_96_wellplate_200ul_pcr', '2', 
		label='Magnetic Bead Plate') 
	beads = bead_plate.wells_by_name()['A1'] # 105ul in each well 
	elution_plate = protocol.load_labware('starlab_96_wellplate_200ul_pcr', '5', 
		label='Elution Plate')
	reservoir = protocol.load_labware('nest_12_reservoir_15ml', '1') 
	
	#col_1-4=ethanol, col_5=water, col_9-12=waste 
	ethanol_1 = reservoir.wells_by_name()['A1']
	ethanol_2 = reservoir.wells_by_name()['A2']
	ethanol_3 = reservoir.wells_by_name()['A3']
	ethanol_4 = reservoir.wells_by_name()['A4']
	water = reservoir.wells_by_name()['A5'] # Fill 2mls only
	waste_1 = reservoir.wells_by_name()['A8']
	waste_2 = reservoir.wells_by_name()['A9']
	waste_3 = reservoir.wells_by_name()['A10']
	waste_4 = reservoir.wells_by_name()['A11']
	waste_5 = reservoir.wells_by_name()['A12']
	
	#Load in Pipettes
	m20 = protocol.load_instrument(M20, M20_MOUNT, tip_racks= m20_tipracks)
	m300 = protocol.load_instrument(M300, M300_MOUNT, tip_racks= m300_tipracks)
	
	#Get ready to start protocol 
	protocol.set_rail_lights(True) #Turn on lights
	start_time = time.time()

    #If the protocol is not simulating take a video of the protocol
	if not protocol.is_simulating() & VIDEO == True:
		video.take_video(VIDEO_LENGTH, PROTOCOL_NAME, USER, DATE, start_time)
	
	odd_cols = ['A1', 'A3', 'A5', 'A7', 'A9', 'A11']

	#Define function for mixing at different heights:
	def mix_heights(HEIGHT, MIX_REPS, MIX_VOL, PLATE, RATE):
		'''Mix at different heights increase efficiency of mix'''
		m20.well_bottom_clearance.dispense = HEIGHT
		for x in range(MIX_REPS):
			m20.aspirate(MIX_VOL, PLATE, rate = RATE)
			m20.dispense(MIX_VOL, PLATE, rate = RATE)
		m20.well_bottom_clearance.dispense = 1

	#Define function for getting rid of ethanol drips: 
	def rm_waste_drips():
		'''Prevent contamination by shaking drips of ethanol off the end of
		pipette tips over the waste wells'''
		for x in range(3):
			m20.move_to(reservoir.wells_by_name()['A9'].top(), speed=400)
			m20.move_to(reservoir.wells_by_name()['A9'].top(10), speed=400)
			m20.move_to(reservoir.wells_by_name()['A12'].top(10), speed=400)
			m20.move_to(reservoir.wells_by_name()['A12'].top(), speed=400)
	
	#Define function for removing reaction supernatant:
	def rm_supernatant(waste_volume, location, bottom, first):
		'''Removing PCR Supernatant Slowly and in Multiple Transfers to
		 Prevent Picking Up the Bead Pellet'''
		m20.flow_rate.blow_out = 50
		m20.move_to(pcr_plate.wells_by_name()[column].top())
		m20.default_speed = 10
		if column in odd_cols:
			m20.aspirate(waste_volume, 
				pcr_plate.wells_by_name()[column].bottom(bottom).move(types.Point(-0.75,0,0)),
				rate = 0.5)
		else:
			m20.aspirate(waste_volume, 
				pcr_plate.wells_by_name()[column].bottom(bottom).move(types.Point(0.75,0,0)),
				rate = 0.5)
		m20.default_speed = 400
		m20.move_to(pcr_plate.wells_by_name()[column].top(), speed=5)
		m20.dispense(waste_volume+2, location, rate=1.75)
		m20.blow_out()
		if first == True:
			m20.air_gap(2)
		m20.flow_rate.blow_out = 7.6

	#Define function for adding ethanol: 
	def add_ethanol(half, ethanol_channel):
		'''Adding Ethanol to Half A Plate at A Time With Air Gap'''
		m300.pick_up_tip()
		for column in half:
			m300.aspirate(200, ethanol_channel, rate = 0.70)
			m300.air_gap(10)
			m300.dispense(215, pcr_plate.wells_by_name()[column].top(), rate = 0.5)
			m300.air_gap(5)
		m300.drop_tip()

	#Define function for removing ethanol supernatant:
	def rm_ethanol(half, waste_channel):
		'''Removing Ethanol from Half A Plate at A Time in Multiple Transfers
		With Different Tips to Prevent Disturbing Bead Pellet'''
		m20.flow_rate.blow_out = 50
		for column in half:
			protocol.comment(f'Remove Supernatant to Reservoir Channel: {waste_channel}')
			#m300
			m300.pick_up_tip()
			m300.well_bottom_clearance.aspirate = 3
			m300.move_to(pcr_plate.wells_by_name()[column].top())
			m300.default_speed = 10 
			m300.aspirate(180, pcr_plate.wells_by_name()[column], rate=0.75)
			m300.air_gap(5)
			m300.default_speed = 400
			m300.dispense(190, waste_channel)
			m300.drop_tip()
			#m20 
			m20.well_bottom_clearance.aspirate = 2
			m20.pick_up_tip()
			m20.move_to(pcr_plate.wells_by_name()[column].top())
			m20.air_gap(2)
			m20.aspirate(18, pcr_plate.wells_by_name()[column])
			m20.dispense(20, waste_channel.top(), rate=1.75)
			m20.blow_out()
			rm_waste_drips()
			m20.aspirate(20, pcr_plate.wells_by_name()[column].bottom(-0.25))
			m20.dispense(20, waste_channel)
			m20.drop_tip()
			m20.well_bottom_clearance.aspirate = 1
		m20.flow_rate.blow_out = 7.6

	#Protocol Commands
	protocol.pause('''Start with the PCR plate on the Elution plate 
		deck slot for initial mixing''')

	protocol.comment('Add Beads to All Wells in PCR Plate and Mix Thoroughly')

	columns = ['A1', 'A2', 'A3', 'A4', 'A5', 'A6', 'A7',
		 'A8', 'A9', 'A10', 'A11', 'A12']
	
	#Add beads from bead plate col 1 into PCR reactions on mag module
	for column in columns: 
		m20.pick_up_tip()
		m20.mix(3,BEAD_VOL, beads, rate = 1)
		m20.aspirate(BEAD_VOL, beads, rate = 0.75)
		m20.move_to(beads.top(), speed=2)
		m20.dispense(BEAD_VOL, elution_plate.wells_by_name()[column], rate = 0.75)
		mix_heights(1.5, 10, 15, elution_plate.wells_by_name()[column], 2.5)
		# MIX AGAIN 5x, DO THIS STEP SLOWLY SO BEADS CONTACT DNA
		m20.mix(5, DNA_MIX_VOL, elution_plate.wells_by_name()[column], rate = 1)
		m20.drop_tip() 

	#Incubate for 10 minutes
	protocol.comment('''Protocol will be delayed for 10 minutes to 
		allow for incubation.''')
	protocol.delay(minutes=10)
	
	protocol.home()
	protocol.pause('''Add PCR Plate to Magnetic Module and Put Clean 
		Elution Plate in its assigned Deck Slot. Also Empty Tip Waste''')

	#Engage magnet and incubate until liquid is clear
	protocol.comment('''Engaging Magnetic Module and Allowing Delay for
	 Beads to Pellet''')
	mag_module.engage(height = ENGAGE_HEIGHT) 
	protocol.delay(minutes=4) 
	
	#Discard PCR Supernatent to Waste Channel 1
	for column in columns:
		protocol.comment('Discarding Supernatant to Waste Channel 1')
		m20.pick_up_tip()
		rm_supernatant(BEAD_WASTE_1, waste_1.top(), 0, True)
		rm_supernatant(BEAD_WASTE_2, waste_1.bottom(), 0, False)
		m20.drop_tip()	

	#Add First Round of Ethanol to All Wells
	protocol.comment('Add Ethanol from Channels 1 & 2 to All Wells')
	half_cols = ['A1', 'A2', 'A3', 'A4', 'A5', 'A6']
	sec_half = ['A7', 'A8', 'A9', 'A10', 'A11', 'A12']
	
	add_ethanol(half_cols, ethanol_1)
	add_ethanol(sec_half, ethanol_2)
	
	#Incubate for ~40s if doing less than full plate
	#protocol.comment('Allow Pellet Formation')
	#protocol.delay(seconds=40)

	#Remove Ethanol Supernatant from First Half of Plate to Waste 2
	rm_ethanol(half_cols, waste_2)
	
	#Add Second Round of Ethanol to First Half of the Plate
	add_ethanol(half_cols, ethanol_3)

	#Remove Supernatant from Second Half of the Plate
	rm_ethanol(sec_half, waste_3)
	protocol.home()
	protocol.pause('''Please Empty Tip Waste Bin and Refill p20 Tip Racks.''')
	m20.reset_tipracks()
	
	#Add Second Round of Ethanol to Second Half of Plate to Waste 3
	add_ethanol(sec_half, ethanol_4)
	
	#Incubate for ~40s if doing less than a plate
	#protocol.delay(seconds=40)

	#Remove Second Ethanol Supernatant from First Half to Waste 4
	rm_ethanol(half_cols, waste_4)
	protocol.delay(minutes=2) #Allow time for drying

	#Add water to First Half to Keep from Drying Out 
	for col in half_cols:
		m20.transfer(ELUTION_VOL+3, water, pcr_plate.wells_by_name()[col], 
			new_tip='always')

	protocol.home()
	protocol.pause('Please Empty Tip Waste')

	#Remove Second Ethanol Supernatant from Second Half to Waste 5
	rm_ethanol(sec_half, waste_5)
	protocol.delay(minutes=2)  #Allow time for drying

	#Add water to Second Half to Keep from Drying Out 
	for sec in sec_half:
		m20.transfer(ELUTION_VOL+3, water, pcr_plate.wells_by_name()[sec], 
			new_tip='always')

	#Disengage magnet
	mag_module.disengage()

	#Go Back and Mix All Wells to Resuspend in Water
	for column in columns:
		m20.pick_up_tip()
		m20.mix(20, 16, pcr_plate.wells_by_name()[column].bottom(0.5), rate=10)
		m20.drop_tip()
	
	#Incubate for 2-10 mins 
	protocol.delay(minutes=10)  

	#Engage magnet
	mag_module.engage(height=ENGAGE_HEIGHT)
	protocol.delay(minutes=2)

	#Transfer sample to new plate
	protocol.comment('Transfer elution to new plate')
	for column in columns:
		m20.pick_up_tip()
		m20.move_to(pcr_plate.wells_by_name()[column].top())
		m20.default_speed = 10
		if column in odd_cols:
			m20.aspirate(ELUTION_VOL, 
				pcr_plate.wells_by_name()[column].bottom().move(types.Point(-0.75,0,0)),
				rate = 0.25)
		else:
			m20.aspirate(ELUTION_VOL,
				pcr_plate.wells_by_name()[column].bottom().move(types.Point(0.75,0,0)),
				rate = 0.25)
		m20.move_to(pcr_plate.wells_by_name()[column].top(), speed=5)
		m20.default_speed = 400
		m20.dispense(ELUTION_VOL, elution_plate.wells_by_name()[column])
		m20.drop_tip()
	
	#Turn lights off 
	protocol.set_rail_lights(False)

	#Write details of OT-2 use to log file 
	if not protocol.is_simulating() & WRITE_TO_LOG == True:
		log.log_record(start_time, M300_MOUNT, M300, M20, DATE,
         USER, PROTOCOL_NAME, VIDEO, MODULES)
```
