## Supplementary Data 1 for "Characterization of recombinase-based genetic parts and circuits using nanopore sequencing": 4_Prep_Quantifluor_Stds.py

```
###############################################################################
# Author: Sarah Cameron, Biocompute Lab <>
# Last Updated: 14/03/22
###############################################################################
# Preparation of QuantiFluor DNA Standards in PCR Plate to allow easy transfer
# by multi-channel to plate with samples for DNA Quantification using 
# Quantifluor ONE dsDNA system. 
###############################################################################
# Protocol Length: 7 minutes 
# Pipettes Required: P20 Single Gen2
# Modules Required: N/A 
# Labware Required: 1x NEST 0.1ml PCR Plate (999-00050), 2x 1.5ml LoBind 
#                   Eppendorf Tubes (0030 108.051), Opentrons Tube Rack with 
#                   24-Tube insert, 1x Opentrons 20ul Tip Rack (999-00014) 
# Reagents Required: dsDNA Standard, 1X TE Buffer from kit.
# Notes Before Starting: Load 50ul of dsDNA to one of the tubes and 150ul of 
#                        1X TE Buffer in the other. Load to rack A1 = TE Buffer
#                        C1 = dsDNA Standard. See Deck Layout 4. 
###############################################################################

from opentrons import protocol_api
from datetime import date
import time
import sys
sys.path.append('/data/user_storage/modules/')
import video, log

metadata = {
	'protocolName': 'Prep_Quantifluor_Stds',
	'author': 'Sarah Cameron <',
	'description': '''Preparing QuantiFluor dsDNA Standards''',
	'apiLevel': '2.11'
}

def run(protocol: protocol_api.ProtocolContext):
	#User defined constants
	USER = 'Sarah'
	PROTOCOL_NAME = 'Prep_Quantifluor_Stds'
	DATE = date.today()
	WRITE_TO_LOG = True
	VIDEO = True
	VIDEO_LENGTH = '00:05:00'
	MODULES = 'NA'
	
	P20 = 'p20_single_gen2'
	P20_MOUNT = 'left'

	M20 = 'NA'
	M20_MOUNT = 'right'

	COL_1 = 1

	#Load in Labware
	tiprack_20_1 = protocol.load_labware('opentrons_96_tiprack_20ul', '3')
	p20_tipracks = [tiprack_20_1]

	prep_tubes = protocol.load_labware('eppendorf_lo_bind_24_tuberack_1500ul', '2')
	stds_plate = protocol.load_labware('nest_96_wellplate_100ul_pcr_full_skirt', '1')

	#Label Reagents and Standards
	buffer = prep_tubes.wells_by_name()['A1'] # 150ul of TE Buffer
	dsDNA = prep_tubes.wells_by_name()['C1'].bottom() #Contains 50ul of Undiluted Standard

	#Load in Pipettes
	p20 = protocol.load_instrument(P20, P20_MOUNT, tip_racks= p20_tipracks)

	#Get ready to start protocol 
	protocol.set_rail_lights(True) 
	start_time = time.time()

    #If the protocol is not simulating take a video of the protocol
	if not protocol.is_simulating() & VIDEO == True:
		video.take_video(VIDEO_LENGTH, PROTOCOL_NAME, USER, DATE, start_time)

	#Protocol Commands
	def prep_stds(amount, source, destination):
		'''Dilute standard from the one before it in the series'''
		p20.pick_up_tip()
		p20.aspirate(amount, source)
		p20.move_to(source.top(), speed = 20)
		p20.dispense(amount, destination)
		p20.mix(10, 10, destination, rate = 2.5)
		p20.drop_tip()

	def prep_stds_set(std_col):
		std_A = stds_plate.wells_by_name()['A' + f'{std_col}'] 
		std_B = stds_plate.wells_by_name()['B' + f'{std_col}']
		std_C = stds_plate.wells_by_name()['C' + f'{std_col}']
		std_D = stds_plate.wells_by_name()['D' + f'{std_col}']
		std_E = stds_plate.wells_by_name()['E' + f'{std_col}']
		std_F = stds_plate.wells_by_name()['F' + f'{std_col}']
		std_G = stds_plate.wells_by_name()['G' + f'{std_col}']
		blank = stds_plate.wells_by_name()['H' + f'{std_col}']

		protocol.comment('Add Buffer to Wells')
		p20.pick_up_tip()
		p20.mix(3, 10, buffer)
		p20.transfer(10, buffer, std_B, new_tip = 'never')
		p20.transfer(15, buffer, std_C, new_tip = 'never')
		p20.transfer(15, buffer, std_D, new_tip = 'never')
		p20.transfer(15, buffer, std_E, new_tip = 'never')
		p20.transfer(15, buffer, std_F, new_tip = 'never')
		p20.transfer(15, buffer, std_G, new_tip = 'never')
		p20.transfer(15, buffer, blank, new_tip = 'never')
		p20.drop_tip()

		protocol.comment('Prepare Standard A')
		p20.transfer(20, dsDNA, std_A, mix_before = (5, 5))
		p20.transfer(10, dsDNA, std_A)

		protocol.comment('Prepare Standard B')
		prep_stds(10, std_A, std_B)

		protocol.comment('Prepare Standard C')
		prep_stds(5, std_B, std_C)

		protocol.comment('Prepare Standard D')
		prep_stds(5, std_C, std_D)

		protocol.comment('Prepare Standard E')
		prep_stds(5, std_D, std_E)

		protocol.comment('Prepare Standard F')
		prep_stds(5, std_E, std_F)

		protocol.comment('Prepare Standard G')
		prep_stds(5, std_F, std_G)

	prep_stds_set(COL_1)
	
	#Turn lights off 
	protocol.set_rail_lights(False)

	#Write details of OT-2 use to log file 
	if not protocol.is_simulating() & WRITE_TO_LOG == True:
		log.log_record(start_time, M20_MOUNT, M20, P20, DATE,
         USER, PROTOCOL_NAME, VIDEO, MODULES)
```
