## Supplementary Data 1 for "Characterization of recombinase-based genetic parts and circuits using nanopore sequencing": 5_Prep_Quantifluor_Samples.py

```
###############################################################################
# Author: Sarah Cameron, Biocompute Lab <>
# Last Updated: 14/03/22
###############################################################################
# Taking Purified DNA from Magnetic Clean Up Protocol and Preparing Plate of 
# Samples for dsDNA Quantification using Quantifluor ONE dsDNA System. Due to 
# the need to include the standard curves in duplicate the 96 samples will be
# read across 2 plates. This requires access to fluorescent plate reader 
# immediately after this protocol to measure fluorescence
# (Ex: 485nm, Em: 535nm).   
###############################################################################
# Protocol Length: 18 minutes 
# Pipettes Required: P20 and P300 Multi Gen2  
# Modules Required: N/A
# Labware Required: 1x NEST 12-Channel Reservoir (999-00076), 2x Greiner Black
#                   96 Well Plates (655087), 2x Opentrons 20ul Tips (999-00014)
#                   , 2x Opentrons 300ul Tips (999-00015). 1x 0.2ml StarLab 
#                   PCR plate (E1403-5200) with elutions from DNA Purification
#                   protocol. 1x 0.1ml PCR plate (999-00050) with Quantifluor 
#                   standards prepared in. 
# Reagents Required: QuantiFluor ONE dsDNA Dye 
# Notes Before Starting: Fill channel 1 and 2 in reservoir with 10.5mls of 
#                        Quantifluor Dye and channel 3 with 7.5ml. See deck
#                        layout 5. 
###############################################################################

from opentrons import protocol_api
from datetime import date
import time
import sys
sys.path.append('/data/user_storage/modules/')
import video, log

metadata = {
	'protocolName': 'Prep_Quantifluor_Samples',
	'author': 'Sarah Cameron',
	'description': '''Adding Samples and Standards to 200ul of QuantiFluor Dye
	 for Reading DNA Concentration''',
	'apiLevel': '2.11'
}

def run(protocol: protocol_api.ProtocolContext):
	#User defined constants
	USER = 'Sarah'
	PROTOCOL_NAME = 'Prep_Quantifluor_Samples'
	DATE = date.today()
	WRITE_TO_LOG = True
	VIDEO = True
	VIDEO_LENGTH = '00:05:00'
	MODULES = 'NA'

	QDYE_1 = 'A1'
	QDYE_2 = 'A2'
	QDYE_3 = 'A3'
	STDS_COL = 'A1'

	M20 = 'p20_multi_gen2'
	M20_MOUNT = 'left'

	M300 = 'p300_multi_gen2'
	M300_MOUNT = 'right'

	#Load in Labware
	tiprack_20_1 = protocol.load_labware('opentrons_96_tiprack_20ul', '3')
	tiprack_20_2 = protocol.load_labware('opentrons_96_tiprack_20ul', '6')
	tiprack_20_3 = protocol.load_labware('opentrons_96_tiprack_20ul', '9')
	m20_tipracks = [tiprack_20_1, tiprack_20_2, tiprack_20_3]

	tiprack_300_1 = protocol.load_labware('opentrons_96_tiprack_300ul', '10')
	tiprack_300_2 = protocol.load_labware('opentrons_96_tiprack_300ul', '11')
	m300_tipracks = [tiprack_300_1, tiprack_300_2]

	stds_plate = protocol.load_labware('nest_96_wellplate_100ul_pcr_full_skirt', '8')
	elution_plate = protocol.load_labware('starlab_96_wellplate_200ul_pcr', '2')
	reservoir = protocol.load_labware('nest_12_reservoir_15ml', '1') 
	reader_plate_1 = protocol.load_labware('greiner_96_wellplate_340ul', '4')
	reader_plate_2 = protocol.load_labware('greiner_96_wellplate_340ul', '5')

	standards = stds_plate.wells_by_name()[STDS_COL] # Column in plate with Standards in 
	quant_dye_1 = reservoir.wells_by_name()[QDYE_1] #QuantiFluor DNA Dye 
	quant_dye_2 = reservoir.wells_by_name()[QDYE_2] #QuantiFluor DNA Dye
	quant_dye_3 = reservoir.wells_by_name()[QDYE_3] #QuantiFluor DNA Dye

	#Load in Pipettes
	m20 = protocol.load_instrument(M20, M20_MOUNT, tip_racks= m20_tipracks)
	m300 = protocol.load_instrument(M300, M300_MOUNT, tip_racks= m300_tipracks)
	
	#Get ready to start protocol
	protocol.set_rail_lights(False) # Keep lights off, light sensitive protocol
	start_time = time.time()

    #If the protocol is not simulating take a video of the protocol
	if not protocol.is_simulating() & VIDEO == True:
		video.take_video(VIDEO_LENGTH, PROTOCOL_NAME, USER, DATE, start_time)

	#Define function to add standards and samples:
	def add_1ul(source, destination):
		'''Transfer step for adding 1ul of standard / sample to 
			Quantifluor Dye'''
		m20.pick_up_tip()
		m20.aspirate(1, source, rate=0.75)
		m20.dispense(1, destination, rate=1)
		m20.mix(3, 5, destination, rate = 3)
		m20.drop_tip()
	
	#Protocol Commands
	full_columns = ['A1', 'A2', 'A3', 'A4', 'A5', 'A6', 'A7',
		'A8', 'A9', 'A10', 'A11', 'A12']
	
	partial_columns = ['A1', 'A2', 'A3', 'A4', 'A5']
	
	protocol.comment('Add QuantiFluor Dye to All Wells')
	m300.pick_up_tip()
	for column in full_columns[:6]: 
		if column == full_columns[0]:
			m300.mix(3, 200, quant_dye_1, rate=0.75)
		m300.aspirate(200, quant_dye_1, rate=0.75)
		m300.move_to(quant_dye_1.top(), speed = 10)
		m300.dispense(200, reader_plate_1.wells_by_name()[column], rate=1)
	for column in full_columns[6:]: 
			m300.aspirate(200, quant_dye_2, rate=0.75)
			m300.move_to(quant_dye_2.top(), speed = 10)
			m300.dispense(200, reader_plate_1.wells_by_name()[column], rate=1)
	for column in partial_columns:
		m300.aspirate(200, quant_dye_3, rate=0.75)
		m300.move_to(quant_dye_3.top(), speed = 10)
		m300.dispense(200, reader_plate_2.wells_by_name()[column], rate=1)
	m300.drop_tip()

	protocol.comment('Add Standards')
	stds_reader_1 = ['A1', 'A12']
	stds_reader_2 = ['A1', 'A4']	

	#Mix Standards
	m20.pick_up_tip()
	m20.mix(5,5, standards.bottom(0.25), rate=2)
	m20.drop_tip()

	for stds in stds_reader_1:
		add_1ul(standards.bottom(), reader_plate_1.wells_by_name()[stds])

	for stds in stds_reader_2:
		add_1ul(standards.bottom(), reader_plate_2.wells_by_name()[stds])

	protocol.comment('Add Samples')
	sample_columns_r1 =  ['A2', 'A3', 'A4', 'A5', 'A6', 'A7', 'A8', 'A9', 'A10', 'A11']
	
	for sample in sample_columns_r1:
		add_1ul(elution_plate.wells_by_name()[sample], reader_plate_1.wells_by_name()[sample])

	add_1ul(elution_plate.wells_by_name()['A1'], reader_plate_2.wells_by_name()['A2'])
	add_1ul(elution_plate.wells_by_name()['A12'], reader_plate_2.wells_by_name()['A3'])

	protocol.comment('Mix Dye')
	for column in full_columns:
		m300.pick_up_tip()
		m300.mix(3, 100, reader_plate_1.wells_by_name()[column], rate = 1.5)
		m300.drop_tip()
	
	for column in partial_columns:
		m300.pick_up_tip()
		m300.mix(3, 100, reader_plate_2.wells_by_name()[column], rate = 1.5)
		m300.drop_tip()

	#Turn lights off 
	protocol.set_rail_lights(False)

	#Write details of OT-2 use to log file 
	if not protocol.is_simulating() & WRITE_TO_LOG == True:
		log.log_record(start_time, M300_MOUNT, M300, M20, DATE,
         USER, PROTOCOL_NAME, VIDEO, MODULES)
```
