## Supplementary Data 1 for "Characterization of recombinase-based genetic parts and circuits using nanopore sequencing": 6_Prep_Barcode_Dilution_Pt_1.py

```
###############################################################################
# Author: Sarah Cameron, Biocompute Lab <>
# Last Updated: 14/03/22
###############################################################################
# Take input file of DNA concentrations and prepare calculations for dilution 
# of all samples to 1.18ng/ul for Barcoding PCR. If any samples are below
# 20ng/ul they will be picked out of the elution plate in this protocol and 
# put into another plate to be diluted separately.  
###############################################################################
# Protocol Length: 5m + (Depending on how many samples below 20ng/ul)
# Pipettes Required: P20 Single Gen 2 
# Modules Required: N/A
# Labware Required: 1x 0.2ml StarLab PCR Plate with DNA samples in, 1x 0.1ml 
#                   NEST PCR Plate (), 1x Opentrons 20ul Tips ().
# Notes Before Starting: Load CSV on OT-2 with well IDs and sample
#                        concentration from Quantifluor analysis. See deck
#                        layout 6. 
# File Path: /data/user_storage/input_files/DATE_Elution_Concentrations.csv 
#            DATE format = YYYY-MM-DD
###############################################################################

from opentrons import protocol_api
import pandas as pd
from datetime import date
import time
import sys
sys.path.append('/data/user_storage/modules/')
import video, log

metadata = {
	'protocolName': 'Prep_Barcode_Dilution_Pt_1',
	'author': 'Sarah',
	'description': 'Calculating Volumes Needed for Dilution to 1.18 ng/ul',
	'apiLevel': '2.11'
}

def run(protocol: protocol_api.ProtocolContext):
    #User defined constants
    USER = 'Sarah'
    PROTOCOL_NAME = 'Prep_Barcode_Dilution_Pt_1'
    DATE = date.today()
    WRITE_TO_LOG = True
    VIDEO = True
    VIDEO_LENGTH = '00:05:00'
    MODULES = 'NA'
    
    M20 = 'p20_multi_gen2'
    M20_MOUNT = 'right'
    
    P20 = 'p20_single_gen2'
    P20_MOUNT = 'left'

    INPUT_PATH = '/data/user_storage/input_files/'
    OUTPUT_PATH = '/data/user_storage/output_files/'
    
    #Load in Labware
    tiprack_20_1 = protocol.load_labware('opentrons_96_tiprack_20ul', '3')
    p20_tipracks = [tiprack_20_1]
    
    elution_plate = protocol.load_labware('starlab_96_wellplate_200ul_pcr', '1')
    pick_plate = protocol.load_labware('nest_96_wellplate_100ul_pcr_full_skirt', '2')
	

	#Load in Pipettes
    p20 = protocol.load_instrument(P20, P20_MOUNT, tip_racks= p20_tipracks)
	
	#Get ready to start protocol
    protocol.set_rail_lights(True) #Turn on lights
    start_time = time.time()

    #If the protocol is not simulating take a video of the protocol
    if not protocol.is_simulating() & VIDEO == True:
        video.take_video(VIDEO_LENGTH, PROTOCOL_NAME, USER, DATE, start_time)

    #Create list of samples needed to be picked from the elution plate (samples 
    # that can't be diluted 1 in 7 in the next protocol)
    input_file = pd.read_csv(f'{INPUT_PATH}{DATE}_Elution_Concentrations.csv',
        header=None)

    df = pd.DataFrame(input_file)

    no_1in7 = df.loc[df[1] < 19.83]

    well_ids = no_1in7[0].tolist()

    protocol.comment('''Remove wells where DNA conc <20 ng/ul so they can be
     diluted separately''')
    for well in well_ids:
        p20.transfer(
			13,
			elution_plate.wells_by_name()[well].bottom(),
			pick_plate.wells_by_name()[well],
			new_tip = 'always')
    
    protocol.comment('Prepare Dilution Calculations')
    #Make CSV of list of wells sample volumes needed to dilute to 1.18 ng/ul
    barcode_dil = df.loc[df[1] >= 19.83]

    # Concentration After 1 in 7 Dilution concentration 
    barcode_dil[2] = round((barcode_dil[1]* 8) / 56, 2)
    
    # Volume of Sample Needed to Dilute to 1.18 ng/ul in 48ul
    barcode_dil[3] = round(56.64/barcode_dil[2], 1)

    file_path = f'{INPUT_PATH}{DATE}_Barcode_Dilutions.csv'
    barcode_dil.to_csv(file_path, header = False, index = False)

    # Make CSV of list of wells that are not possible to dilute to 1.18ng/ul 
    # They will just be made up to 24 ul with all of elution 
    no_dilution = no_1in7.loc[no_1in7[1] <= 5.66]
    if len(no_dilution) >= 16:
        protocol.comment('''Warning: Too Many Samples Below 1ng/ul, 
            Prepare Gel to Check Heat Lysis and PCR Worked''')

    if len(no_dilution) > 1:
        file_path = f'{INPUT_PATH}{DATE}_Below_Threshold_Dilutions.csv'
        no_dilution.to_csv(file_path, header = False, index = False)

    # Make CSV of list of wells sample volumes needed to dilute to 
    # 1.18 ng/ul for those not diluted 1 in 7 in 48ul 
    elute_dilute = no_1in7.loc[no_1in7[1] > 5.66]
    elute_dilute[2] = 'Dilute Directly to 1.18ng/ul'
    elute_dilute[3] = round(56.64/elute_dilute[1], 1)

    if len(elute_dilute) > 1:
        file_path = f'{INPUT_PATH}{DATE}_Elution_Dilutions.csv'
        elute_dilute.to_csv(file_path, header = False, index = False)

    #Make merged output file of all dilutions
    merges = [barcode_dil, no_dilution, elute_dilute]
    output_file = pd.concat(merges)
    output_file = output_file.sort_index()
    output_file.columns = ['Well ID', 'Elution Concentration (ng/ul)',
         'Concentration after 1 in 7 Dilution', 'Volume to Dilute By']

    file_path = f'{OUTPUT_PATH}{DATE}_Barcode_PCR_Preparation_Calculations.csv'
    output_file.to_csv(file_path, header = True, index = False)

    #Turn lights off
    protocol.set_rail_lights(False)
    
    #Write details of OT-2 use to log file 
    if not protocol.is_simulating() & WRITE_TO_LOG == True:
        log.log_record(start_time, P20_MOUNT, P20, M20, DATE,
         USER, PROTOCOL_NAME, VIDEO, MODULES)
```
