## Supplementary Data 1 for "Characterization of recombinase-based genetic parts and circuits using nanopore sequencing": 7_Prep_Barcode_Dilution_Pt_2.py

```
###############################################################################
# Author: Sarah Cameron, Biocompute Lab <>
# Last Updated: 14/03/22
###############################################################################
# Dilute all samples in elution plate 1 in 7. 
###############################################################################
# Protocol Length: 15 minutes 
# Pipettes Required: P20 and P300 Multi Gen2  
# Modules Required: N/A
# Labware Required: 1x NEST 12-Well Reservoir (999-00076), 1x 0.1ml PCR Plate
#                   (999-00050), 1x 0.2ml StarLab PCR Plate (E1403-5200) with 
#                    purified DNA samples. 
# Reagents Required: Nuclease Free Water
# Notes Before Starting: Load 6mls of NFW in Column 12 of the reservoir. See 
#                        deck layout 7.
###############################################################################

from opentrons import protocol_api
from datetime import date
import time
import sys
sys.path.append('/data/user_storage/modules/')
import video, log

metadata = {
	'protocolName': 'Prep_Barcode_Dilution_Pt_2',
	'author': 'Sarah Cameron <>',
	'description': 'Diluting Clean Up Reactions 1:7',
	'apiLevel': '2.11'
}

def run(protocol: protocol_api.ProtocolContext):
	#User defined constants
	USER = 'Sarah'
	PROTOCOL_NAME = 'Prep_Barcode_Dilution_Pt_2'
	DATE = date.today()
	WRITE_TO_LOG = True
	VIDEO = True
	VIDEO_LENGTH = '00:05:00'
	MODULES = 'NA'

	M20 = 'p20_multi_gen2'
	M20_MOUNT = 'left'

	M300 = 'p300_multi_gen2'
	M300_MOUNT = 'right'

	#Load in Labware
	tiprack_20_1 = protocol.load_labware('opentrons_96_tiprack_20ul', '3')
	m20_tipracks = [tiprack_20_1]

	tiprack_300_1 = protocol.load_labware('opentrons_96_tiprack_300ul', '6')
	m300_tipracks = [tiprack_300_1]

	elution_plate = protocol.load_labware('starlab_96_wellplate_200ul_pcr', '2') 
	dilute_plate = protocol.load_labware('nest_96_wellplate_100ul_pcr_full_skirt', '4')
	reservoir = protocol.load_labware('nest_12_reservoir_15ml', '1') 
	NFW = reservoir.wells_by_name()['A12'] #Fill 6mls

	#Load in Pipettes
	m20 = protocol.load_instrument(M20, M20_MOUNT, tip_racks= m20_tipracks)
	m300 = protocol.load_instrument(M300, M300_MOUNT, tip_racks= m300_tipracks)
	
	#Get ready to start protocol and ensure temp block set to 4C
	protocol.set_rail_lights(True) #Turn on lights
	start_time = time.time()

    #If the protocol is not simulating take a video of the protocol
	if not protocol.is_simulating() & VIDEO == True:
		video.take_video(VIDEO_LENGTH, PROTOCOL_NAME, USER, DATE, start_time)

	#Define function for mixing at different heights:
	def mix_heights(HEIGHT, MIX_REPS, MIX_VOL, PLATE, RATE):
		'''Function to mix at different heights'''
		m20.well_bottom_clearance.dispense = HEIGHT
		for x in range(MIX_REPS):
			m20.aspirate(MIX_VOL, PLATE, rate = RATE)
			m20.dispense(MIX_VOL, PLATE, rate = RATE)
		m20.well_bottom_clearance.dispense = 1

	#Protocol Commands
	columns = ['A1', 'A2', 'A3', 'A4', 'A5', 'A6', 'A7',
		'A8', 'A9', 'A10', 'A11', 'A12']
	

	protocol.comment('Add 48ul of Water to All Wells')
	
	m300.pick_up_tip()
	for column in columns:
		m300.aspirate(48, NFW, rate=1)
		m300.dispense(48, dilute_plate.wells_by_name()[column].bottom(), rate=1)
	m300.drop_tip()

	protocol.comment('Add 8ul of Sample to All Wells')
	
	for column in columns:
		m20.pick_up_tip()
		m20.aspirate(8, elution_plate.wells_by_name()[column], rate=0.5)
		m20.dispense(8, dilute_plate.wells_by_name()[column].bottom(), rate=0.5)
		mix_heights(5, 10, 20, dilute_plate.wells_by_name()[column], 2)
		m20.drop_tip()
	
	protocol.comment('''Plate Diluted 1:7, Please Change Pipettes to p20 and 
		p300 to Continue Dilution of Individual Samples''')

	#Turn lights off 
	protocol.set_rail_lights(False)

	#Write details of OT-2 use to log file 
	if not protocol.is_simulating() & WRITE_TO_LOG == True:
		log.log_record(start_time, M300_MOUNT, M300, M20, DATE,
         USER, PROTOCOL_NAME, VIDEO, MODULES)
```
