## Supplementary Data 1 for "Characterization of recombinase-based genetic parts and circuits using nanopore sequencing": 8_Prep_Barcode_Dilution_Pt_3.py

```
###############################################################################
# Author: Sarah Cameron, Biocompute Lab <>
# Last Updated: 14/03/22
###############################################################################
# Continue from Barcode Dilution Preparation protocol part 1 & 2. Dilute all 
# samples to 1.18ng/ul from known concentrations. 
###############################################################################
# Protocol Length: 1 hour 32 minutes
# Pipettes Required: P20 & P300 Single Gen2
# Modules Required: N/A 
# Labware Required: 1x NEST 12-Well Reservoir (999-00076), 3x 0.1ml PCR Plate
#                   (999-00050) - one is the pick plate from Part 1 of this 
#                   protocol, another is the 1 in 7 dilution plate from Part 2,
#                   2x Opentrons 20ul Tips (999-00014), 2x Opentrons 300ul Tips
#                  (999-00015)
# Reagents Required: Nuclease Free Water
# Notes Before Starting: Load 6mls of NFW in Column 12 of the reservoir. See
#                        deck layout 8. 
###############################################################################

from opentrons import protocol_api
from os.path import exists
import pandas as pd
from datetime import date
import time
import sys
sys.path.append('/data/user_storage/modules/')
import video, log

metadata = {
	'protocolName': 'Prep_Barcode_Dilution_Pt_3',
	'author': 'Sarah Cameron <>',
	'description': '''Diluting all Samples to 1.18ng/ul''',
	'apiLevel': '2.11'
}

def run(protocol: protocol_api.ProtocolContext):
	#User defined constants
	USER = 'Sarah'
	PROTOCOL_NAME = 'Prep_Barcode_Dilution_Pt_2'
	DATE = date.today()
	WRITE_TO_LOG = True
	VIDEO = True
	VIDEO_LENGTH = '00:05:00'
	MODULES = 'NA'
	
	P20 = 'p20_single_gen2'
	P20_MOUNT = 'left'

	P300 = 'p300_single_gen2'
	P300_MOUNT = 'right'

	INPUT_PATH = '/data/user_storage/input_files/'

	#Load in Labware
	tiprack_20_1 = protocol.load_labware('opentrons_96_tiprack_20ul', '3')
	tiprack_20_2 = protocol.load_labware('opentrons_96_tiprack_20ul', '6')
	p20_tipracks = [tiprack_20_1, tiprack_20_2]

	tiprack_300_1 = protocol.load_labware('opentrons_96_tiprack_300ul', '9')
	tiprack_300_2 = protocol.load_labware('opentrons_96_tiprack_300ul', '10')
	p300_tipracks = [tiprack_300_1, tiprack_300_2]

	dilute_plate = protocol.load_labware('nest_96_wellplate_100ul_pcr_full_skirt', '4',
		label='Diluted Elutions 1 in 7')
	final_dilute_plate = protocol.load_labware('nest_96_wellplate_100ul_pcr_full_skirt', '2', 
		label='Final Dilution Plate')
	pick_plate = protocol.load_labware('nest_96_wellplate_100ul_pcr_full_skirt', '5', 
		label='Pick Plate')

	reservoir = protocol.load_labware('nest_12_reservoir_15ml', '1') 
	NFW = reservoir.wells_by_name()['A11']

	#Load in Pipettes
	p20 = protocol.load_instrument(P20, P20_MOUNT, tip_racks= p20_tipracks)
	p300 = protocol.load_instrument(P300, P300_MOUNT, tip_racks= p300_tipracks)

	#Define mix heights function
	def mix_heights(HEIGHT, MIX_REPS, MIX_VOL, PLATE, RATE):
		'''Function to mix at different heights'''
		p20.well_bottom_clearance.dispense = HEIGHT
		for x in range(MIX_REPS):
			p20.aspirate(MIX_VOL, PLATE, rate = RATE)
			p20.dispense(MIX_VOL, PLATE, rate = RATE)
		p20.well_bottom_clearance.dispense = 1
	
	#Get ready to start protocol 
	protocol.set_rail_lights(True) #Turn on lights
	start_time = time.time()
	
    #If the protocol is not simulating take a video of the protocol
	if not protocol.is_simulating() & VIDEO == True:
		video.take_video(VIDEO_LENGTH, PROTOCOL_NAME, USER, DATE, start_time)

	#Protocol Commands
	#Read in CSV File with Sample Volumes Needed for Elutions diluted 1 in 7 
	barcode_dilutions = pd.read_csv(f'{INPUT_PATH}{DATE}_Barcode_Dilutions.csv',
	 header=None)
	df = pd.DataFrame(barcode_dilutions)
	sample_vols = df[3].tolist()
	well_ids = df[0].tolist()
	
	protocol.comment('Transfer water to PCR plate needed for final dilution') 
	p300.pick_up_tip()
	for well, volume in zip(well_ids, sample_vols): 
		p300.transfer(
			(48-volume), 
			NFW, 
			final_dilute_plate.wells_by_name()[well], 
			new_tip = 'never')
	p300.drop_tip()
		
	protocol.comment('Add volume of 1 in 7 diluted samples and dilute to 1.18ng/ul')
	for well, volume in zip(well_ids, sample_vols):
		p20.pick_up_tip()
		p20.transfer(
			volume,
			dilute_plate.wells_by_name()[well],
			final_dilute_plate.wells_by_name()[well],
			new_tip = 'never')
		mix_heights(3, 10, 20, final_dilute_plate.wells_by_name()[well], 3)
		p20.drop_tip()
	
	#Read in CSV with sample volumes needed for those samples not diluted
	elute_file_path = f'{INPUT_PATH}{DATE}_Elution_Dilutions.csv'
	if exists(elute_file_path) == True:
		elution_dilutions = pd.read_csv(elute_file_path, header=None)
		ed = pd.DataFrame(elution_dilutions)
		ed_well_ids = ed[0].tolist()
		ed_sample_vols = ed[3].tolist()

		p300.pick_up_tip()
		for well, volume in zip(ed_well_ids, ed_sample_vols):
			p300.transfer(
				(48-volume),
				NFW,
				final_dilute_plate.wells_by_name()[well],
				new_tip = 'never')
		p300.drop_tip()
	
		protocol.comment('Add volume of undiluted elutions to dilute to 1.18ng/ul')
		for well, volume in zip(ed_well_ids, ed_sample_vols):
			p20.pick_up_tip()
			p20.transfer(
				volume,
				pick_plate.wells_by_name()[well].bottom(),
				final_dilute_plate.wells_by_name()[well],
				new_tip = 'never')
			mix_heights(3, 10, 20, final_dilute_plate.wells_by_name()[well], 3)
			p20.drop_tip()

	
	#Read in CSV with wells where the samples cannot be diluted to 1.18ng/ul
	below_file_path = f'{INPUT_PATH}{DATE}_Below_Threshold_Dilutions.csv'
	if exists(below_file_path) == True:
		no_dilutions = pd.read_csv(below_file_path, header=None)
		nd = pd.DataFrame(no_dilutions)
		nd_well_ids = nd[0].tolist()
		
		p20.pick_up_tip()
		for well in nd_well_ids:
			p20.transfer(18, NFW, final_dilute_plate.wells_by_name()[well],
				new_tip='never')
		p20.drop_tip() 

		protocol.comment('Add 8ul of samples below threshold to 18ul of NFW')
		for well in nd_well_ids:
			p20.pick_up_tip()
			p20.transfer(8, pick_plate.wells_by_name()[well].bottom(),
				final_dilute_plate.wells_by_name()[well], 
				new_tip = 'never')
			mix_heights(3, 10, 20, final_dilute_plate.wells_by_name()[well], 3)
			p20.drop_tip()
	
	
	protocol.comment('''All samples prepared for Barcoding PCR''')
		
	#Turn lights off 
	protocol.set_rail_lights(False)

	#Write details of OT-2 use to log file 
	if not protocol.is_simulating() & WRITE_TO_LOG == True:
		log.log_record(start_time, P300_MOUNT, P300, P20, DATE,
         USER, PROTOCOL_NAME, VIDEO, MODULES)
```
