## Supplementary Data 1 for "Characterization of recombinase-based genetic parts and circuits using nanopore sequencing": 9_PCR_Barcoding_Reaction.py

```
###############################################################################
# Author: Sarah Cameron, Biocompute Lab <>
# Last Updated: 15/03/22
###############################################################################
# Preparation and execution of Nanopore Barcoding PCR Reaction for 
# High-Throughput Memory Readout Pipeline. 
###############################################################################
# Protocol Length: 1 hour 45 mins (2 hours 35 mins for Cascade Experiment)
# Pipettes Required: P20 & M300 Multi Gen2
# Modules Required: Thermocycler 
# Labware Required: 2x 0.2ml StarLab PCR plates (E1403-5200), 1x 0.1ml NEST PCR
#                   plate (999-00050) with DNA Samples Diluted to 1.18ng/ul, 
#                   1x NEST 12-Channel Reservoir (999-00016), 1x Opentrons 20ul
#                   Tips (999-00014), 3x Opentrons 300ul Tips (999-00015). 
#                   1x StarLab Polyolefin StarSeal (E2796-9793). 
# Reagents Required: LongAmp Taq Mastermix (M0287L), 
#                    Nanopore Barcoding Primers (EXP-PBC096).  
# Notes Before Starting: **If running Cascade version of the pipeline change
#                        'CASCADE_EXP' to True**.
#                        Spin Nanopore barcode plate at 4,000 rpm for 1 min to
#                        ensure all liquid at the bottom of the tubes. Load
#                        3.5mls of Taq Mastermix in Column 4 of reservoir. If 
#                        DNA samples have been left overnight vortex before 
#                        use. See deck layout 9. 
###############################################################################

from opentrons import protocol_api
from datetime import date
import time
import sys
sys.path.append('/data/user_storage/modules/')
import video, log

metadata = {
	'protocolName': 'PCR_Tailing_Reaction',
	'author': 'Sarah Cameron <>',
	'description': 'Preparation and execution of tailing PCR reaction',
	'apiLevel': '2.11'
}

def run(protocol: protocol_api.ProtocolContext):
	#User defined constants
	USER = 'Sarah'
	PROTOCOL_NAME = 'PCR_Tailing_Reaction'
	DATE = date.today()
	WRITE_TO_LOG = True
	VIDEO = False
	VIDEO_LENGTH = '00:05:00'
	MODULES = 'thermocycler module'

	M20 = 'p20_multi_gen2'
	M20_MOUNT = 'left'

	M300 = 'p300_multi_gen2'
	M300_MOUNT = 'right'


	CASCADE_EXP = False #Change to true if running cascade version of pipeline
	# This will increase the elongation time in the PCR to 8 minutes. 

	#Load in Modules
	tc_module = protocol.load_module(MODULES)

	#Load in Labware
	tiprack_20_1 = protocol.load_labware('opentrons_96_tiprack_20ul', '3')
	m20_tipracks = [tiprack_20_1]

	tiprack_300_1 = protocol.load_labware('opentrons_96_tiprack_300ul', '6')
	tiprack_300_2 = protocol.load_labware('opentrons_96_tiprack_300ul', '9')
	m300_tipracks = [tiprack_300_1, tiprack_300_2]


	pcr_plate = tc_module.load_labware('starlab_96_wellplate_200ul_pcr',
		label='PCR Plate - StarLab 0.2ml PCR')
	final_dilute_plate = protocol.load_labware('nest_96_wellplate_100ul_pcr_full_skirt', '4')
	barcode_primers = protocol.load_labware('nanopore_96_tuberack_750ul', '1')
	end_plate = protocol.load_labware('starlab_96_wellplate_200ul_pcr', '5', 
		label='End PCR Plate - StarLab 0.2ml PCR')
	reservoir = protocol.load_labware('nest_12_reservoir_15ml', '2')  
	MM = reservoir.wells_by_name()['A4']

	#Load in Pipettes
	m20 = protocol.load_instrument(M20, M20_MOUNT, tip_racks= m20_tipracks)
	m300 = protocol.load_instrument(M300, M300_MOUNT, tip_racks= m300_tipracks)

	#Get ready to start protocol 
	protocol.set_rail_lights(True) #Turn on lights
	start_time = time.time()

    #If the protocol is not simulating take a video of the protocol
	if not protocol.is_simulating() & VIDEO == True:
		video.take_video(VIDEO_LENGTH, PROTOCOL_NAME, USER, DATE, start_time)

	#Protocol Commands
	protocol.comment('Add Taq Polymerase Mastermix to PCR Plate')
	columns = ['A1', 'A2', 'A3', 'A4', 'A5', 'A6', 'A7',
		'A8', 'A9', 'A10', 'A11', 'A12']

	protocol.comment('Add PCR Master Mix to PCR Plate in Thermocycler')
	m300.pick_up_tip()
	for column in columns:
		m300.move_to(MM.top())
		m300.default_speed = 20
		m300.aspirate(25, MM.bottom(), rate=0.75)
		protocol.delay(seconds=2)
		m300.default_speed = 400
		m300.move_to(MM.top(), speed=10)
		m300.move_to(pcr_plate.wells_by_name()[column].top())
		m300.dispense(25, pcr_plate.wells_by_name()[column].bottom(), rate=1) 
		protocol.delay(seconds=1)
	m300.drop_tip()

	protocol.comment('Add Barcoding Primers to Master Mix')
	for column in columns:
		m20.pick_up_tip()
		m20.aspirate(1, barcode_primers.wells_by_name()[column])
		m20.dispense(1, pcr_plate.wells_by_name()[column].bottom()) 
		m20.mix(5, 15, pcr_plate.wells_by_name()[column], rate = 2)
		m20.drop_tip()

	protocol.comment('Add Samples to Master Mix')
	for column in columns:
		m300.pick_up_tip()
		m300.aspirate(24, final_dilute_plate.wells_by_name()[column].bottom(0.1))
		m300.dispense(24, pcr_plate.wells_by_name()[column].bottom()) 
		m300.mix(10, 40, pcr_plate.wells_by_name()[column].bottom(), rate = 1)
		m300.drop_tip()

	protocol.pause('Add Thermocycler Seal')

	protocol.comment('Start PCR Reaction')
	tc_module.close_lid()
	tc_module.set_lid_temperature(105)
	tc_module.set_block_temperature(95, hold_time_minutes = 3, block_max_volume = 50)
	
	protocol.comment(f'Starting Cycle of {12} Repetitions')

	profile = [
		{'temperature': 95, 'hold_time_seconds': 15},
		{'temperature': 62, 'hold_time_seconds': 15},
		{'temperature': 65, 'hold_time_minutes': 3, 'hold_time_seconds': 50}
	]

	cascade_profile = [
		{'temperature': 95, 'hold_time_seconds': 15},
		{'temperature': 62, 'hold_time_seconds': 15},
		{'temperature': 65, 'hold_time_minutes': 8}
	]
	
	if CASCADE_EXP == True:
		tc_module.execute_profile(steps = cascade_profile, repetitions = 12, 
			block_max_volume = 50)
	else:
		tc_module.execute_profile(steps = profile, repetitions = 12, 
			block_max_volume = 50)

	
	protocol.comment('Cycling Complete, Holding at 65C for 4 mins')
	tc_module.deactivate_lid()
	tc_module.set_block_temperature(65, hold_time_minutes = 4,
		block_max_volume = 50)
	
	protocol.comment('PCR Reaction Complete, Holding Samples at 10C')
	tc_module.set_block_temperature(10)
	tc_module.open_lid()

	protocol.pause('''Put Fresh 300ul Tip Rack on Slot 6 and
	 Remove Thermocycler Seal''')
	m300.reset_tipracks()

	protocol.comment('Move Reactions to Clean Plate')
	for column in columns:
		m300.pick_up_tip()
		m300.aspirate(40, pcr_plate.wells_by_name()[column], rate=0.75)
		m300.dispense(40, end_plate.wells_by_name()[column]) 
		m300.air_gap(5) #To stop drips and contamination
		m300.drop_tip()

	protocol.comment('Please Remove End Plate and Store at 4C')

	tc_module.deactivate_block()
	
	#Turn lights off 
	protocol.set_rail_lights(False)

	#Write details of OT-2 use to log file 
	if not protocol.is_simulating() & WRITE_TO_LOG == True:
		log.log_record(start_time, M20_MOUNT, M20, 'NA', DATE,
         USER, PROTOCOL_NAME, VIDEO, MODULES)
```
