## Supplementary Data 1 for "Characterization of recombinase-based genetic parts and circuits using nanopore sequencing": 11_Prep_DNA_Libraries.py

```
###############################################################################
# Author: Sarah Cameron, Biocompute Lab <>
# Last Updated: 15/03/2022
###############################################################################
# Dilute all samples to concentrations needed to add equal amount of DNA for
# Nanopore sequencng. To prevent pipetting outside of the p20 pipette range
# (below 1ul) samples will be diluted in 10ul to 2x amount needed / ul) and 1ul
# of each sample will be added to a tube, vortexed and half the total volume 
# added to a final library tube. Samples with a concentration between the amount
# of DNA needed and 2x the amount, will be diluted to 1x concentration in 5ul 
# and 1ul of each will be added to the final library tube. 
###############################################################################
# Protocol Length: 1 hour 45 minutes
# Pipettes Required: P20 & P300 Single Gen2
# Modules Required: N/A 
# Labware Required: 1x NEST 0.1ml PCR Plate (999-00050), 2x Eppendorf LoBind 
#                   1.5 ml Tubes (0030 108.051), 1x NEST 12-Channel Reservoir 
#                   (999-00076), 3x Opentrons P20 Tip Racks (999-00014), 1x 
#                   Opentrons P300 Tip Racks (999-00015).
# Reagents Required: Nuclease Free Water
# Notes Before Starting: Load CSV on OT-2 with well IDs and sample
#                        concentration from Quantifluor analysis.
#                        Fill channel 12 in reservoir with 1ml of NFW. See 
#                        deck layout 11. 
# File Path: 
# 	     /data/user_storage/input_files/DATE_Library_Elution_Concentrations.csv 
#            DATE format = YYYY-MM-DD
###############################################################################

from opentrons import protocol_api
import pandas as pd
from datetime import date
import time
import sys
sys.path.append('/data/user_storage/modules/')
import video, log

metadata = {
	'protocolName': 'Prep_DNA_Libraries',
	'author': 'Sarah Cameron <>',
	'description': '''Preparing Samples to Correct Concentration for Nanopore 
		Sequencing in 2 Flow Cells''',
	'apiLevel': '2.11'
}

def run(protocol: protocol_api.ProtocolContext):
	#User defined constants
	USER = 'Sarah'
	PROTOCOL_NAME = 'Prep_DNA_Flow_Cell'
	DATE = date.today()
	WRITE_TO_LOG = True
	VIDEO = True
	VIDEO_LENGTH = '00:05:00'
	MODULES = 'NA'
	
	P20 = 'p20_single_gen2'
	P20_MOUNT = 'left'

	P300 = 'p300_single_gen2'
	P300_MOUNT = 'right'

	NO_SAMPLES = 48 #Enter the number of samples in each half the plate 

	IN = '/data/user_storage/input_files/'
	OUT = '/data/user_storage/output_files/'

	#Load in Labware
	tiprack_20_1 = protocol.load_labware('opentrons_96_tiprack_20ul', '3')
	tiprack_20_2 = protocol.load_labware('opentrons_96_tiprack_20ul', '6')
	tiprack_20_3 = protocol.load_labware('opentrons_96_tiprack_20ul', '9')
	p20_tipracks = [tiprack_20_1, tiprack_20_2, tiprack_20_3]

	tiprack_300_1 = protocol.load_labware('opentrons_96_tiprack_300ul', '10')
	p300_tipracks = [tiprack_300_1]

	barcode_elution_plate = protocol.load_labware('starlab_96_wellplate_200ul_pcr', '5')
	final_dilute_plate = protocol.load_labware('nest_96_wellplate_100ul_pcr_full_skirt', '2')
	reservoir = protocol.load_labware('nest_12_reservoir_15ml', '1') 
	NFW = reservoir.wells_by_name()['A12']
	low_bind_tubes = protocol.load_labware('eppendorf_lo_bind_24_tuberack_1500ul', '4')

	tube_1 = low_bind_tubes.wells_by_name()['A1']
	FC_tube_1 = low_bind_tubes.wells_by_name()['A2']
	tube_2 = low_bind_tubes.wells_by_name()['C1']
	FC_tube_2 = low_bind_tubes.wells_by_name()['C2']

	# Load in Pipettes
	p20 = protocol.load_instrument(P20, P20_MOUNT, tip_racks= p20_tipracks)
	p300 = protocol.load_instrument(P300, P300_MOUNT, tip_racks= p300_tipracks)

	# Read in CSV file with sample volumes needed for each well. 
	# Load this onto OT-2 robot using SSH before simulating protocol
	# through Opentrons App. 
	library_concentrations = pd.read_csv(
		f'{IN}{DATE}_Library_Elution_Concentrations.csv', header = None)
	df = pd.DataFrame(library_concentrations)

	# Split the s into 2 separate flow cells 
	flow_cell_1 = df.iloc[:NO_SAMPLES,:]
	flow_cell_2 = df.iloc[NO_SAMPLES:,:]

	#Get ready to start protocol 
	protocol.set_rail_lights(True) #Turn on lights
	start_time = time.time()
	
    #If the protocol is not simulating take a video of the protocol
	if not protocol.is_simulating() & VIDEO == True:
		video.take_video(VIDEO_LENGTH, PROTOCOL_NAME, USER, DATE, start_time)

	#Protocol Commands
	def prepare_flow_cell(fc_number, flow_cell_df, prep_tube, lib_tube):
		# To prepare a Nanopore barcoded library for sequencing the final 
		# library tube must contain 1 mg of DNA in a final volume of 47ul. 
		# Samples below 2 ng/ul will not be included in this calculation. 
		# If there is some volume spare the library will be topped up with
		# these samples instead of nuclease free water. 
		samples_fc = len(flow_cell_df[(flow_cell_df[2] >= 2)]) 
		protocol.comment(
			f'''Number of Samples to Include in Flow Cell {fc_number}: 
				{samples_fc}''')

		below_fc = flow_cell_df[(flow_cell_df[2] < 2)]
		below_2_fc = len(below_fc)
		protocol.comment(
			f'''Number of Samples below 2 ng/ul in Flow Cell {fc_number}:
			 {below_2_fc}''')

		#Work out ng/ul needed of each sample in final library tube 
		sample_conc = round(((1 / samples_fc) * 1000), 2) 
		protocol.comment(
			f'''{sample_conc} ng/ul Needed of Each Sample in Flow Cell 
				{fc_number}''')
		
		flow_cell_nd = flow_cell_df.loc[(flow_cell_df[2] >= 2) &
			(flow_cell_df[2] <= sample_conc)]
		nd_samples = len(flow_cell_nd)
		protocol.comment(
			f'''{nd_samples} Samples to be Added to Final Libray Tube 
				without Dilution for Flow Cell {fc_number}''')

		# To prevent pipetting samples into final library tube below 1ul, we 
		# will dilute them to 2x the final concentration, transfer 1ul of this
		# to a preparation tube then transfer half of this to the final library
		# tube. Therefore the final library will contain 1x concentration of 
		# these samples.
		flow_cell_2x_sample_conc = sample_conc * 2 
		protocol.comment(
			f'''{flow_cell_2x_sample_conc} ng/ul Needed of Each Sample in 2x 
				Tube for Flow Cell {fc_number}''')

		# There is a threshold concentration needed to be able to dilute 
		# samples to 2x concentration in 10ul final volume with only 10ul of 
		# sample available. Samples above the sample concentration required for
		# final library and below the threshold needed to dilute to 2x final 
		# concentration will be diluted to 1x concentration in 5 ul. Then 1 ul 
		# of each of these samples will be added to the final library tube. 
		max_conc = ((flow_cell_2x_sample_conc*10)/10) 
		flow_cell_5ul = flow_cell_df.loc[(flow_cell_df[2] >= sample_conc) & 
			(flow_cell_df[2] < max_conc)]
		dil_5ul = len(flow_cell_5ul)
		
		#ul to transfer into final library tube
		fc_vol = (samples_fc - nd_samples - dil_5ul)  / 2 
		protocol.comment(
			f'''{fc_vol} ul of 2x Samples Needed to Transfer into Library Tube
			 for Flow Cell {fc_number}''')

		#Calculate volume of each sample needed for dilution to 2x final conc
		flow_cell_2x_10ul = flow_cell_df.loc[flow_cell_df[2] >= max_conc]
		flow_cell_2x_10ul[3] = round(
			(flow_cell_2x_sample_conc*10)/flow_cell_2x_10ul[2], 1)
		flow_cell_2x_10ul[4] = 'diluted in 10ul'
		flow_cell_2x_10ul[5] = sample_conc

		well_ids = flow_cell_2x_10ul[0].tolist()
		sample_vols = flow_cell_2x_10ul[3].tolist()

		protocol.comment(
			f'''Transfer NFW for Samples to be Diluted to 2x Concentration for
			 Flow Cell {fc_number}''') 
		p20.pick_up_tip()
		for well, volume in zip(well_ids, sample_vols): 
			p20.transfer(
				(10-volume), 
				NFW, 
				final_dilute_plate.wells_by_name()[well], 
				new_tip = 'never')
		p20.drop_tip()
		
		protocol.comment(
			f'''Add volume of each Sample to dilute to 
				{flow_cell_2x_sample_conc} ng/ul for Flow Cell {fc_number}''')
		for well, volume in zip(well_ids, sample_vols):
			p20.pick_up_tip()
			p20.transfer(
				volume,
				barcode_elution_plate.wells_by_name()[well].bottom(0.5),
				final_dilute_plate.wells_by_name()[well],
				new_tip = 'never')
			p20.mix(5, 8, final_dilute_plate.wells_by_name()[well].bottom(0.5),
				rate = 3)
			p20.drop_tip()

		protocol.comment(
			f'''Add 1ul of each of the 2x diluted samples to Flow Cell 
				{fc_number} 2x Prep Tube''')
		for well in well_ids:
			p20.transfer(1, final_dilute_plate.wells_by_name()[well], 
				prep_tube.bottom(0.2), new_tip = 'always')
	
		protocol.pause(f'Remove Flow Cell {fc_number} 2x Tube and Vortex')

		protocol.comment(
			f'''Transfer {fc_vol}ul of 2x Samples to Flow Cell {fc_number} 
				Final Library Tube''')
		if fc_vol > 20: 
			p300.transfer(fc_vol, prep_tube, lib_tube)
		else:
			p20.transfer(fc_vol, prep_tube, lib_tube)


		protocol.comment(
			f'''Calculate Volumes Needed to Dilute Samples to 1x Sample 
				Concentration in 5ul''')
		flow_cell_5ul[3] = round((sample_conc*5)/flow_cell_5ul[2], 1)
		flow_cell_5ul[4] = 'diluted in 5ul'
		flow_cell_5ul[5] = sample_conc
	
		well_ids = flow_cell_5ul[0].tolist()
		sample_vols = flow_cell_5ul[3].tolist()


		protocol.comment(
			f'''Transfer NFW for Samples to be Diluted to 1x Concentration for
				Flow Cell {fc_number}''') 
		p20.pick_up_tip()
		for well, volume in zip(well_ids, sample_vols): 
			p20.transfer(
				(5-volume), 
				NFW, 
				final_dilute_plate.wells_by_name()[well], 
				new_tip = 'never')
		p20.drop_tip()

		
		protocol.comment(
			f'Dilure sample for Flow Cell {fc_number} to {sample_conc} ng/ul')
		for well, volume in zip(well_ids, sample_vols):
			p20.pick_up_tip()
			p20.transfer(
				volume,
				barcode_elution_plate.wells_by_name()[well],
				final_dilute_plate.wells_by_name()[well],
				new_tip = 'never')
			p20.mix(5, 3, final_dilute_plate.wells_by_name()[well].bottom(0.5),
				rate = 3)
			p20.drop_tip()


		protocol.comment(
			f'''Add 1ul of each of 1x diluted samples to Flow Cell {fc_number} 
				Final Library Tube''')

		for well in well_ids:
			p20.transfer(1, final_dilute_plate.wells_by_name()[well].bottom(), 
					lib_tube.bottom(0.2), new_tip = 'always')


		#Samples that can be added neat to final flow cell tube 
		flow_cell_nd[3] = round(sample_conc/flow_cell_nd[2], 1) 
		flow_cell_nd[4] = 'neat'
		flow_cell_nd[5] = round(flow_cell_nd[2] * flow_cell_nd[3], 2)
		total_neat = round(flow_cell_nd[3].sum(), 2)
		protocol.comment(
			f'''{total_neat}ul of Samples being Added Neat to Final Library 
				for Flow Cell {fc_number}''')
	
		nd_well_ids = flow_cell_nd[0].tolist()
		nd_sample_vols = flow_cell_nd[3].tolist()
	
		for well, volume in zip(nd_well_ids, nd_sample_vols):
			p20.transfer(volume, barcode_elution_plate.wells_by_name()[well], 
			lib_tube, mix_after = (5, 10),  new_tip = 'always')

		#Work out how much volume left to top up to 47ul
		total_vol = 47
		left_vol = round(total_vol - fc_vol - total_neat - dil_5ul, 2)
		protocol.comment(f'Volume left to top up: {left_vol}')

		if samples_fc < 48:
			add_vol = left_vol / below_2_fc
			if add_vol > 8:
				add_vol = 8
				nfw_vol = left_vol - add_vol
			below_fc[3] = add_vol
			below_fc[4] = 'neat'
			below_fc[5] = (below_fc[2] * below_fc[3])
			protocol.comment(f'Add {add_vol}ul of each sample below 2ng/ul')
			for sample in below_fc[0].tolist():
				p20.transfer(add_vol, 
					barcode_elution_plate.wells_by_name()[sample], lib_tube)
			if add_vol == 8:
				p20.transfer(nfw_vol, NFW, lib_tube)
			dfs = [flow_cell_2x_10ul, flow_cell_5ul, flow_cell_nd, below_fc]
			output_file = pd.concat(dfs)
			output_file = output_file.sort_index()
			output_file.columns = [
				'Well ID', 'Sample ID', 'Elution Concentration ng/ul', 
				'Volume to Add', 'Preparation for Library', 
				'Concentration in Library']

		else:
			nfw_vol = left_vol
			protocol.comment(f'''Add {nfw_vol}ul of NFW to Top Up Flow Cell 
				{fc_number} to 47ul''')
			if nfw_vol > 20:
				p300.transfer(nfw_vol, NFW, lib_tube, mix_after = (10, 30))
			else:
				p20.transfer(nfw_vol, NFW, lib_tube, mix_after = (10, 20))
			dfs = [flow_cell_2x_10ul, flow_cell_5ul, flow_cell_nd]
			output_file = pd.concat(dfs)
			output_file = output_file.sort_index()
			output_file.columns = [
				'Well ID', 'Sample ID', 'Elution Concentration ng/ul', 
				'Volume to Add', 'Preparation for Library', 
				'Concentration in Library']

		file_path = f'''
			{OUT}{DATE}_Flow_Cell_{fc_number}_Library_Preparation_Calculations.csv'''
		output_file.to_csv(file_path, header = True, index = False)

	prepare_flow_cell(1, flow_cell_1, tube_1, FC_tube_1)
	prepare_flow_cell(2, flow_cell_2, tube_2, FC_tube_2)

	protocol.comment('Close lids, Pipeline complete! Time to sequence!')

	#Turn lights off 
	protocol.set_rail_lights(False)

	#Write details of OT-2 use to log file 
	if not protocol.is_simulating() & WRITE_TO_LOG == True:
		log.log_record(start_time, P300_MOUNT, P300, P20, DATE,
         USER, PROTOCOL_NAME, VIDEO, MODULES)
```
