## Supplementary Data 1 for "Characterization of recombinase-based genetic parts and circuits using nanopore sequencing": README.md

# Automated Memory Readout Pipeline using the OT-2

## Set Up

### Software & API:
App & Robot Software tested to 5.0.2<br>
API Version: 2.11

### OT-2(s)
This pipeline has been run with one OT-2 over two extended days where the pipettes and deck was reconfigured after each protocol. When running for the first time in this way please allow at least 3 days including the Nanopore sequencing set up.

The best set up we achieved was using two OT-2s - one with both mult-channels installed for use with all three modules interchangeably (Thermocycler, Magnetic Module, Temperature Module) and one with the single channels for dilutions and reagent preparation steps. This meant only one pipette change and that some protocols could be run in parallel / set up in advance to speed up the pipeline. With this set up, the pipeline can be run comfortably in two working days including Nanopore sequencing preparation.

**Before each protocol was started a calibration step was taken to check calibration of all labware used and apply any offsets if needed.**

### SSH Connection Set Up: Transferring Files to and from the OT-2
During the pipeline there will be input files required by certain protocols and output files generated by them. Prior to running the pipeline please ensure you have [SSH access](https://support.opentrons.com/en/articles/3203681-setting-up-ssh-access-to-your-ot-2) set up to the OT-2 and can use [SCP](https://support.opentrons.com/en/articles/3690659-copying-files-to-and-from-your-ot-2-with-scp) to transfer files over command line from and to your computer to the OT-2.

To view the directories and files on the OT-2 you can [connect](https://support.opentrons.com/en/articles/3287453-connecting-to-your-ot-2-with-ssh) to your OT-2 using the SSH connection.<br>
Once connected set up two directories in the user storage directory for the input and output files to be saved to using the terminal:

```
cd /
cd /data/user_storage
mkdir input_files
mkdir output_files
```

### Labware & Reagents:
Prior to running any protocols ensure you have [set up your custom labware definitions folder](../labware/README.md) in the Opentrons App.
<br>
<br>

## Tailing PCR Reaction
### Protocol 1: Preparing the Tailing PCR Mastermix
This protocol creates a mastermix for the tailing PCR in a reservoir to allow quick transfer to the 96-well plate in the next protocol.

#### [Protocol 1](1_Prep_Tailing_PCR_Mastermix.py) - Creates one mastermix for 96 samples amplifying memory registers.<br>
#### [Protocol 1a](1a_Prep_Tailing_Cascade_Mastermixes.py) - Creates two mastermixes with two sets of primers.

**Protocol Length:** 7 - 10 minutes<br>
**Pipettes Needed:** P300 Single Gen2, P1000 Single Gen2<br>
![Deck Layout](deck_layouts/Deck_Layout_1_%26_1a.png)

### Protocol 2: Lysis of Cells & Tailing PCR
The temperature block is used to lyse sample cultures at 95°C before they are spun down (external centrifuge 4,000rpm for 2 minutes) and supernatant transferred to a storage plate. The mastermix and each sample is added to the on deck thermocycler and used to complete tailing PCR reaction.

#### [Protocol 2](2_PCR_Tailing_Reaction.py) - Amplifying memory registers in 96 samples.<br>
#### [Protocol 2a](2a_PCR_Tailing_Reaction_Cascade.py) - Amplifying the memory register and integrase plasmid for each sample.

**Protocol Length:** 3 hours 32 minutes (Protocol 2), 5 hours 20 minutes (Protocol 2a)<br>
**Pipettes Needed:** P20 Multi Gen2, P300 Multi Gen2<br>
**Modules Needed:** Thermocycler, Temperature Module Gen2<br>
![Deck Layout](deck_layouts/Deck_Layout_2_&_2a.png)

## DNA Purification
### Protocol 3: DNA Purification of Tailing PCR Products
The magnetic module is used to purify the tailing PCR reactions with 0.4x KAPA magnetic beads to select for the larger DNA fragments. These are eluted into 13 uL of nuclease free water.

#### [Protocol 3](3_Pur_DNA_Mag_20ul.py)<br>

**Protocol Length:** 1 hour 51 minutes<br>
**Pipettes Needed:** P20 Multi Gen2, P300 Multi Gen 2<br>
**Modules Needed:** Magnetic Module Gen2<br>
![Deck Layout](deck_layouts/Deck_Layout_3_&_10.png)

## DNA Quantification
DNA is prepared for quantification using the [Quantifluor ONE dsDNA system](https://www.promega.co.uk/products/rna-analysis/dna-and-rna-quantitation/quantifluor-one-dsdna-system/?catNum=E4870) kit across two black, clear bottomed 96 well plates. These are then read using a fluorescent plate reader from the bottom (Excitation: 485 nm, Emission: 535 nm) and analysed in the available [Excel workbook](https://www.promega.co.uk/resources/tools/quantifluor-dye-systems-data-analysis-workbook/).

### Protocol 4: Prepare Standards
Prepare the standards by creating a dilution series of the Lambda dsDNA standard in 1X TE Buffer (provided in the kit). This protocol makes enough for the analysis to be run twice and so the standards may kept overnight and used in the second DNA quantification step in the pipeline.

#### [Protocol 4](4_Prep_Quantifluor_Stds.py)<br>

**Protocol Length:** 7 minutes<br>
**Pipettes Needed:** P20 Single Gen2<br>
![Deck Layout](deck_layouts/Deck_Layout_4.png)

### Protocol 5: Add Standards and Samples to Quantifluor Dye
Adds 200 uL of Quantifluor dye across one full plate and 4 columns in a second plate. In the first plate Columns 1 and 12 will contain the standards in duplicate and samples from the elution plate will be from and in Columns 2 to 11. The second plate will also contain the standards in duplicate in Columns 1 and 4. Samples from Column 1 in the sample plate will be read in Column 2 of the reader plate and Column 12 read in Column 3.

#### [Protocol 5](5_Prep_Quantifluor_Samples.py)<br>

**Protocol Length:** 18 minutes<br>
**Pipettes Needed:** P20 Multi Gen2, P300 Multi Gen2<br>
![Deck Layout](deck_layouts/Deck_Layout_5.png)

## Barcoding PCR
Samples are diluted to 1.18 ng/uL in nuclease free water and added to a PCR reaction using NEB's [LongAmp Taq 2X Mastermix](https://international.neb.com/products/m0287-longamp-taq-2x-master-mix#Product%20Information) and [Oxford Nanopore's PCR Barcoding Expansion Kit](https://store.nanoporetech.com/uk/pcr-barcoding-expansion-1-96.html).

### Protocol 6: Calculate Dilution Steps for Each Sample based on DNA Quantification
A simple CSV (example [here](/ot2_pipeline_files/2022-03-01_Elution_Concentrations.csv)) with the well IDs and DNA concentrations is uploaded the OT-2 using the terminal via USB connection: <br>
***Note: All dates are required in YYYY-MM-DD format when `DATE` is used***
```
scp -i ot2_ssh_key FILE_PATH_ON_YOUR_COMPUTER/DATE_Elution_Concentrations.csv root@ROBOT_IP:/data/user_storage/input_files/
```
The protocol calculates the concentrations and volumes needed of each sample to dilute them to 1.18 ng/uL after a 1 in 7 dilution. These calculations are written to CSV files saved on the OT-2 for when protocol 8 is run. Any samples with concentrations below 20 ng/uL are too low to be diluted 1 in 7 and so during this protocol are removed from the elution plate into another plate to be diluted individually. <br>
A record of all dilutions and calculations can be obtained from OT-2 using:
```
scp -i ot2_ssh_key -r root@ROBOT_IP:/data/user_storage/output_files/DATE_Barcode_PCR_Preparation_Calculations.csv /FILE_PATH_ON_YOUR_COMPUTER/
```

#### [Protocol 6](6_Prep_Barcode_Dilution_Pt_1.py)<br>

**Protocol Length:** 5 minutes + <br>
**Pipettes Needed:** P20 Single Gen2<br>
![Deck Layout](deck_layouts/Deck_Layout_6.png)

### Protocol 7: Dilute All Samples 1 in 7
8 uL of each elution is added to 48 uL of nuclease free water and thoroughly mixed.

#### [Protocol 7](7_Prep_Barcode_Dilution_Pt_2.py)<br>

**Protocol Length:** 15 minutes <br>
**Pipettes Needed:** P20 Multi Gen2, P300 Multi Gen2<br>
![Deck Layout](deck_layouts/Deck_Layout_7.png)

### Protocol 8:
Dilution of all samples to 1.18 ng/uL.

#### [Protocol 8](8_Prep_Barcode_Dilution_Pt_3.py)<br>

**Protocol Length:** 1 hour 32 minutes <br>
**Pipettes Needed:** P20 Single Gen2, P300 Single Gen2<br>
![Deck Layout](deck_layouts/Deck_Layout_8.png)

### Protocol 9:
Barcoding PCR set up and execution in 45 uL reaction. Set `CASCADE_EXP` to `True` in the script if sequencing integrase plasmids as well as memory registers.

#### [Protocol 9](9_PCR_Barcoding_Reaction.py)<br>

**Protocol Length:** 1 hour 45 minutes (2 hours 32 minutes if amplifying integrase plasmids) <br>
**Pipettes Needed:** P20 Multi Gen2, P300 Multi Gen2<br>
**Modules Needed:** Thermocycler<br>
![Deck Layout](deck_layouts/Deck_Layout_9.png)

## DNA Purification & Quantification
Purfify the DNA using Protocol 10 then repeat Protocol 5 to quantify the eluted DNA.

### Protocol 10:
The magnetic module is used to purify the barcoding PCR reactions with 0.4x KAPA magnetic beads to select for the larger DNA fragments. These are eluted into 13 uL of nuclease free water.

#### [Protocol 10](10_Pur_DNA_Mag_40ul.py)<br>

**Protocol Length:** 2 hours 5 minutes <br>
**Pipettes Needed:** P20 Multi Gen2, P300 Multi Gen2<br>
**Modules Needed:** Magnetic Module Gen2<br>
![Deck Layout](deck_layouts/Deck_Layout_3_&_10.png)

## Library Preparation for Nanopore Sequencing
This protocol prepares two libraries for Nanopore sequencing each containing 48 samples. Each library is required to be 47 uL and contain 1 mg of DNA for sequencing protocol.

### Protocol 11:
The required concentration of each sample is determined by dividing the number of samples above 2 ng/uL by 1 mg DNA. Depending on the concentration of the eluted the DNA, each sample is either first diluted to 2x the concentration needed in the final library or 1x. Those diluted to 2x are all combined in one tube, vortexed and half of this mixture is added to the final library tube. This ensures that the required DNA concentration is met without pipetting below 1 uL which is outside of the range of the P20 pipette. Those samples diluted to 1x are added directly to final library tube in a 1 ul transfer. The library tube is then topped up to 47 uL with any samples below 2 ng/uL and or nuclease free water.

A simple CSV with the well IDs, sample IDs and DNA concentrations is uploaded the OT-2 using the terminal via USB connection: <br>
```
scp -i ot2_ssh_key FILE_PATH_ON_YOUR_COMPUTER/DATE_Library_Elution_Concentrations.csv root@ROBOT_IP:/data/user_storage/input_files/
```
Records of all dilutions and concentrations of DNA in each flow cell can be obtained from OT-2 using:
```
scp -i ot2_ssh_key -r root@ROBOT_IP:/data/user_storage/output_files/DATE_Flow_Cell* /FILE_PATH_ON_YOUR_COMPUTER
```

#### [Protocol 11](11_Prep_DNA_Libraries.py)<br>

**Protocol Length:** 1 hour 45 minutes <br>
**Pipettes Needed:** P20 Single Gen2, P300 Single Gen2<br>
![Deck Layout](deck_layouts/Deck_Layout_11.png)
