## Supplementary figures and images for "Characterization of recombinase-based genetic parts and circuits using nanopore sequencing"

### Deck_Layout_1.png

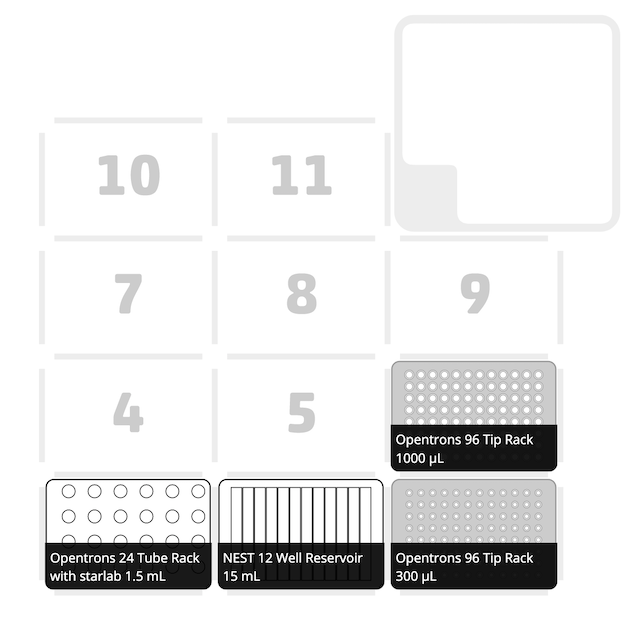

### Deck_Layout_2.png

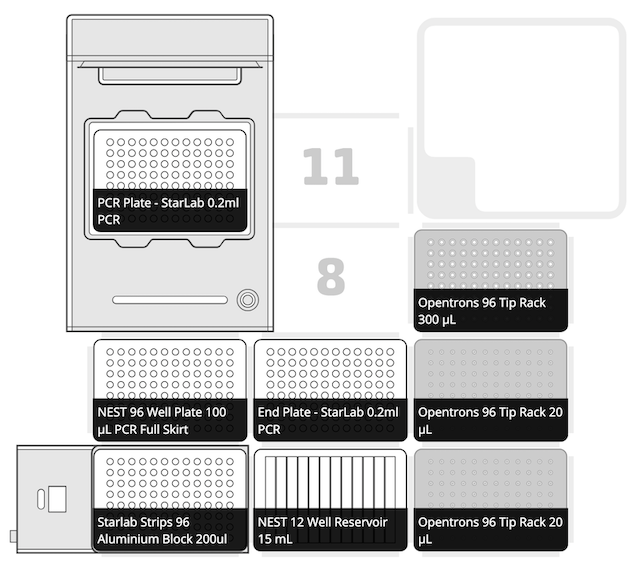

### Deck_Layout_3_&_10.png

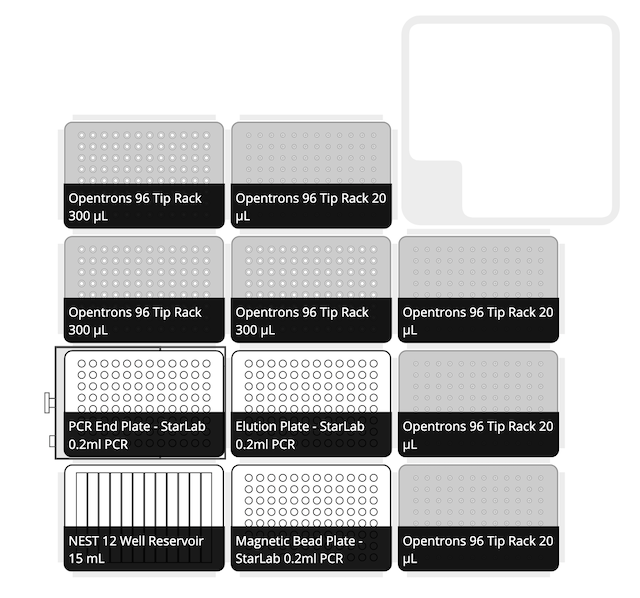

### Deck_Layout_4.png

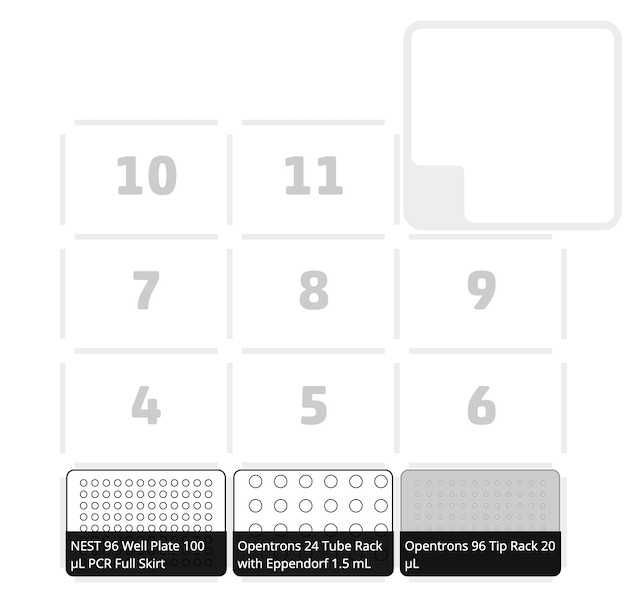

### Deck_Layout_5.png

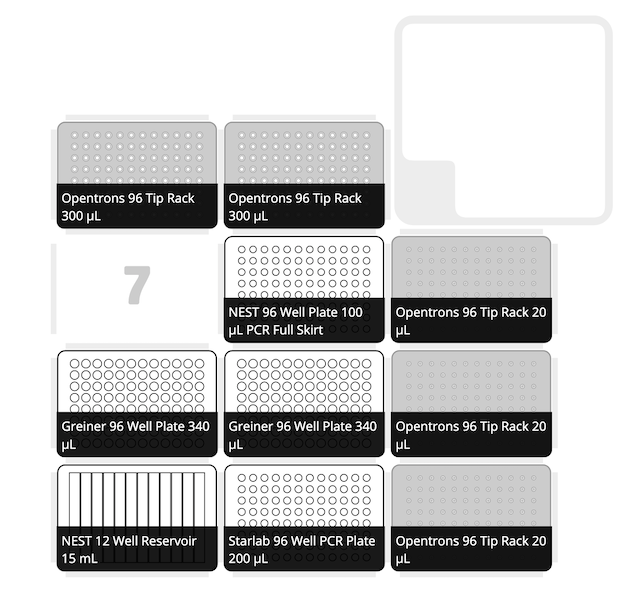

### Deck_Layout_6.png

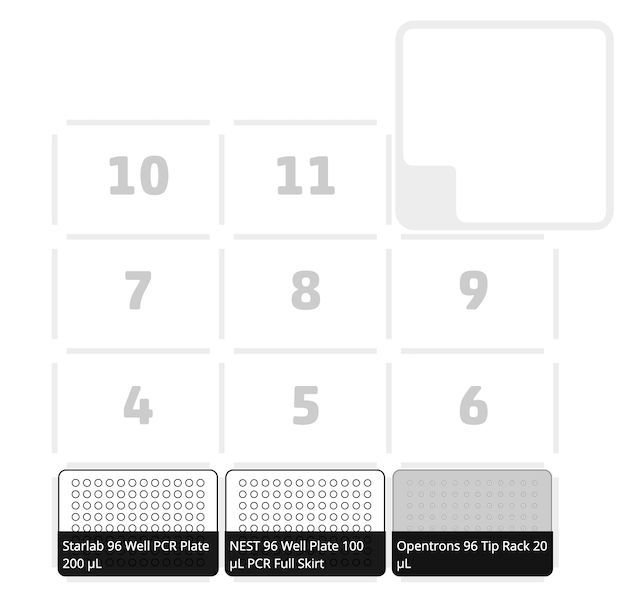

### Deck_Layout_7.png

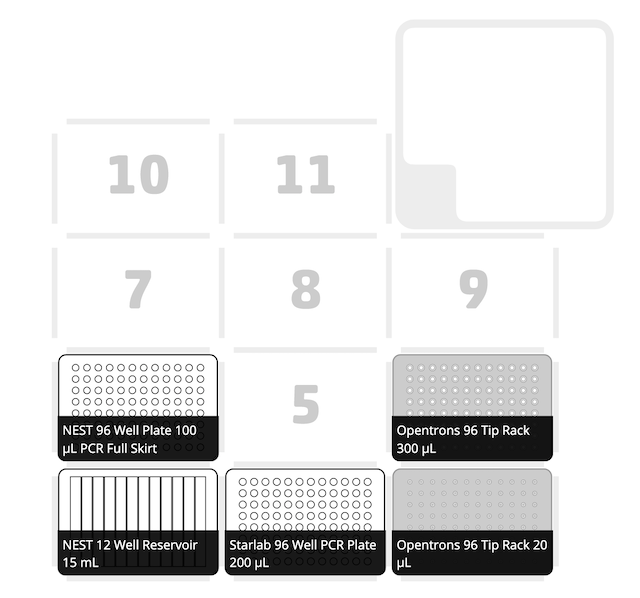

### Deck_Layout_8.png

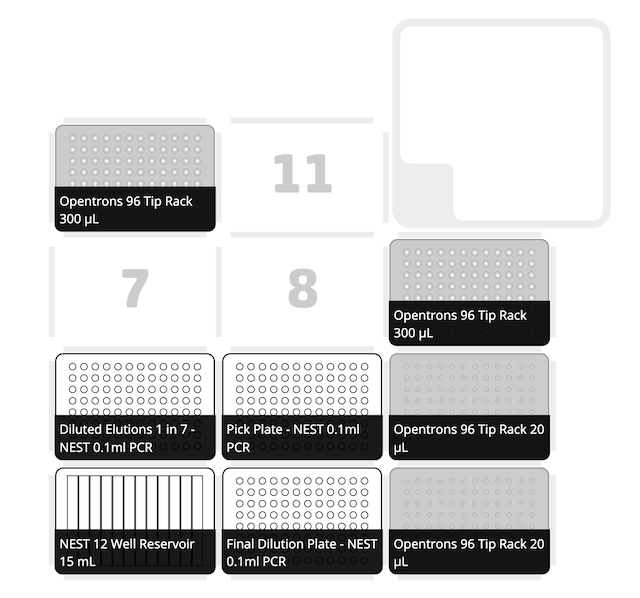

### Deck_Layout_9.png

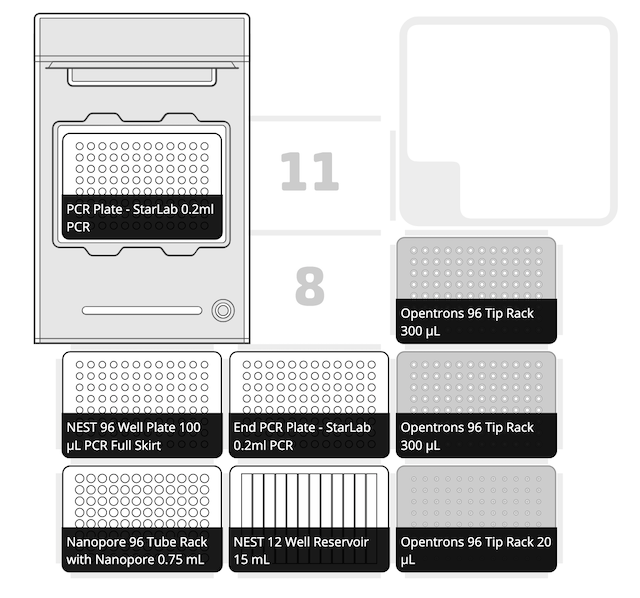

### Deck_Layout_11.png

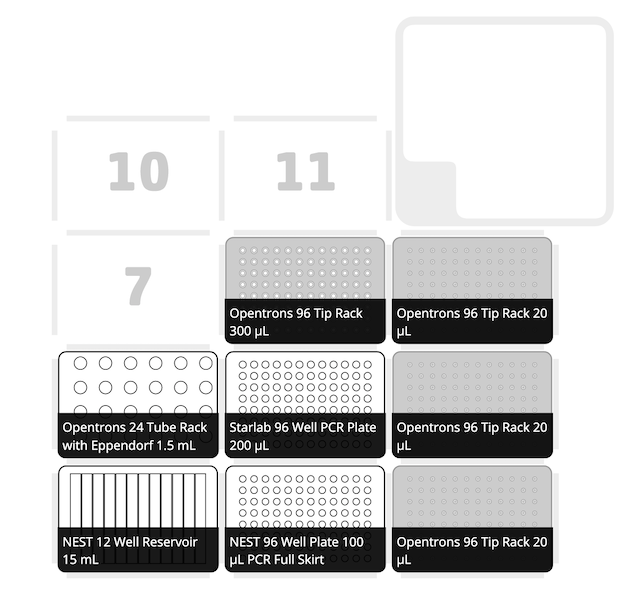
